# Cognitive control and automatic interference in mind and brain: A unified model of saccadic inhibition and countermanding

**DOI:** 10.1101/535104

**Authors:** Aline Bompas, Anne Eileen Campbell, Petroc Sumner

## Abstract

Countermanding behavior has long been seen as a cornerstone of executive control – the human ability to selectively inhibit undesirable responses and change plans. However, scattered evidence implies that stopping behavior is entangled with simpler automatic stimulus-response mechanisms. Here we operationalize this idea by merging the latest conceptualization of saccadic countermanding with a neural network model of visuo-oculomotor behavior that integrates bottom-up and top-down drives. This model accounts for all fundamental qualitative and quantitative features of saccadic countermanding, including neuronal activity. Importantly, it does so by using the same architecture and parameters as basic visually guided behavior and automatic stimulus-driven interference. Using simulations and new data, we compare the temporal dynamics of saccade countermanding with that of saccadic inhibition (SI), a hallmark effect thought to reflect automatic competition within saccade planning areas. We demonstrate how SI accounts for a large proportion of the saccade countermanding process when using visual signals. We conclude that top-down inhibition acts later, piggy-backing on the quicker automatic inhibition. This conceptualization fully accounts for the known effects of signal features and response modalities traditionally used across the countermanding literature. Moreover, it casts different light on the concept of top-down inhibition, its timing and neural underpinning, as well as the interpretation of stop-signal reaction time, the main behavioral measure in the countermanding literature.

## 1. Introduction

There is a long tradition in psychology and neuroscience of drawing a conceptual distinction between ‘top-down’ volitional processes and ‘bottom-up’ automatic responses. However, this does not mean there is a clear distinction in the brain. Nor is it likely that any behavior produced by any elaborate animal is entirely bottom-up or top-down in nature. Rather, one can envisage an enmeshed relationship whereby increasingly selective or “voluntary” systems have grown out of, and remain entwined with, phylogenetically older automatic mechanisms (see Harrison, Freeman, & Sumner, 2014; McBride, Boy, Husain, & Sumner, 2012; Sumner & Husain, 2008; Verbruggen, Best, Bowditch, Stevens, & McLaren, 2014; Verbruggen & Logan, 2008; Wessel & Aron, 2017). Conceptually, several fields are moving away from the idea of an “executive controller”, and working toward characterizing the “army of idiots” that allow successful action control (Monsell & Driver, 2000; Verbruggen, McLaren, & Chambers, 2014).

Here we address a long-standing topic in top-down control: the ability to *withhold* action. Just as music is about the spaces as well as the notes, behavior is about the actions we don’t make as well as the actions we do make (Logan, Yamaguchi, Schall, & Palmeri, 2015; Noorani & Carpenter, 2017; Schall, Palmeri, & Logan, 2017). Clearly, humans are able to control their motor systems and refrain from always acting reflexively, habitually or impulsively. We have the flexibility to halt and change action plans in rapidly changing situations, such as sport, social interactions, or driving a car. The precise mechanisms that might enable us to do this have been a major focus of psychology and cognitive neuroscience. Although stopping behavior has always been broadly conceptualized as top-down control, a range of stimulus-driven or habitual influences were envisaged early on (Hanes & Schall, 1995; Logan & Cowan, 1984; Schall & Thompson, 1999), before being further discussed and demonstrated (Schmidt & Berke, 2017; Verbruggen, Best, et al., 2014; Verbruggen, McLaren, et al., 2014; Wessel & Aron, 2017).

Here we build on this nuanced background and make three further contributions. Focusing on the ability to withhold eye-movements, we identify the critical first phase of stopping with a known but previously unconnected automatic interference mechanism. Second, we argue that the ability to withhold action can be best understood through models that clearly delineate two types of signal with different origins and dynamics, the first being a transient automatic drive triggered by any change in the visual environment. These automatic signals interfere with ongoing action plans, temporarily delaying their execution, buying time for slower and more selective drives to cancel or change the plan. Third, understanding the neural underpinnings of decision then shifts from mainly focusing on *move* neurons to including *sensorimotor* neurons, given that the successful model is an implementation of the latter.

Animal brains are full of inhibitory connections (see Noorani & Carpenter, 2017 for a review), many of which can be considered very basic and automatic properties of neural maps or local networks. We believe these low-level mechanisms critically shape behaviors traditionally ascribed to top-down control and, in some conditions, even form the main basis for well-known hallmarks of ‘control’ behavior. Even though they may be rather indiscriminate and simple, the potential advantage of stimulus-driven inhibitory circuits would be their speed - a quick interruption allowing slower more complex processes time to update action plans (e.g. Schmidt & Berke, 2017). If we can understand how automatic, rapid and indiscriminate mechanisms work within tasks associated with top-down control, it should help us unify literatures on control and distraction (e.g. Wessel & Aron, 2017) and also better integrate the functional consequences of basic sensorimotor processes with concepts of higher cognitive functions.

Computational models are important tools to develop and test our understanding of these mechanisms. In recent years, their number and complexity have increased, with models becoming more biologically grounded, attempting to capture not only behavioral data, but also neuronal recordings (Bompas, Hedge, & Sumner, 2017; Bompas & Sumner, 2011; Boucher, Palmeri, Logan, & Schall, 2007; Cutsuridis, Smyrnis, Evdokmds, & Perantonis, 2007; Kopecz, 1995; Lo, Boucher, Pare, Schall, & Wang, 2009; Logan et al., 2015; Meeter, Van der Stigchel, & Theeuwes, 2010; Purcell et al., 2010; Ramakrishnan, Sureshbabu, & Murthy, 2012; Shadlen, Britten, Newsome, & Movshon, 1996; Trappenberg, Dorris, Munoz, & Klein, 2001; Wiecki & Frank, 2013). However, the focus on different tasks, animal models and anatomical subsystems has led to partly segregated subfields in the literature, and sometimes to the parallel development of distinct models attempting to capture different instantiations of similar cognitive functions. As a result, most current psychological models have been designed and constrained to capture mainly one task, and the generalizability to new tasks is not often tested. Although this limitation is inevitable in the early days of biologically inspired computational models of action decision, a desirable perspective for the field would be to move away from modeling tasks and start modeling the biological system trying to perform it. To achieve this, a first step is to draw modeling attempts together and develop more general models, ultimately able to predict human or animal behavior in new experimental conditions.

### Stopping

A prevalent paradigm of top-down inhibition used widely within the psychological, psychiatric and neurophysiological literatures is ‘countermanding’, epitomized by the stop signal task (Logan & Cowan, 1984; Noorani & Carpenter, 2017). Participants make simple responses to the presentation of a target and, on a minority of trials, are required to cancel (‘countermand’) their response following the onset of a stop-signal (Figure 1A). Hence, this task is designed to assess the volitional ability to rapidly inhibit responses that are already being planned.

**Figure 1.**
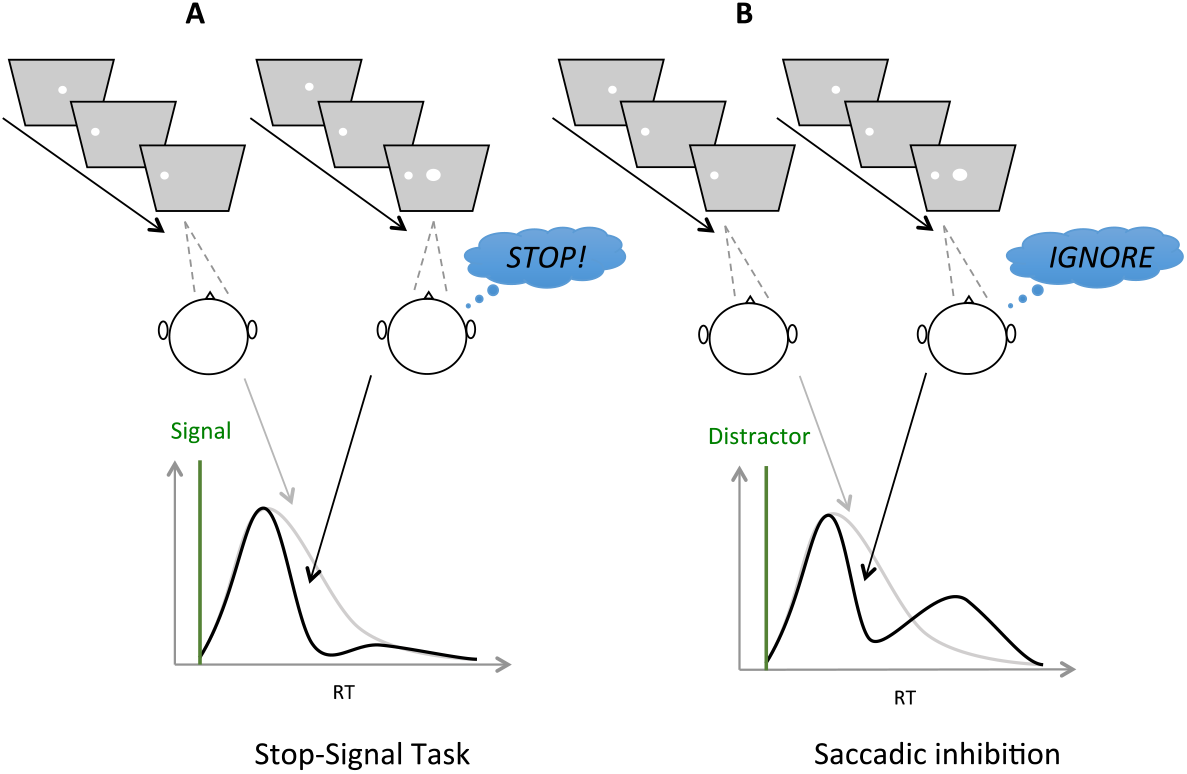
Typical design (above) and results (below) in the saccadic Stop-Signal Task (SST, panel A) and Saccadic Inhibition (SI, panel B) paradigms. Both paradigms involve a stimulus jump from center to periphery, sometimes followed by the onset of a central signal (right subpanels above, black lines below), sometimes not (left subpanels, grey lines). The signal onset time is indicated by vertical green lines and the delay between the target jump and the signal is referred to as the stimulus onset asynchrony (SOA). The two tasks differ in the instruction associated with the signal onset, withhold the saccade in the SST, ignore the signal and perform the saccade in the SI. **A**. Instructions to stop remove slower responses from the RT distribution, but fast responses escape (’failed stops’). **B.** The same visual events associated with an ignore instruction typically produce a dip in the latency distribution, where saccades are delayed and subsequently recover, so that the total number of saccades are about the same between signal present and no-signal distributions. We propose that on trials where participants are told to stop their saccade in response to the signal onset (A), the initial reduction in saccade probability has the same automatic source and therefore will happen at the same time as the dip in the ignore condition (B), but the recovery from the dip will be diminished or absent due to later top-down inhibition.

The process of such top-down inhibition has long been conceptualized as a race between competing “go” and “stop” mechanisms within the independent horse-race model (Logan & Cowan, 1984). If the countermand activity can overtake the go activity, then the response is not executed; whereas if the go activity reaches its threshold before the stop-response activity overtakes it, then the response is executed (known as a failed stop). Failed stops tend to have short latencies with respect to the stop signal, consistent with the idea that top-down inhibition did not have sufficient time to act.

Countermanding tasks have used a variety of response modalities and stimulus designs, but the basic principles of design and of behavioral outcomes are shared. The saccade (eye movement) countermanding task (Hanes & Schall, 1995) has been the dominant modality for monkey experiments, and has allowed the bridging of psychology and neurophysiology through the development of biologically inspired computational models. The conceptual race between go and stop processes was then mapped onto more complex models capturing the neural architecture of the saccadic control network (Boucher, Palmeri, et al., 2007; Lo et al., 2009; Ramakrishnan et al., 2012; Schall et al., 2017; Wong-Lin, Eckhoff, Holmes, & Cohen, 2010), implementing an antagonistic relationship between fixation and movement processes (Hanes, Patterson, & Schall, 1998; Munoz & Wurtz, 1993a, 1993b). This development allowed models to take a more nuanced approached to ‘top-down’ signals, wherein the stop signal becomes partly a visual drive to fixation neurons (Lo et al., 2009; Logan et al., 2015; Wong-Lin et al., 2010). Indeed, this dual effect captures pre-existing discussions that saccade countermanding using a central visual stop-signal might reflect a combination of automatic visual as well as top-down volitional inhibition, possibly due to stimulus-invoked activity of fixation cells of the superior colliculus (SC’, Cabel, Armstrong, Reingold, & Munoz, 2000; Morein-Zamir & Kingstone, 2006; Schall & Thompson, 1999).

At the same time, behavioral evidence has accumulated that low-level visual effects modulate most visuo-oculomotor behavior, even stopping behavior. After Hanes & Schall (1995) noted that the intensity of the stop-signal influences stopping ability, alterations to the stimuli were seen to affect the main outcome measure - the stop signal reaction time (SSRT) - across all types of countermanding tasks. For example a central visual signal provides a shorter SSRT than an auditory signal or a peripheral visual signal (Armstrong & Munoz, 2003; Asrress & Carpenter, 2001; Boucher, Stuphorn, Logan, Schall, & Palmeri, 2007; Cabel et al., 2000; Hanes & Carpenter, 1999; Hanes et al., 1998; Hanes & Schall, 1995; Ito, Stuphorn, Brown, & Schall, 2003; Morein-Zamir & Kingstone, 2006; Paré & Hanes, 2003; Stuphorn, Taylor, & Schall, 2000). In addition, introducing a 200 ms gap between fixation offset and target onset can reduce both reaction time and SSRT (Stevenson, Elsley, & Corneil, 2009).

Below we take a step further, proposing that the most characteristic part of rapid saccadic countermanding is initially *entirely* automatic, with slower endogenous signals built on top of rapid automatic disruption. We will argue that, in order to understand the interplay of volition and automaticity within a task or behavior, it is actually helpful to start with a model in which they are articulated separately as distinct inputs.

### Pausing and carrying on

In oculomotor behavior, new stimuli produce a hallmark phenomenon known as saccadic inhibition (SI, Bompas & Sumner, 2011; Buonocore & McIntosh, 2008, 2012; Buonocore, Purokayastha, & McIntosh, 2017; Edelman & Xu, 2009; McIntosh & Buonocore, 2014; Reingold & Stampe, 2002, 2004). SI was first discovered in the context of reading (hence the name, to distinguish it from latency effects due to word processing). It happens under most scenarios in which a flash or new stimulus occurs while the system is planning a saccade, whether when reading text, viewing a scene, in simple saccade experiments and even in optokinetic and infantile nystagmus (Harrison et al., 2014). When these irrelevant stimuli occur during saccade planning, a population of would-be saccades is temporarily withheld, creating a dip in the latency distribution time-locked to the onset of this distractor signal (Figure 1B). This inhibition is thought to be a purely automatic process where the distractor elicits competing activation in saccade planning areas (such as the Superior Colliculus) that limits the accumulating activity for the planned saccades (Bompas & Sumner, 2011; Edelman & Xu, 2009; Reingold & Stampe, 2002). The evidence that it is automatic comes from its ubiquitous appearance across all tasks and all participants tested, even when participants have explicit instructions to ignore new stimuli, and not doing so is detrimental to the task at hand.

SI is therefore identified by a characteristic latency distribution with three phases following the distractor signal (Figure 1B): the first 70-100 ms saccades entirely escape influence and are executed as usual (the distribution of saccades with or without signal exactly overlap); then there is a dip - a reduction in the number of saccades produced compared with baseline conditions (with no signal); lastly there is a recovery phase where the disrupted saccades are produced later in the distribution.

Volitional countermanding and automatic saccadic inhibition have so far been discussed in separate literatures and have different computational models associated with them. However, the only important difference between the two paradigms is the instruction associated with the signal: ignore in SI and stop in countermanding (Figure 2A). And indeed, we note that the first part of the latency distributions typical of both phenomena show a similar pattern: failed stops executed shortly after the signal escape inhibition and then, at some delay following the signal, there is a rapid reduction of response probability. In our hypothesis, this is the very same automatic dip as seen in SI. More selective control could then evolve later to inhibit the recovery phase, piggy-backing on the process begun by the automatic mechanism.

**Figure 2.**
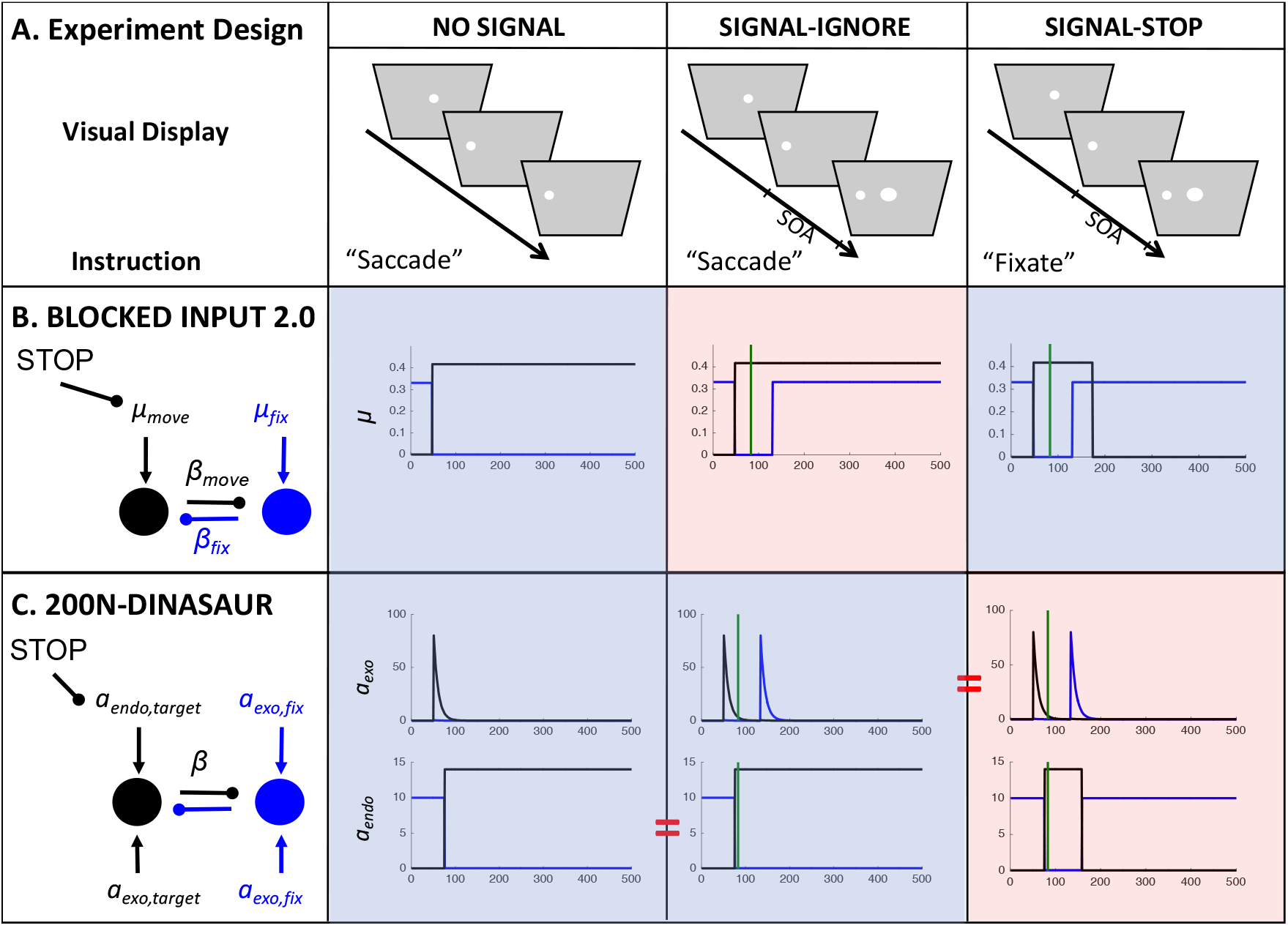
Inputs to Blocked Input 2.0 and 200N-Dinasaur for each task condition, based on published versions (blue shaded areas; Logan et al., 2015; Bompas & Sumner, 2011) or parsimonious generalizations to new conditions (red shaded areas, using SOA = 83 ms as in the new experiments introduced below). **A.** Schematic task conditions (see Figure 1 for description). **B.** Blocked Input 2.0 was originally designed for the stop task encompassing the NO-SIGNAL and SIGNAL-STOP conditions (blue shade). In the most parsimonious generalization to the IGNORE instructions (red shade), the late “blocking” of move input does not occur (black line), just as in NO-SIGNAL conditions, while the stimulus onset reactivates fixation input (blue line) just as in the SIGNAL-STOP condition. **C.** 200N-DINASAUR was shown to capture saccadic inhibition (NO-SIGNAL = pro-saccade, SIGNAL-IGNORE = distractor condition, blue shade). Out of the 200 nodes, here only the fixation and target nodes are shown. The model categorizes inputs as exogenous (stimulus-elicited and transient, upper plots) or endogenous (instruction-related and sustained, lower plots). A straightforward generalization to the STOP instruction (red shade) is to assume the exogenous inputs are unchanged, while the endogenous input switches from the target back to fixation, like in Blocked Input 2.0. Note that in Blocked Input 2.0, this switch is not simultaneous: fixation drive reappears before move drive is blocked to allow for the extra rapidity of a stimulus-driven response. In DINASAUR, the exogenous input already accounts for the rapid stimulus-elicited activity, so parsimoniously the endogenous switch can be simultaneous: the onset of endogenous fixation drive is given the same delay as the offset of endogenous saccade drive.

This kind of hypothesis has been proposed before, but never formally tested (Akerfelt, Colonius, & Diederich, 2006; E. Salinas & Stanford, 2018). It shares conceptual similarity with the Pause-then-Cancel theory (Schmidt & Berke, 2017) derived from studying basal ganglia in rats (although the specific concepts and implementations are different as explained further in Discussion). We also consider it belongs in a growing family of proposals attempting to integrate processes traditionally categorized as either volitional or reflexive. For example, Wessel and Aron (2017) propose that rapid stopping in humans entails the same fronto-basal-ganglia network that disrupts motor plans following unexpected events, potentially unifying literatures on countermanding with post-error slowing and attentional distraction in humans. Although Wessel and Aron’s theory is at the cognitive level, while ours is a mechanistic model of oculomotor planning, both carry the implication that countermanding is built on top of – and during evolution has grown out of – an indiscriminate response to novel visual stimuli. Likewise in the domain of motor priming, Sumner and Hussain (2008) argued that automatic inhibition is one of the building blocks for conscious voluntary planning and control, while others merged the concepts of reflex and volition in the concept of conditional automaticity (see Kunde, Kiesel, & Hoffmann, 2003 for a discussion). Back in the oculomotor domain, Harrison et al. (2014) proposed that voluntary saccade control shares mechanisms with, and probably emerged in evolution from the quick phases of stimulus-driven nystagmus.

### Contrasting and merging models

The computational models of countermanding and saccadic inhibition, while currently separate, are both biologically grounded and inspired by neuronal recordings. In fact, they share many properties. This being said, they also rely on fundamentally different assumptions. It therefore appears desirable to contrasts these models and use both paradigms to constrain a common model, able to capture both tasks. Below we outline how these differences may affect the ability for models to generalize across tasks.

The latest model for countermanding is the Blocked Input 2.0 model (Logan et al., 2015). In this model (Figure 2B), the visual onset of the “stop signal” is proposed to trigger two events: a quick return of sustained excitatory input to fixation node, followed by a blocking of the excitatory input to the movement node. While the second element is clearly described as top-down in nature, the short latency of the first event is strongly suggestive of a bottom-up nature. More generally, in all recent models of saccadic countermanding (Lo et al., 2009; Logan et al., 2015; Wong-Lin et al., 2010), fixation and movement neurons receive inputs tightly tied to the visual stimuli (targets and stop-signals), with onsets and offsets leading to step changes some 35 to 50 ms later. These changes typically precede inputs emanating from control neurons whose role is to cancel the action plan. However, we would argue that the early visually-driven signals conceptually merge two types of influences to decision: bottom-up and goal-driven inputs, as explained below.

Other models, previously developed to capture visual interferences in saccadic decision, more explicitly model the bottom-up signal as an automatic transient (i.e. they happen irrespective of the task and rapidly decay’, Bompas & Sumner, 2011; Kopecz, 1995; Kopecz & Schoner, 1995; Trappenberg et al., 2001). As such, they mimic signals typically observed in anaesthetized animal (Schmolesky et al., 1998) or in response to task-irrelevant distractors (Dorris, Olivier, & Munoz, 2007). In contrast, endogenous drives are captured as sustained inputs and depend on stimulus-response mapping. Therefore, in these models, a visual onset that is also task-relevant (such as the target onset) would trigger both a fast, transient input and a slower sustained input. The model nodes integrate these two pathways and therefore behave like visuo-movement neurons. Recordings in visuo-movement FEF and SC neurons of monkeys performing a visuo-oculomotor task under a speed or accuracy condition (Reppert, Servant, Heitz, & Schall, 2018) show that the delay and gain of the early visual response is unaffected by strategic adjustments (see also Heitz & Schall, 2012). Similarly, under this framework, whether the instruction is to ignore or stop to the signal, the same early interference would be expected, reflecting the effect of the unchanged automatic input associated with the signal onset.

In contrast, in all models of saccade countermanding, signals conceived as visually-driven change with onsets and offsets, like automatic signals, but are sustained for the whole duration of a stimulus, like goal-filtered signals in response to task-relevant stimuli. As a result, bottom-up and top-down inputs are tied into one stream and their modulations by visual events and instructions cannot be directly disentangled. This conceptualization is aligned with the assumption that decisions are most closely related to the activity within movement neurons, rather than within visuomovement neurons. Movement neurons in FEF and SC do delay the onset of their response on trials following stop-signals compared with trials following go-trials, consistent with strategic adjustments leading to behavioral slowing between these two conditions (Pouget et al., 2011). Similarly, under this conceptualization, the early adjustments in response to the signal could reflect task-related drives (or a mixture of automatic and task-related influences), and may therefore be different depending on whether the instruction is to ignore or stop to the signal.

Here, we hypothesize that releasing the assumption that decision is best captured by movement rather than visuomovement neurons would allow a more general understanding of the relationship between automatic and volitional influences in decision and facilitate model generalization across tasks. We propose to translate the insights gained from the countermanding modeling literature into the modeling framework that has been successful in accounting for saccadic inhibition (Bompas & Sumner, 2011). In this model, visuo-oculomotor decision is explicitly mapped onto the activity of visuo-movement neurons, receiving distinct influences of transient automatic and sustained goal-directed inputs. This separation adds versatility at the cost of mathematical elegance. However, perhaps counterintuitively, although this approach introduces more parameters, it also allows a more constrained and conservative approach to prediction and testing, because it clarifies which parameters should not be allowed to change between different tasks, for instance when only varying the stimuli or only the instructions. Importantly, this model was not originally developed to capture saccadic inhibition, but it readily did so when tested against the relevant experimental conditions. It was designed to account for other typical aspects of oculomotor control, including express saccades, anti-saccades, variation of target probability and the gap effect, using the basic characteristics of exogenous and endogenous neural signals and lateral inhibition in the intermediate layers of the SC (Trappenberg et al., 2001). Although originally based on superior colliculus, the model architecture is also more general because similar behavioral phenomena and model principles extend to manual responses (Bompas et al., 2017).

In **Section 2**, we first employ the latest models applied respectively to the stop task and saccadic inhibition and test the direct generalizability of each model to the conditions to which it had not been previously applied, using changes to inputs consistent with the internal logic of each model and as inspired by the alternative model. We do this both to learn how different implementations of bottom-up and top-down signals map onto existing behavioral and neuronal data, and to derive testable hypotheses.

From this initial exercise, we learn that separate transient and sustained signals are important, and we make two key predictions to test empirically. First, if decision dynamics are indeed best captured by visuomotor neurons, the early interference effects should be the same whether the instruction is to stop or ignore the signal. More specifically, the time at which the two distributions (in the presence and absence of signal) depart should be aligned across tasks. To confirm this, we designed three experiments combining saccade countermanding and saccadic inhibition paradigms using the same stimuli and participants but varying the instruction (**Sections 3 and 4**).

The second prediction is that, when using a model appropriately distinguishing automatic and selective drives, stopping behavior should be predictable from the parameters obtained from basic oculomotor behavior: We should not need a new top-down element for countermanding, and we should not need to fit the model to the stopping behavior itself. To test this, we extract the parameters from the conditions with simple saccades and saccadic inhibition (or inherit them from previous work), and we then test whether behavior in the stopping condition naturally follows **(Section 5)**. It is worth emphasizing this point, because a generalizable model is bound to have multiple parameters. Crucially we do not allow them to vary when transferring across tasks.

## 2. Model exposition and predictions

In this section, we first describe the best current models for saccadic countermanding and saccadic inhibition. Using published parameters from two separate studies, we simulate RT distributions from each model under the condition it has modeled before, as well as the alternative condition. We are testing whether each model qualitatively captures the shapes of the distributions in each task, as shown in Figure 1 (since we are inheriting parameters from different literatures, we do not use quantitative measures at this stage). To illustrate the key properties allowing a model to capture both response distributions, we then describe how the least generalizable model is improved by inheriting a property from the most generalizable one, i.e. an explicit dissociation between transient automatic and sustained goal-related inputs. We then present the simulated firing rates from each model in the countermanding task, emphasizing their similarity with neuronal recordings inspiring previous models of countermanding (Boucher, Palmeri, et al., 2007; Lo et al., 2009; Logan et al., 2015). Last, we derive two empirical predictions to be tested in the following sections.

### 2.1. Blocked Input 2.0

This model was developed to capture the stop signal task and is described in Logan et al. (2015). Although this model provides a similar fit to behavioral data as the simpler independent race model (or equally complex alternative models, see Logan et al., 2015), it better reflects the pattern of activity recorded in fixation and movement neurons within the frontal eye field of monkeys performing the stop-signal task. Being closer to the neuronal implementation of saccade planning opens the door to an increased ability to generalize to new tasks in ways that can be tested by both behavior and neurophysiology. BI 2.0 is a leaky accumulator with 2 nodes, representing the fixation and movement options, which are mutually inhibitory (Figure 2B). The go-signal is associated with a switch of input from the fix to the move node, occurring shortly after target onset (*D_move_* and *D_fix_* both less than 50 ms, here grouped as a single parameter *D* as they turned out to be numerically almost identical). The stop signal triggers two additional events: the fixation node quickly receives excitatory input again (following about the same delay D), then the input to the move node is switched off (‘blocked’) by a stop module (some *D_control_* delay after the signal; see Figure 2B right-hand blue panel). Node activity *a* directly maps onto firing rate, and evolves over time according to the following equation:

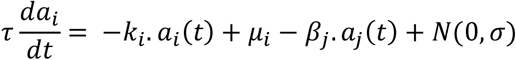

with *i* being either fixation or move node and *j* being the other node, *k* representing leakage, *μ* the intensity of inputs projecting from other areas, β the weight of inhibition from the other node and *σ* the amplitude of normally distributed noise added independently to each time step.

The most straightforward generalization of the model to the IGNORE instruction is to assume that visual inputs should be the same irrespective of the instruction, and only control inputs would be allowed to change. Thus, the presence of the signal requires fixation input to return identically as in the STOP condition, while the absence of the instruction to stop requires that move input is not blocked (Figure 2B red shaded panel). By default, we will assume that visual delays are unaffected by instruction, as we are not making quantitative latency comparisons in this section (we return to the question of visual delay below).

### 2.2. 200N-DINASAUR

This model was initially developed by Trappenberg et al. (2001) to extract and simplify key features of the SC based on both known neurophysiology and established principles of leaky interactive accumulators (Usher & McClelland, 2001). Subsequently, Bompas and Sumner (2011, 2015) and Bompas et al. (2017) showed that it predicted the characteristic dips of saccadic inhibition (the model is conceptually similar to the explanation given for saccadic inhibition by Reingold & Stampe, 2002), and in return these dips directly specify the delay time for exogenous input.

200N-DINASAUR shares many features with Blocked Input as both are noisy leaky competing accumulators. DINASAUR has 200 nodes representing the horizontal dimension of the visual field, and the average spiking rate *A_i_* of neuron *i* is a logistic function of its internal state *u_i_*:

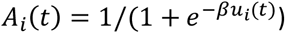

while *u_i_* varies across time *t* depending on normally distributed noise as well as the input received, either external to the map (endogenous or exogenous) or internal via lateral connections:

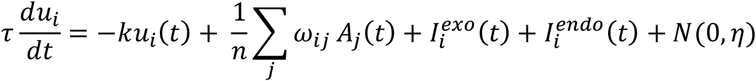

A key aspect of DINASAUR is that visual events can trigger two types of inputs: exogenous inputs (transients tied to visual changes) and endogenous signals (later, sustained and linked to the instructions). Of course, this is still a gross simplification of the many sensory pathways (exogenous) and other pathways (endogenous) that feed oculomotor planning. Endogenous inputs vary as step functions (similar to inputs in the Blocked Input models), while exogenous inputs are transient, reaching their maximal amplitude (*a_exo_*) at *t* = *t_onset_* + *δ_vis_*, and then

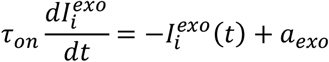

Exogenous inputs are tied to visual stimuli (e.g. targets, distractors or stop signals) and our key assumption is their properties (delay, strength, temporal profile) are not affected by instructions. Exogenous inputs also allow the model to produce express saccades (early mode at 70-110 ms on Figure 3B,D). All inputs have Gaussian spatial profiles (with SD *σ*): are maximal at the targeted nodes but also affect nearby nodes. Lateral connections show a Gaussian spatial profile that changes from positive (excitation) at short distance to negative (inhibition) at longer distance, described by the following equation:

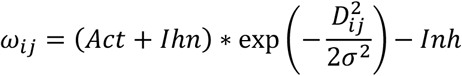

**Figure 3.**
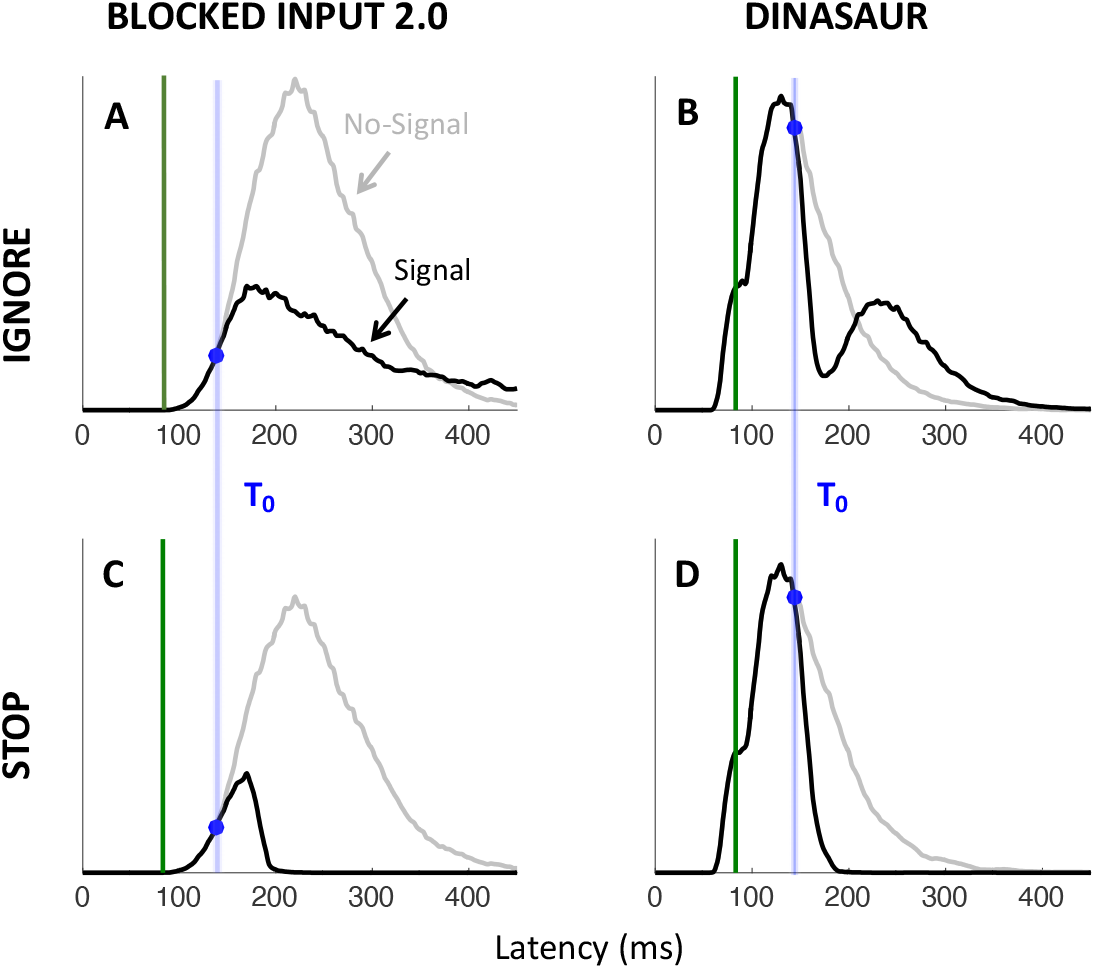
Simulated RT distributions from 10,000 trials using Blocked Input 2.0 (A,C) and 200N-DINASAUR (B,D) for signal onset (green line) at SOA 83 ms. Blue shaded areas indicate those instantiations of each models as published. Red shaded areas indicate predictions for new conditions based on the assumptions described in Figure 2. The DINASAUR model (with blocked input for stopping) captures well the typical pattern of results obtained in both paradigms. Blocked Input 2.0 (with automatic fixation activity for ignore conditions) is not able to produce the sharp dips expected from the saccadic inhibition literature (but see Blocked Input 3.0 and Figures 4-5). Both models predict a perfect alignment across instructions of the time when the signal RT distribution (black) departs from the NO-SIGNAL RT distribution (grey), indicated by the blue dots (T_0_) and highlighted by blue vertical bars. Note that the difference in mean and variance of the RT distributions between the models simply reflects the parameters inherited from previous publications; they have never been fitted to the same behavioral distributions. Relatedly, the position of T_0_ (blue dots) relative to the baseline distribution merely depends on where that distribution lies relative to signal onset (the SOA). The important aspect here is generalization ability of each model across instructions.

In 200-DINASAUR, the NO-SIGNAL condition is characterized by a single exogenous (visual) transient from target onset (occurring *δ_vis_* after target onset), shortly followed by a switch of endogenous support from fixation to target (*δ_endo_* after target onset). The SIGNAL-IGNORE condition differs from the NO-SIGNAL condition solely by the presence of a second visual transient, triggered by the signal appearing (the instruction being the same, no alteration of the endogenous inputs is assumed).

To generalize the model to SIGNAL-STOP conditions, we assume that only the endogenous input should differ from the SIGNAL-IGNORE condition, since the visual display is identical and only the instructions differ. Following the logic of Blocked Input 2.0, the endogenous input to the target is switched off (blocked) *δ_endo_* after the stop-signal, while the endogenous input to the fixation is switched on again. This amendment is fully consistent with the way endogenous signals are typically switched on and off in the DINASAUR model. Although the timings of these two events could in theory be free parameters and differ between peripheral and central nodes, in this section we use for parsimony a single *δ_vis_* parameter inherited from the SIGNAL-IGNORE condition, and a single *δ_endo_* parameter for both target and fixation nodes (with the delay between *δ_vis_* and *δ_endo_* inherited from previous work). Importantly, there is no need for this fixation drive to come back early, since the early stimulus-driven effect of any stimulus is already captured by the exogenous signal.

### 2.3. Generalization to new paradigms from Blocked Input 2.0 and 200N-DINASAUR

To test the generalization from both models to new tasks, we inherit as many parameter values as possible from previous publications, and make changes only where dictated by stimulus arrangement or the logic outlined above. For Blocked Input 2.0, parameter values are given in Table 1 and come from Monkey C in Logan et al. (2015) as its results were always shown first in their article. Using parameters from Monkey A did not alter our conclusion in any respect. For DINASAUR, parameter values are given in Table 2, and come from Bompas and Sumner (2011).

**Table 1.**
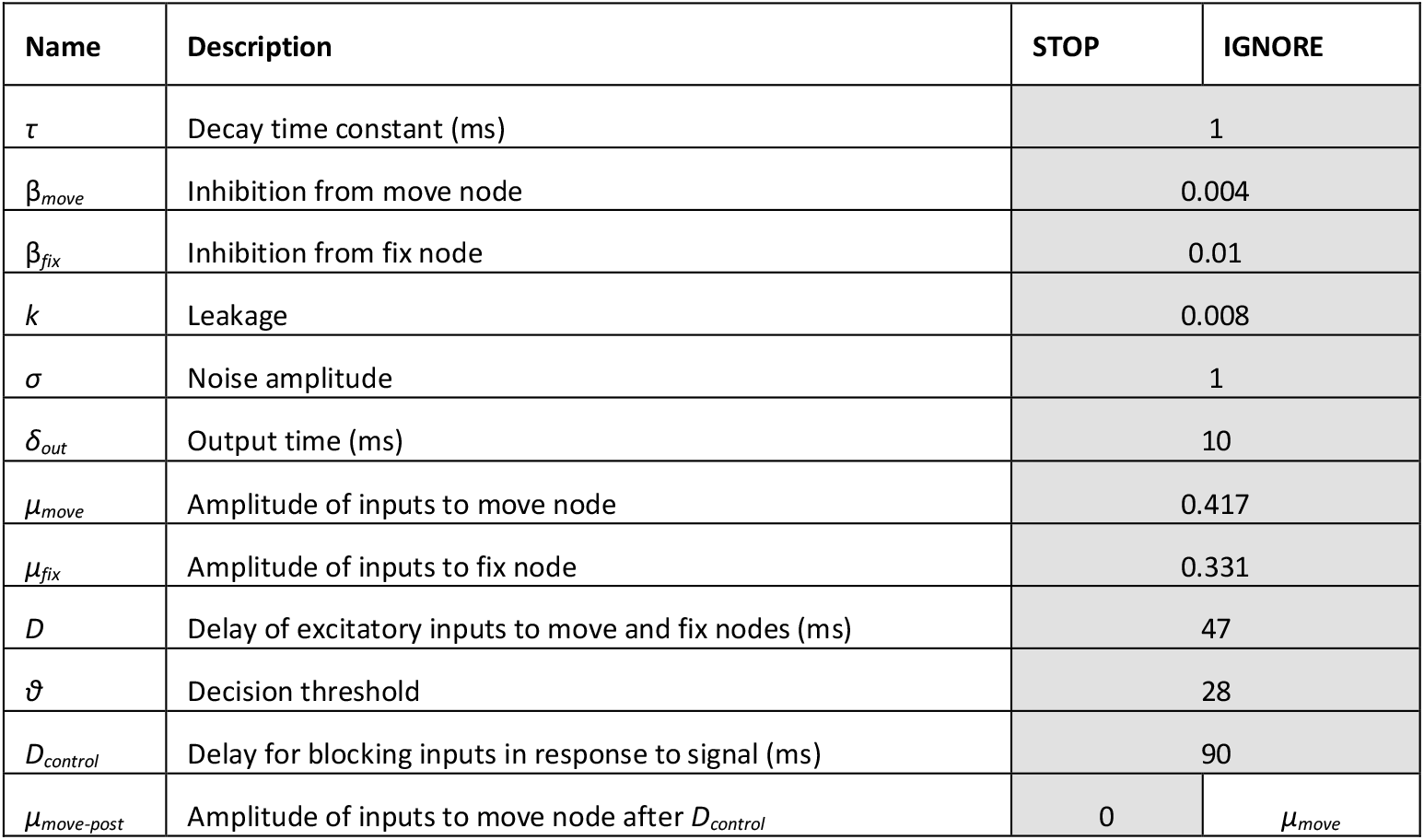
Model parameters for Blocked Input 2.0, as used in Figures 2 and 3. Grey boxes indicate parameters that were inherited from Monkey C in Logan et al. (2015), and correspond to the STOP instruction. The only alteration is that, in the IGNORE condition, μ_move_ remains up whether a signal appears or not (white box) but no new parameter is introduced.

| Name | Description | STOP | IGNORE |
| --- | --- | --- | --- |
| $\tau$ | Decay time constant (ms) | 1 | |
| $\beta_{move}$ | Inhibition from move node | 0.004 | |
| $\beta_{fix}$ | Inhibition from fix node | 0.01 | |
| $k$ | Leakage | 0.008 | |
| $\sigma$ | Noise amplitude | 1 | |
| $\delta_{out}$ | Output time (ms) | 10 | |
| $\mu_{move}$ | Amplitude of inputs to move node | 0.417 | |
| $\mu_{fix}$ | Amplitude of inputs to fix node | 0.331 | |
| $D$ | Delay of excitatory inputs to move and fix nodes (ms) | 47 | |
| $\vartheta$ | Decision threshold | 28 | |
| $D_{control}$ | Delay for blocking inputs in response to signal (ms) | 90 | |
| $\mu_{move-post}$ | Amplitude of inputs to move node after $D_{control}$ | 0 | $\mu_{move}$ |

**Table 2.** Model parameters for 200N-DINASAUR, as used in Figures 2 and 3. Grey boxes indicate those parameters unchanged from Bompas & Sumner (2011). The IGNORE condition is identical to previous work, except the distractor is now central instead of opposite to the target. The STOP condition differs from the IGNORE condition only in the endogenous response to the signal onset (white boxes) but no new parameter is introduced.

| Name | Description | IGNORE | STOP |
| --- | --- | --- | --- |
| $ECC_{dist}$ | Distractor eccentricity in SC (mm) | 0 | |
| $ECC_{targ}$ | Target eccentricity in SC (mm) | 1.76 | |
| $\beta$ | Steepness of spiking function | 0.07 | |
| $\tau$ | Decay time constant (ms) | 10 | |
| $\tau_{on}$ | Transience of exo inputs | 10 | |
| $Act$ | Short-range activation | 40 | |
| $Inh$ | Long-range inhibition | 55 | |
| $\sigma$ | SD of spatial profile for lateral connections and inputs in SC (mm) | 0.7 | |
| $k$ | Leakage | 1 | |
| $\eta$ | Noise amplitude | 50 | |
| $Th$ | Decision threshold | 0.85 | |
| $\delta_{out}$ | Output time (ms) | 20 | |
| $\delta_{vis}$ | Visual delay (ms) | 50 | |
| $\delta_{endo}$ | Endogenous delay (ms) | 75 | |
| $a_{exo}$ | Amplitude of exo inputs | 80 | |
| $a_{endo,target}$ | Amplitude of endo inputs to the target | 14 | |
| $a_{endo,fix}$ | Amplitude of endo inputs at fixation | 10 | |
| $a_{endo,target-post}$ | Amplitude of endo inputs to the target after SOA + $\delta_{endo}$ in signal trials | $a_{endo,target}$ | 0 |
| $a_{endo,fix-post}$ | Amplitude of endo inputs at fixation after SOA + $\delta_{endo}$ in signal trials | 0 | $a_{endo,fix}$ |

As expected, both models capture well the paradigm to which they have been applied previously (blue shaded panels on Figure 3). When using the published parameters, the SIGNAL-IGNORE scenario in Blocked Input 2.0 was not able to produce the characteristic phenomenon of saccadic inhibition: dips in the distribution (Figure 3A). Instead, the model predicts only a partial recovery from the interference, leading to many saccades being inhibited (51% for Monkey C, 78% for Monkey A), despite the instruction to ignore. Clearly, the prolonged interference from the sustained input triggered by the signal onset prevents the recovery of many saccade plans. In contrast, integrating the main idea from Blocked Input into the endogenous node within DINASAUR provides good generalization between IGNORE and STOP conditions (Figure 3B and D). Despite these differences, we note that the time of early interference (blue vertical line on Figure 3) is aligned across tasks for both models. This directly derives from our assumption that visual delay is not modulated by instruction and we will come back to this in Section 2.6.

### 2.4. Blocked Input 3.0

Although Blocked Input 2.0, like DINASAUR, contains both visual and control inputs, it was unable to generalize to the IGNORE instruction, at least not under the most straightforward assumptions. The main reason for this is the way visual signals are conceived in the model. Currently, these are simply tied to the presence of a stimulus in the neuron’s receptive field: they are on shortly after the stimulus is on and remain on until shortly after the stimulus goes off. As developed earlier, from the perspective of DINASAUR, inputs like this resemble the sum of two separate drives: fast transient inputs which would happen irrespective of the task (exogenous), followed by sustained inputs related to the task demands (endogenous). Here we hypothesize that the merging of these two influences within one parameter stands in the way of the generalization to the ignore condition, in which the same visual events occur but are associated with a different instruction.

To illustrate this, we test a simple upgrade of the Blocked Input 2.0 model. Taking inspiration from DINASAUR’s ability to capture the saccadic inhibition paradigm, we introduce a similar split between fast exogenous and slower selective signals into Blocked Input, which now allows the amplitude of these 2 streams of signal to vary independently as a function of the instruction. Figure 4 illustrates the relationship across all three models discussed in the present article.

**Figure 4.**
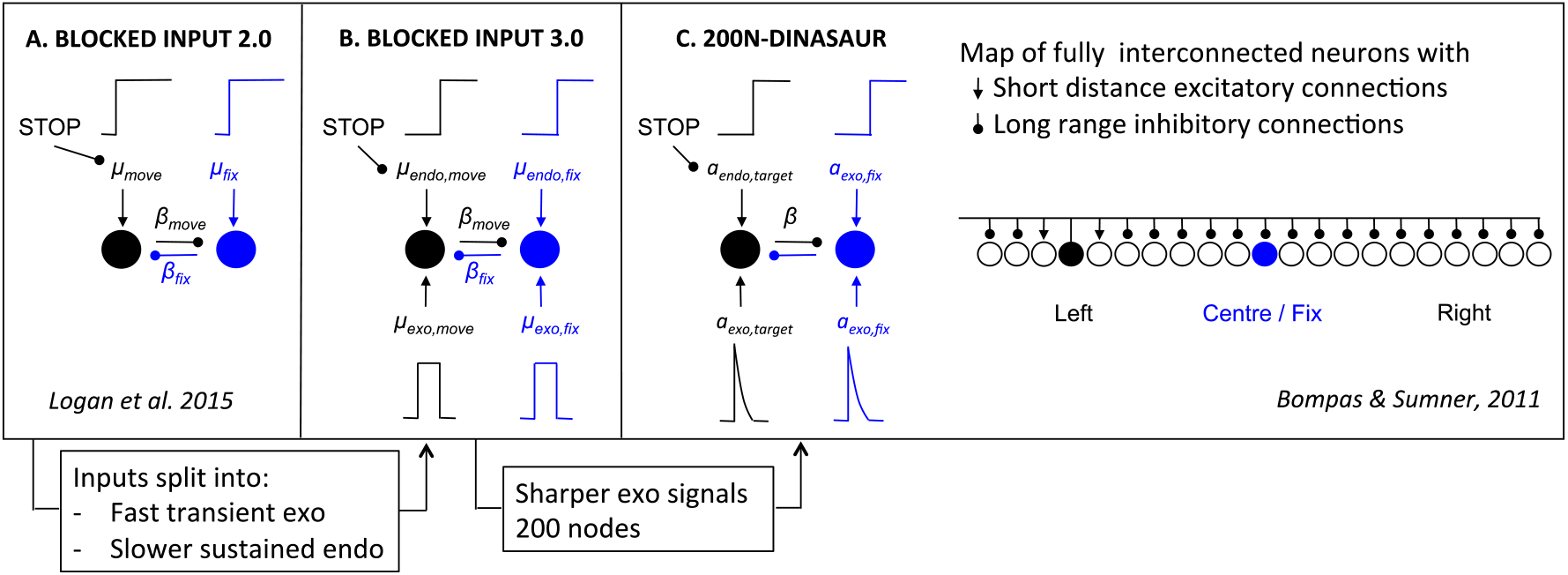
Overview of models and their relationships. **A.** Blocked Input 2.0 as in Logan et al. 2015**. B.** Blocked Input 3.0 integrates aspects of DINASAUR into Blocked Input 2.0 in an attempt to capture the SIGNAL-IGNORE condition. Its inputs are split into two conceptually different streams: a fast and transient drive tied to visual onsets (exogenous) and a slower sustained drive tied to instructions (endogenous). **C.** 200N-DINASAUR is a map of fully interconnected neurons representing part of the left, central and right visual fields, invented to capture simplified SC dynamics. The temporal dynamics of its exogenous signals (quick growth and exponential decay) is a key factor for creating sharp dips and quick recovery.

Table 3 presents the parameters specific to Blocked Input 3.0. We first attempt to inherit all parameter values from Blocked Input 2.0, without adding any new free parameter. In order to leave the NO-SIGNAL distribution unchanged between Blocked Input 2.0 and 3.0, we set the duration of exogenous signals as the difference between *D_control_* and *D*. Therefore, the inputs to the target node following target onset are the same under both models (a step function starting after delay D, Figure 5A-B). As can be seen on the simulated RT distributions (Figure 5C), this variant improves on Blocked Input 2.0 in that most saccades now recover from distractor interference in the IGNORE condition, which is crucial to observe dips, the hallmark of saccadic inhibition. The reason for this improved recovery is that the bottom-up signal associated with the return of fixation is now temporary (discontinued blue line on Figure 5A), rather than sustained (compare with Figure 2B).

**Figure 5.**
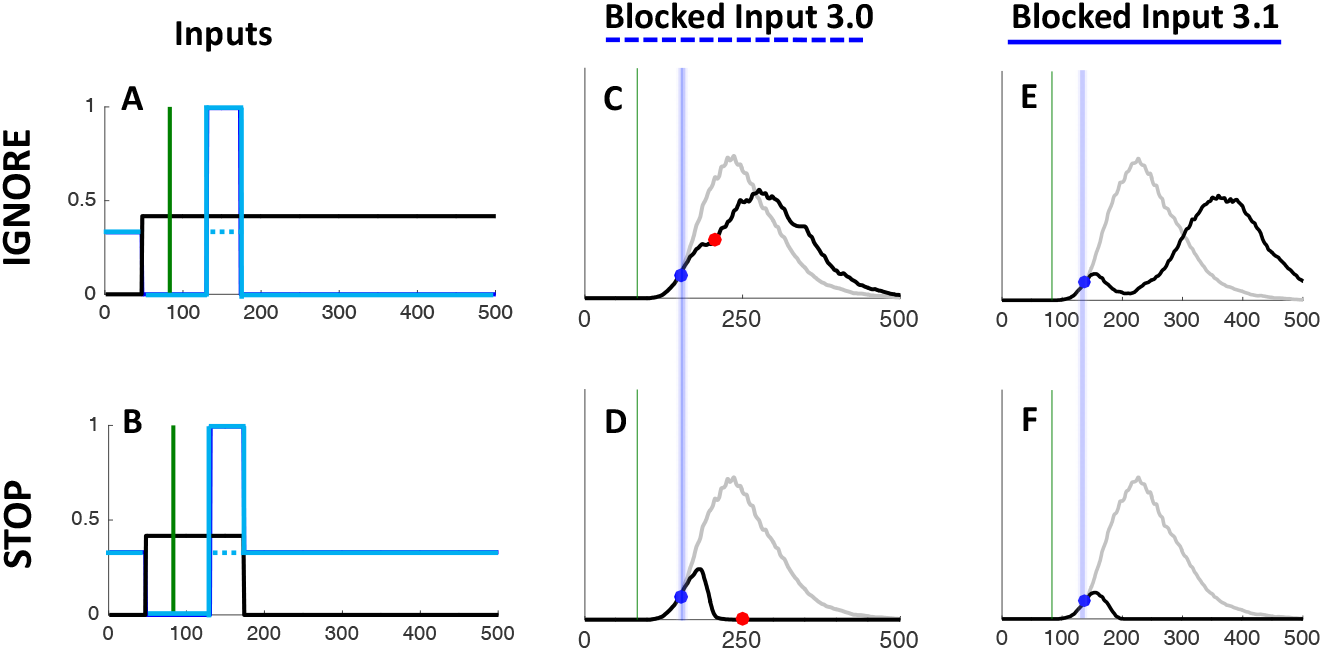
Inputs and simulations from Blocked Input 3.0 and 3.1. **A-B.** In the most straightforward generalization from Blocked Input 2.0, we assume in Blocked Input 3.0 that the transient visual signals associated with signal onset are the same size as the original fixation inputs in Blocked Input 2.0 (discontinuous blue line). Blocked Input 3.1 assumes that the transient activity from the signal is larger (in this case 3 times higher) than the baseline fixation amplitude (continuous blue line). **C.** Simulated RT for Blocked Input 3.0 shows some dip, but this remains very shallow. **D.** The STOP condition for Blocked Input 3.0 is the same as for Blocked Input 2.0. **E-F.** Simulated RTs for Blocked Input 3.1 now show a clear dip and recovery as expected in the SIGNAL-IGNORE condition (E), while still capturing the SIGNAL-STOP condition (F).

**Table 3.** Description and values of new parameters introduced in Blocked Input 3.0 and 3.1. Blocked Input 3.0 assumes all parameter values are equal to published values from Blocked Input 2.0 (grey boxes), while Blocked Input 3.1 adds one free parameter: the amplitude of the exogenous input triggered by signal onset (white box).

| Name | Description | 3.0 | 3.1 |
| --- | --- | --- | --- |
| $\mu_{exo,move}$ | Amplitude of exo inputs to move node | $\mu_{move}$ (from 2.0) | |
| $\mu_{exo,fix}$ | Amplitude of exo inputs to fix node | $\mu_{fix}$ (from 2.0) | $\mu_{fix} * 3$ |
| $\mu_{endo,move}$ | Amplitude of endo inputs to move node | $\mu_{move}$ (from 2.0) | |
| $\mu_{endo,fix}$ | Amplitude of endo inputs to fix node | $\mu_{fix}$ (from 2.0) | |
| $D$ | Delay of exogenous inputs (ms) | $D$ (from 2.0) | |
| $D_{Control}$ | Delay of endogenous inputs (excitatory and inhibitory, ms) | $D_{Control}$ (from 2.0) | |

However, the simulated dip remains much shallower than in behavioral data. In Blocked Input 3.1, we therefore decoupled the amplitude of exogenous and endogenous signals, to allow the exogenous transient signals to be larger (continuous blue line on Figure 5A). For instance, multiplying the exogenous signals by 3 creates much larger dips, now comparable in amplitude to typical data observed in saccadic inhibition. The STOP condition would now also contain this initial strong fixation signal, dropping back to the sustained level in Blocked input 2.0 after a short delay (Figure 5B). This slightly reduces the number of failed stops (Figure 5F). This upgrade is reminiscent of the Boosted Fixation model, also proposed (but less favored) in Logan et al. (2015). However, contrary to Boosted Fixation, the extra fixation drive here is only temporary.

Blocked Input 3.1 confirms that splitting signals into distinct transient exogenous and sustained endogenous drives is an important property for allowing the model to capture new tasks. Not only does this splitting allow us to decouple the amplitude of both drives, but it also creates a straightforward relationship between, on the one hand, visual events and exogenous signals, and on the other hand, the instructions and endogenous signals.

### 2.5. Comparison to recordings in FEF neurons

One of the strengths of Blocked Input 2.0 was its ability to capture not only monkey behavior but also that of fixation and movement neurons recorded within the frontal eye field of these monkeys, as previously published in Hanes et al. (1998) and Boucher et al. (2007). As explained above, DINASAUR appears better able to generalize across behavior in different tasks than Blocked Input 2.0. The next critical question is how well DINASAUR approximates activity in fixation and movement-related neurons. Figure 6 shows that firing rates from DINASAUR and Blocked Input models are quite comparable (**panels A-C**), and that DINASAUR accounts equally well for the growth and decay rates from FEF neurons during successful inhibition (**panel D**) highlighted in Logan et al. (2015). Figure 6 was designed to match Figures 13 and 14 in Logan et al. (2015) and the reader should refer to this work for a full justification.

**Figure 6.**
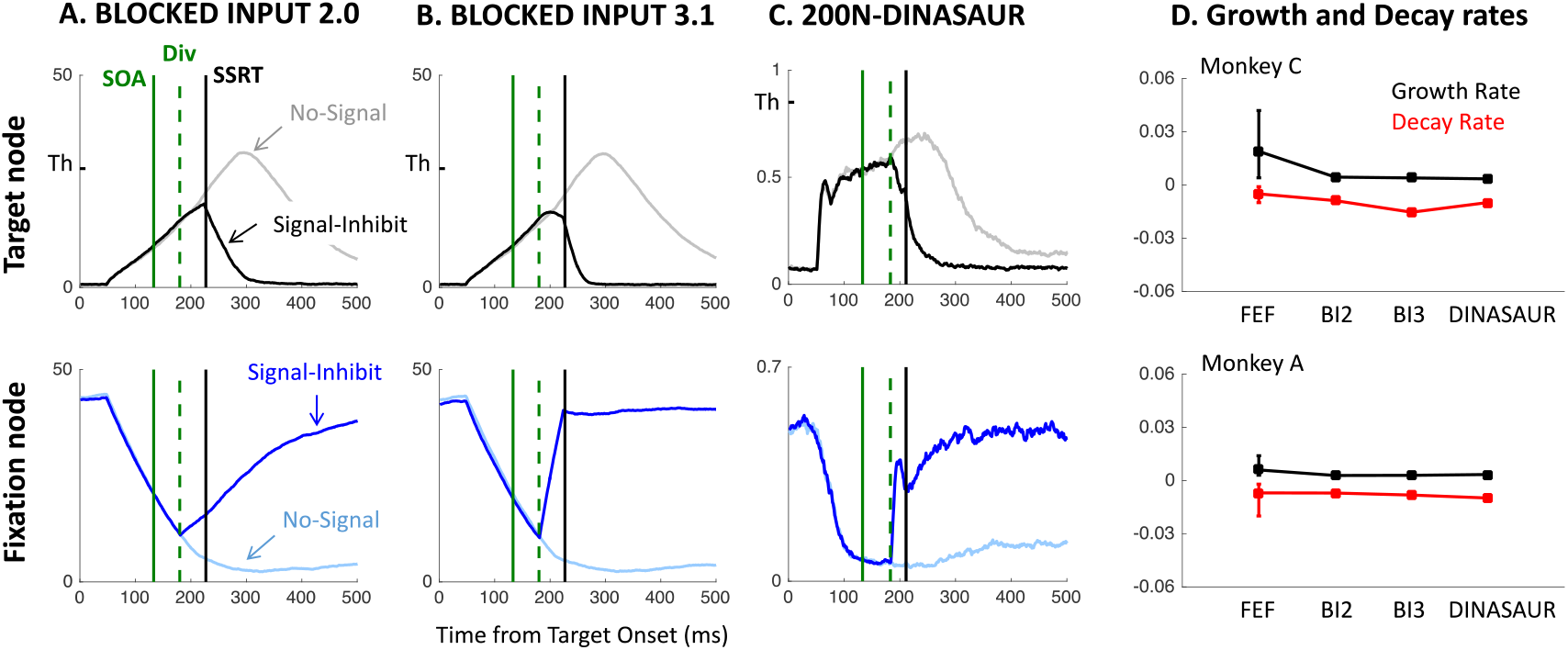
**A-C.** Mean firing rates from 1000 simulated trials using each model under the STOP condition, at the target and fixation nodes. The solid green line indicates the signal onset, here chosen at SOA 133, matching the experiments presented in section 4. The dashed green line shows the divergence time, i.e. the time at which this signal starts having an effect on the neuronal map, while the black vertical line indicates the SSRT, estimated from the simulated RT from each model. Activity was averaged across trials leading to successful inhibition (black and dark blue lines, Signal-Inhibit trials) and compared with “latency matched” No-Signal trials (grey and light blue lines; i.e. No-Signal trials in which latency is greater than SOA + non-decision time). On the y-axis for the target node, Th indicates the saccade initiation threshold (although this is not directly relevant for average firing rates, see text). **D.** Mean growth and decay rates from FEF neurons and simulations from each model (BI2 and BI3 refer to Blocked Input 2.0 and 3.1 respectively), using the same format as Figure 14 in Logan et al. (2015).

**Figure 7.**
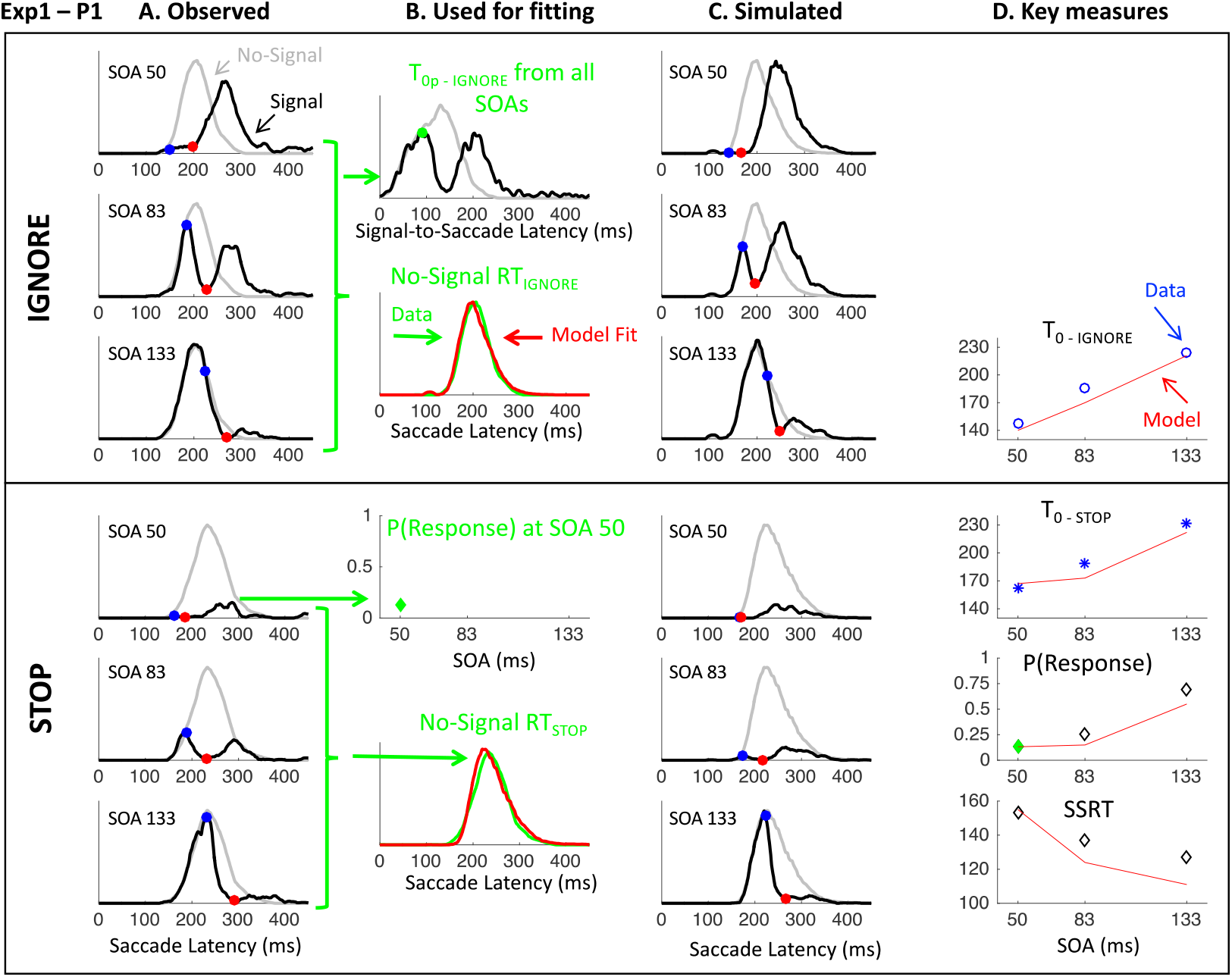
**A.** Latency distributions for Participant 1 in Experiment 1 across SOAs (rows) in the IGNORE and STOP contexts. Grey lines indicate distributions in which no signal was presented. Black lines indicate distributions of trials in which a signal occurred. Blue dots indicate the dip onset (i.e. where the two distributions first diverge); red dots show dip maximum. **B.** Green indicate the only data used for fitting the DINASAUR model: dip onsets from the IGNORE condition after pooling across all SOAs, NO-SIGNAL distributions from the IGNORE and STOP contexts, and the proportion of failed stops at SOA 50. Red lines show the fitted NO-SIGNAL distributions for this participant (see Section 5 for modeling details). **C.** Simulated RT distributions across all conditions for this participant. **D.** Observed (points) versus simulated (red lines) key measures at each SOA: dip onset in the IGNORE and STOP conditions, proportion of failed stops and SSRT, all in ms (see Figures S2-S3 for all individual data).

Panels A-C on Figure 6 contrast the mean firing rates between successful inhibition in SIGNAL trials and comparable NO-SIGNAL trials (i.e. NO-SIGNAL trials leading to a saccade being executed after the dip onset). In all models, target activity starts rising after a delay following target onset, while fixation activity decreases following fixation offset, irrespective of whether a signal is present or absent. On NO-SIGNAL trials, the fixation activity carries on decreasing (light blue lines), while the move activity carries on rising until it reaches a peak and then returns to baseline (grey lines). In neuronal recordings, this return to baseline is presumably related to triggering a saccade, and to mimic this effect in all our simulations, we interrupted the visual input to the peripheral target node each time a saccade was triggered in the model. This has of course no effect on the simulated RT distribution.

On SIGNAL trials, following the signal (green solid lines), activity rises again at fixation (dark blue lines), resulting in a decrease in move activity (mediated by lateral inhibition), further emphasized by the suppression of inputs to the move/target node. Panels A-C also show the divergence time (green dashed lines); the time at which this signal starts having an effect on the target node (the separation of dark and light blue lines). In all models, this time is equal to SOA + *δ_vis_*, and can be inferred from the RT distribution as dip onset time (*T_0_*) - *δ_out_*. All trials where the threshold is reached before this divergence time escape all influence from the signal and will therefore result in a failure to withhold the saccade (Signal-Respond trials). All trials where the threshold has not been reached by this time will be influenced by the signal to some extent. On some trials, the interference will be sufficient for the saccade to be correctly withheld (Signal-Inhibit category). On others, this interference may not be strong enough and the saccade is produced with a delay. This delay can be very short (as little as 1ms if the firing rate was very close to the threshold when the signal starts interfering), or much longer (up to 200 ms, see Bompas & Sumner, 2015). This variety means that recovery of saccades is already happening throughout the behavioral dips, rather than being restricted to the observed ‘recovery phase’. Although *δ_vis_* is kept constant and thus the interference starts at the same time on every trial, the dips in the generated behavioral distribution are more spread, matching those observed in empirical data.

The key difference between the models is that interference from the signal (the return of fixation activity and consequent lateral inhibition) increases in sharpness when going from Blocked Input 2.0 to Blocked Input 3.1 and to DINASAUR, illustrating the key property that makes DINASAUR able to produce sharp dips. Note that the downturn of target activity is already dramatic at the divergence time in DINASAUR, caused by the exogenous signal alone. In Blocked Input, the initial divergence is more subtle, and relies on the blocking of endogenous input for activity to take a severe downturn. Nevertheless, Panel B confirms the intuition from Logan et al. (2015) that a temporary boost of fixation following the signal (Blocked Input 3.1) would indeed capture neural dynamics.

While firing rates from Blocked Input 2.0 bear most resemblance to those motor neurons recorded in Monkey A, firing rates from DINASAUR resemble closely those visuo-movement neurons recorded in Monkey C (Figure 5 in Logan et al., 2015). Although there are important differences between the two neuronal populations (Ray, Pouget, & Schall, 2009), activity within both neuron types modulate at about the same time and show similar growth and decay rates, as stated in Logan et al. (2015). Figure 6D shows that DINASAUR provides growth and decay rates very similar to those in Blocked Input 2.0, accounting well for neuronal recordings in both monkeys. To construct panel D, we digitized the FEF data from Figure 14 in Logan et al. (2015), and ran simulations from each model following the same procedure as they used (see their Appendix C). Briefly, we simulated the models using the same SOAs and trial numbers as those from the FEF recordings (SOA ranging from 68 to 184 ms, and trial numbers varying from 61 to 130). For each SOA and monkey, the firing rate was averaged across the trials and divided by the initiation threshold. Minimum (M) and peak (P) mean firing rates were extracted, as well as the difference between these (D = P - M). The growth and decay rates were calculated for two sections of the curve, where the growth and decay are almost linear (i.e. the portion increasing from 25 to 75% of D (M + D * 0.25 to M + D * 0.75) for the growth rate, and the portion decreasing from 75 to 25% of D for the decay rate). It is clear that estimates from each model were within the range of estimates from neurons, similarly so across models.

The figure also shows the SSRT estimated from the simulated behavior for comparison (black vertical lines), using the integration method (Verbruggen, Chambers, & Logan, 2013). We can see that the SSRT follows the divergence time and the delay between the two has been referred to before as the cancel time (Boucher, Palmeri, et al., 2007; Lo et al., 2009; Logan et al., 2015). We will come back to the relationship between these two measures and *T_0_* in Discussion.

Lastly, note that, when averaged over a large number of trials, mean node activity in DINASAUR never reaches the initiation threshold, contrary to Blocked Input models. However, whether and when the *mean* activity reaches threshold is not directly relevant: in either class of model, the RT on each trial is determined by when the noisy activity reaches the threshold, and – due to the noise – this happens most of the time before the average trace reaches the threshold. Therefore, this apparent difference across models merely reflects the temporal profiles of accumulation (affected by the balance of self-excitation and leakage).

### 2.6. Empirical prediction: universality of dip onsets

Irrespective of how well each model performs overall, a crucial observation in all our model simulations is that the time point when latency distributions diverge is exactly the same under both instructions (blue dots and lines on Figure 3 and 5). This is a basic prediction as soon as the initial neuronal response to the stop signal is conceptualized as automatic, i.e. non-decision time is not modulated by context. In our previous work on saccadic inhibition, we have referred to this divergence point as dip onset or *T_0_* and, using DINASAUR, we have shown that *T_0_* - SOA directly reflects non-decision time (Bompas et al., 2017; Bompas & Sumner, 2011, 2015). Below we explain why the relationship between *T_0_* and non-decision time should hold overall irrespective of the model, and why we expect *T_0_* to remain unchanged across instructions.

#### Dip onset reflects non-decision time

The conceptual approach that dip-onset is a direct reflection of the sum of the sensory delay and the motor output delay (non-decision time) was validated by varying the luminance contrast and color of distractors (Bompas & Sumner, 2011), using dips as behavioral electrodes for precisely determining sensory delay. This relationship is not expected to be model-specific, since it depends simply on the logic of what non-decision time is – the portion of the RT that is not influenced by decision / selection processes (i.e. not influenced by a distractor signal). Neither should T_0_ *theoretically* depend on the shape of what follows – a sharp or gradual divergence or a true ‘dip’ (which implies divergence and then recovery). However, it should be noted that *T_0_* is only directly observable in simulations or data if the distractor signal SOA allows the dip to fall within the main body of the RT distribution and if there are enough trials to allow little or no smoothing (smoothing is known to anticipate dip onsets). Its estimate could therefore slightly vary across models depending on the shape of the distributions. In Figure 3, simulations from Blocked Input 2.0 and DINASAUR were smoothed using the same procedure as previous real data and produce *T_0_* respectively at 138 ms and 143 ms at SOA 83, irrespective of the instruction; that is respectively 55 and 60 ms following the distractor, while their respective non-decision times are 60 and 70 ms. Note that the differences in non-decision time across models are not relevant here as these result from fitting model parameters over completely different datasets and have never been contrasted before. What matters for now is that *T_0_* offers a good estimate of non-decision times for any model (but will often anticipate it by 5 to 10 ms depending on the RT distribution and smoothing).

#### Should T0 remain unchanged across contexts?

Earlier we described how mapping visuo-oculomotor decisions with the activity within visuomovement neurons predicted the early effects of the signal should temporally align between the ignore and stop contexts, while focusing on movement neurons would predict they should differ. Below we outline the intuitive reasons for expecting a difference and review the empirical evidence most closely related. One could argue that non-decision time may well differ under stop and ignore instructions, because of the associated attentional or strategic pro-active adjustments participants would likely make. Indeed, previous work using selective stopping paradigms (Bissett & Logan, 2014) has shown that, under the stop instruction, participants slow down to avoid making too many errors, in a similar fashion as when adjusting their behavior under accuracy versus speed instructions. It is therefore conceivable that *T_0_* would be longer under the stop condition compared with the ignore condition if non-decision time was to contribute to the overall slowing. On the other hand, the stop condition requiring more attention to be paid to the stop signal, it is also conceivable that this would lead to improved sensory processing of the signal (Elchlepp, Lavric, Chambers, & Verbruggen, 2016) and therefore possibly to a shortening of *T_0_* compared with a condition where the signal should be ignored.

However, previous research in the field of saccadic inhibition has consistently shown that *T_0_*, and therefore non-decision time, is mostly insensitive to pro-active slowing. For instance, (Reingold & Stampe, 2002) showed that dip timing was on average 4 ms *later* during pro-saccade blocks than during anti-saccade blocks, despite RTs being 100 ms faster. This being said, this difference was significant, which could suggest small but genuine modulations of non-decision time by instructions or ‘task-set’. In any case, these remain negligible compared with the modulations in decision time.

Although the SSRT has long been conceived as the delay required to inhibit action, it is now clear that a large proportion of this time is devoted to non-decision time, while the inhibitory component is rapid and late (Boucher, Palmeri, et al., 2007; Lo et al., 2009; Wong-Lin et al., 2010). SSRT is sensitive to the salience of the stop signal and insensitive to fixation offsets (Camalier et al., 2007; Morein-Zamir & Kingstone, 2006), just like *T_0_* in a saccadic inhibition paradigm (Bompas & Sumner, 2011; Reingold & Stampe, 2002). These findings suggest that SSRT likely behaves like *T_0_*, and therefore we expect the early part of the interference from stop-signals and distractors should be very similar in saccadic inhibition and countermanding. This leads to the strong prediction that *T_0_* should remain the same across contexts (within a few ms), providing the same stimuli are used and only the instructions differ. In Sections 3 and 4, we test this empirical prediction, which constitutes the 1^st^ step for our approach of unifying paradigms and models by regarding the first ‘inhibitory’ signal as fully automatic and therefore fully independent of instructions (note that this is actually an overly stringent definition of automatic; we will return in Discussion to the concept of conditional automaticity, whereby cascades of neuronal activation considered automatic are nevertheless modulated by context).

### 2.7. Modeling prediction: “one top-down fits all”

A second key consequence from Section 2 is that stopping does not necessarily need a specific cancel mechanism (with a specific strength and delay), but may be predicted from the combination of automatic interference and a switch of endogenous support from periphery to fixation. Crucially, once endogenous and exogenous signals are explicitly separated, like in DINASAUR and Blocked Input 3, they can be constrained from the NO-SIGNAL and the IGNORE conditions, and the generalization to the STOP condition should naturally follow. Although they could conceivably vary, a parsimonious hypothesis is that endogenous delays may all be captured by one variable, which constrains the latency of four events: 1) endogenous support for the target following target onset, 2) the removal of endogenous support for fixation following target onset, 3) the removal of endogenous support for the target following the signal under the stop instruction and 4) endogenous support returning to fixation following the stop instruction. This makes strong predictions when directly contrasting behaviors across conditions and paradigms, as this single parameter will now directly influence the NO-SIGNAL, SIGNAL-IGNORE and SIGNAL-STOP distributions across all SOAs.

Furthermore, this single endogenous delay is not even a free parameter in DINASAUR, but is defined as exogenous delay + a fixed delay of 25 ms. The assumption that *δ_endo_* directly depends on *δ_vis_* reflects the idea that both exogenous and endogenous delays in sensorimotor decision tasks are linked to sensory signals, but endogenous signals are filtered by task relevance (Bompas & Sumner, 2011). This filtering, imposed by the stimulus-response mapping, incurs an extra delay compared to raw visual signals (such as an onset at some location in the visual field) but the time at which these selective signals can be made available remains dependent on how fast the raw signals can reach these higher-level areas, i.e. the exogenous delay. Therefore, stronger signals will travel quicker within the brain, both straight to the decision area (*δ_vis_*), and via the filtering process for task relevance (*δ_endo_*). This 25 ms difference would in principle vary depending on the exact task and participants without changing the spirit of DINASAUR. The original model, inspired from the activity in SC neurons of monkeys, actually used a 50 ms difference (with δvis of 70, Trappenberg et al., 2001). Here the 25 ms is simply inherited from our previous modeling of saccades in humans (Bompas & Sumner, 2011).

In Section 5, we test whether DINASAUR can, under these strict assumptions and with the stopping behavior inspired from Blocked Input, capture all aspects of our data. We show that this is the case, as long as we allow two minor refinements to the model. Ultimately, our aim is not to pitch one model against another, but rather highlight key properties that inputs may have in order to reproduce the fine dynamics of visuo-oculomotor behaviors across a range of tasks. To a large extent, these considerations are independent of the peculiarities of each model’s architecture. From this perspective, it makes sense to test the predictions above using DINASAUR, as it has been used to model several other standard visuo-motor phenomena and its spatial extent lends itself to more hypothesis testing (see “Empirical predictions and future directions” section in Discussion), rather than upgrading Blocked Input further, which has been designed specifically to account for the countermanding task and had not been used for any other tasks until now. Therefore, in the remaining sections of the article, we use DINASAUR as the base model and inherit the spirit of Blocked Input for the behavior of endogenous signals during countermanding. This merger already captures the iconic behavior of the two paradigms as shown in Figure 3.

## 3. Empirical data – Methods

### 3.1. Rationale

The behavior of humans and monkeys during the saccadic stop task or saccadic inhibition has been described many times, forging strong expectations for what empirical distributions will look like in each paradigm separately (Figure 1) and justifying the modeling endeavor from both fields (Figures 2-3). However, in order to test the predictions laid out above, these paradigms must be tested on the same participants with the same stimuli, and with enough trials to support detailed distribution analyses and modeling. Ignore conditions have been used in stop paradigms, a paradigm known as “stimulus selective stopping” (see Bissett & Logan, 2014 for a review). This paradigm would typically introduce two types of signals, one requiring a stop and the other indicating the action should carry on (Verbruggen & Logan, 2009; Xu et al., 2017). However, the nature of the analysis performed in these previous studies was quite different to our present ambition.

As described above, our main aim for introducing new empirical data was two-fold. First, we aimed to test the prediction that the initial disruption to RT distributions is the same irrespective of instruction, suggesting that it is driven by automatic, rather than top-down inhibition (or a mixture of both). More specifically, this can be assessed by directly comparing dip onsets across instructions, as all models under this generalization hypothesis predicted perfect temporal alignment of dip onsets across conditions. Secondly, we aimed to test whether the later effects of the signal under each instruction can be captured within one single model with one set of parameters. This would suggest that distributions of failed stops can be fully predicted from the ignore condition by simply blocking the ability for saccades to recover, ultimately linking both phenomena to automatic interference from exogenous signals.

In order to answer these questions, we needed to directly compare aspects of the RT distributions under each instruction. However, there is no simple way of doing this without introducing additional complications. We therefore ran three experiments to provide converging evidence. We report these in the order they were implemented.

The easiest way to compare the two protocols using identical stimuli is to have separate blocks of trial where the instruction is to ignore the signal (pure saccadic inhibition design) and other blocks where the instruction is to stop to the signal (pure countermanding design). However, identical baseline (NO-SIGNAL) trials produce slower responses when participants know they might have to occasionally stop (as in a countermanding experiment) compared with when they are always allowed to ignore stimuli that come after the target (as in a saccadic inhibition experiment). This context-dependency is known as ‘proactive slowing’ (slowing of responses as a preparatory precaution given the possibility of having to stop, Verbruggen, Best, et al., 2014; Verbruggen & Logan, 2009). For this reason, we must compare SIGNAL-IGNORE and SIGNAL-STOP trials to their own NO-SIGNAL trials from the same block. But further, too much distribution shift between conditions would hamper direct comparison. When RTs are very quick, only short SOAs produce detectable dips (as later ones only affect the very tail of the distribution, where hardly any saccades occur). But very short SOAs are not optimal to study stopping behavior, as only very few fails would then be observed. To be able to compare behavior using an identical set of SOAs, we needed to ensure that the baseline distributions in the two contexts would overlap to considerable degree, even though some difference was inevitable.

In Experiment 1 we aimed to minimize the difference in proactive slowing between our two contexts, but at the same time we wished to compare ignore trials and stop trials that all had identical stimuli. We took inspiration from the selective-stopping paradigm and introduced two types of signal (white signals in 35% of trials and dark signals in 5%), but crucially we compared paradigms using the white signals only. The dark signals were present only to reduce differences in proactive slowing between the blocks. In the IGNORE context, participants were asked to ignore the white signal but stop to the dark signal, therefore encouraging some proactive slowing. In the STOP context, participants were asked to stop to the white signal but ignore the dark signal. Only responses to no-signal and white-signal trials were included in further analyzes.

We then validated our findings in two independent experiments. In **Experiment 2**, we simply removed the dark signal trials, creating a pure version of stop-task in half the blocks and a pure saccadic inhibition design in the other half. This removed the complication that participant had to remember two instructions simultaneously, but it created the expected large shift between the two baseline distributions, making the long SOAs inefficient in the IGNORE context, and the short SOAs suboptimal in the STOP context. Nevertheless, data were sufficient to act as a convergent validation.

**Experiment 3** was a standard selective stopping paradigm, where each block contained the same proportions of white and dark signals, one stimulus being associated with the ignore instruction and the other with the stop instruction. This mapping was alternated across blocks and the order counterbalanced across subjects (following Xu et al., 2017).

### 3.2. Participants

These experiments took a psychophysical approach in which few participants provided thousands of trials (between 5000 and 8000 each) to generate reaction time distributions, akin to neurophysiology studies that use non-human primates as subjects. The reason for this approach is that dips are a very robust phenomenon, found in every participant tested throughout the saccadic inhibition literature on humans and primates, while the critical aspect is the accurate estimate of *T_0_*, which benefits from collecting a large number of trials per condition. Thirteen participants (9 female) with normal or corrected to normal vision took part (4 in Exp 1, 5 in Exp 2 and 5 in Exp 3). One participant in Exp 2 was excluded because their accuracy on the stop task was around 2%.

### 3.3. Materials

A Tobii TX300 eye tracker with a 300 Hz sampling rate was used to collect saccade data. Participants were seated approximately 60cm from the screen where exact position of the eye in 3D space was calculated through algorithms supplied by the Tobii software for each time-point sampled. Eye position was calibrated using a 9-point calibration array at the start of every session and after every 600 trials (one block). A 23 inch (51 by 29cm) LCD screen with a 60Hz refresh rate was used to present stimuli. The lights in the room were switched off but the room was not in total darkness.

### 3.4. Stimuli and procedure

The two main trial types are illustrated in Figure 1 and 2A. Briefly, all trials began with a central fixation point, a white circle 0.4° visual angle in diameter (200 cd/m^2^), presented on a grey background for 700ms (58 cd/m^2^). This was immediately followed by a target with the same properties as the fixation point but either 12° visual angle to the left or right of the center of the screen on the vertical midpoint. For no-signal trials (60% of trials), the target appeared for 1000ms and no other stimuli were presented. Participants were instructed to fixate on the central fixation point and then saccade as quickly as possible to the target that appeared randomly on the left or right of fixation (in equal frequencies).

All experiments also contained trials in which the target was followed by a larger stimulus (1° diameter), either white (120 cd/m^2^, Figure 2A) or dark (9 cd/m^2^, not illustrated), appearing in the center of the screen after varying stimulus onset asynchronies and until the end of the trial (i.e. until the peripheral go-signal disappeared). The three experiments differed in the frequency of these white and dark signal trials, the associated instructions and the range of SOA covered, as detailed below.

In Experiment 1, 35% of trials contained a white signal and in half the blocks the instruction was to ignore these stimuli (thereafter called IGNORE blocks), while in the other half of blocks the instruction was to withhold the eye movement if these stimuli appeared (STOP blocks). The remaining 5% of trials were dark and were associated with the alternative instruction (stop in the IGNORE blocks and ignore in the STOP blocks). These were not analyzed and were added only to minimize the difference in pro-active slowing between blocks. Therefore, in the analyses below, the SIGNAL-IGNORE and SIGNAL-STOP trials contained the same visual stimuli (peripheral white discs followed by central white discs), while only the required responses varied. The SOA were 50, 83 and 133 ms (due to the 60Hz refresh rate).

Experiment 2 was identical to Experiment 1, except the dark stimuli (and any instruction about them) were absent, bringing the number of trials with a white signal to 40%.

Experiment 3 was identical to Experiment 1 except for stimulus frequency and additional SOA. White and dark signals occurred in equal proportion (20% each). In half the blocks, participants were instructed to ignore the white stimuli and stop to the dark one. In the other half, the instruction was reversed. SOAs were 50, 83, 133 and 183 ms.

All participants were instructed to ‘respond as fast as possible whilst minimizing errors’. At the end of each block participants were given feedback on mean reaction time, percentage of failed stops and percentage successful ignores for the relevant stimuli. Each participant completed a training session of 20 minutes. This was followed by over 5000 trials (8640 in Exp 1, 5472 in Exp 2 and 5760 in Exp 3), spread over 4 sessions. Each session in Experiment 1 contained a run of 3 blocks under one instruction followed by 3 blocks of the alternate instruction, presented in a counterbalanced order both within sessions and across participants. Each block was 15 minutes long, bringing each session to around 90 minutes. The same procedure was used in Experiments 2 and 3, except only 4 blocks (2 runs under each instruction) were run per session, bringing the session duration to 60 minutes.

### 3.5. Data Analysis

Response saccades were detected using a velocity criterion of 35°/s, an acceleration of 6000°/s, and an amplitude of at least 6° (halfway to the target). Trials were excluded if there was loss of tracking, blinks or small saccades (under 6°) in the period between target onset and response saccade onset or during the 500 ms following target onset in the absence of a response saccade. Each trial was visually inspected to ensure correct saccade detection by the algorithm and corrected where needed. Trials containing a saccade to the location opposite the visual target were also excluded, but these were extremely rare (less than 0.1%). Overall, this resulted in excluding on average 3% of trials (ranging from 0.3 to 5.3% of trials across all participants and experiments). Saccade latencies were calculated as the difference between target onset and saccade onset and then classified by trial type and context. All following analyzes are collapsed across left and right targets.

Next, saccade latency distributions were obtained for each participant for no-signal and signal trials for each SOA collapsed across all sessions, separated by instruction. Latency distributions were obtained with a bin size of 3.33ms (the refresh rate of the eye tracker was 300Hz). Given the difference in trial numbers between signal and no-signal trial-types, all distributions were scaled according to the number of trials still present within that condition after the exclusions listed above. Distributions of correct responses were then lightly smoothed using a Gaussian kernel with 7ms window size and 3ms standard deviation and interpolated to obtain 1ms precision, in line with Bompas et al. (2017) using similar trial numbers. Distributions using pooled data across observers and/or SOA used less smoothing (window = 5, SD = 1), in line with Bompas & Sumner (2011) using larger datasets. Note that for noisy distributions, smoothing is necessary to robustly extract dip onset, but also anticipates dip onset. When more trials are available, smoothing becomes less necessary and less desirable for this reason.

In order to determine the onset and peak amplitude of the dip in saccade latency distributions a distraction ratio was calculated for each time-bin of the latency distributions where at least 1 trial was present in the no-signal condition (e.g Bompas & Sumner, 2011; Reingold & Stampe, 2002). This distraction ratio is the proportional change in the number of saccades made in the signal-present distribution relative to the number in the no-signal distribution. This is calculated for each time bin as:

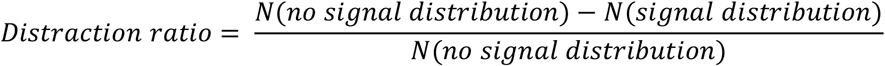

The peak dip amplitude was calculated as the first time point of the maximum of the distraction ratio where the difference in the two distributions was greater than 2 saccades and the ratio was greater than 20%. Onsets of dips were defined as the point at which the distraction ratio fell below 2% working backwards in time from the dip peak.

In Experiments 1 and 2, the analyzed IGNORE and STOP trials (white signals only) were collected in different blocks. Baseline distributions (NO-SIGNAL trials) were therefore analyzed separately for IGNORE and STOP blocks. In Experiment 3, there were four SIGNAL trial types to analyze: WHITE-IGNORE, DARK-IGNORE, WHITE-STOP, DARK-STOP. Analyses were first performed separately for white and dark stimuli but no statistical difference was observed (2-way repeated measures ANOVA was performed on dip onset times, with instruction and contrast as factors, revealing no effect of contrast: F(1,3) = 0.16, p=0.7; the effect of instruction is reported below, along with the other two experiments). This allowed us to pool across dark and light signals, leaving us with the same conditions as in Experiments 1 and 2: SIGNAL-IGNORE and SIGNAL-STOP. However, in Experiment 3, the same GO trials serve as baseline for both signal conditions, as these trials were interleaved.

For the essential question of whether dip onsets were aligned across tasks, we used bootstrapping to estimate the stability of any difference in estimated dip onset times within each observer and its mean at the group level (considering that our number of participants is small but our number of trials per participant is very high). The extraction of dip onset time was performed on the signal-to-respond latencies locked on signal onset, pooled across SOAs (i.e. the distributions in Figure 8D-F). For each participant, we generated 1000 surrogate distributions for each condition from the observed distributions (NO-SIGNAL and SIGNAL at each SOA, each under both IGNORE and STOP instructions), by randomly sampling the same number of trials from each original distribution with replacement. On each iteration, we applied the same dip onset extraction procedure as for observed data, subtracted the dip onset time under the surrogate IGNORE instruction (T*_0p-IGNORE_*) from that in the surrogate STOP condition (*T_0p-STOP_*), and calculated the 95% (uncorrected percentile) confidence intervals over these 1000 bootstrapped differences. Then, for each experiment, we averaged the bootstrap estimates across participant, producing 1000 estimates of the difference in mean *T_0p_* on each group. Each difference was considered insignificant when the 95% confidence interval included zero, under similar assumptions as those used to calculate a p-value.

**Figure 8.**
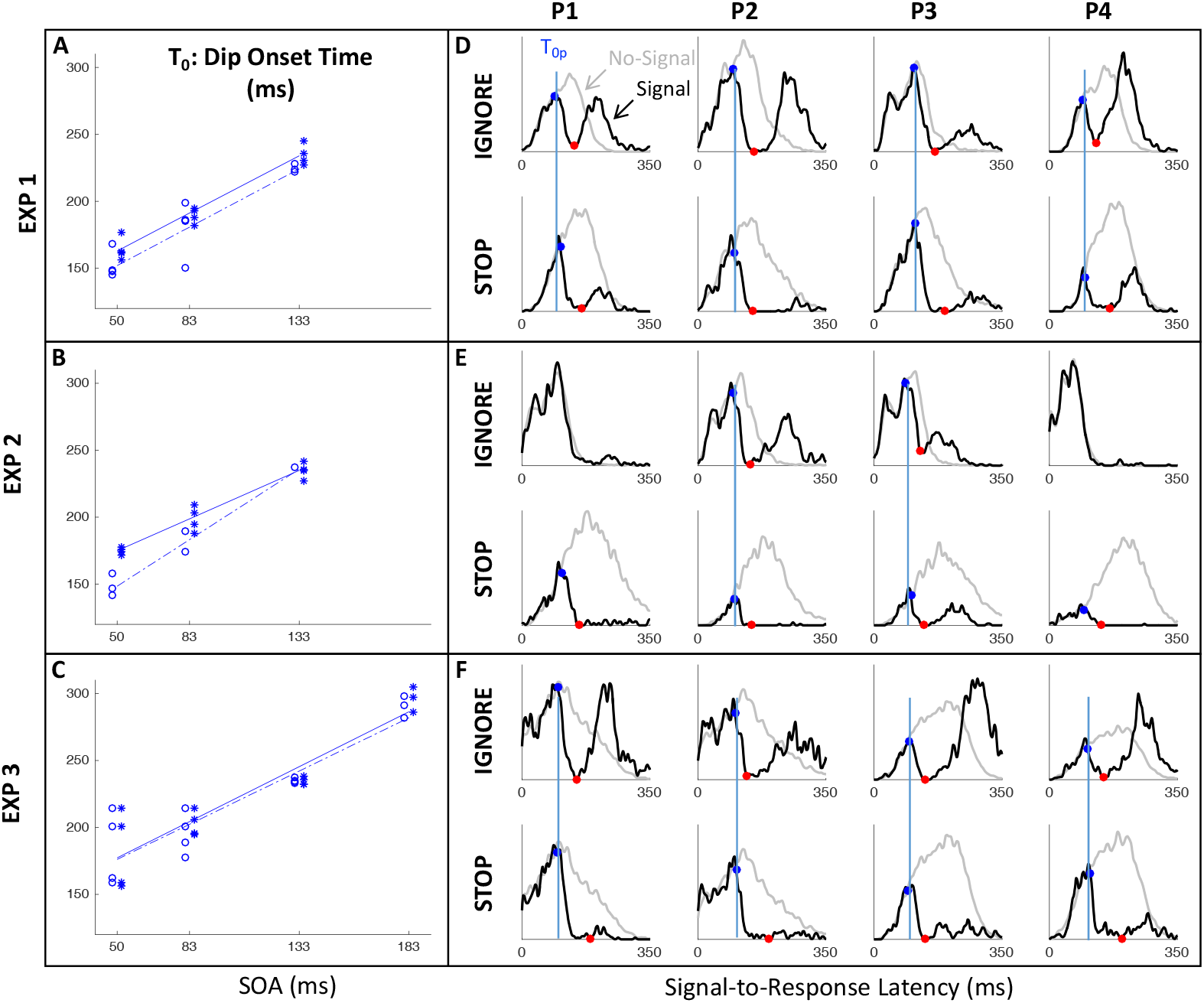
**A-C.** Dip onset times (T_0_) for each participant in the IGNORE (open circles) and STOP (stars) contexts of each experiment, along with regression lines across SOAs on each group (whenever sufficient data was available). As predicted, dip onsets are locked on signal onset and are temporally aligned between the IGNORE and STOP contexts, consistently across experiments. **D-F.** Overlap of dip timing between the IGNORE and STOP contexts in each experiment, highlighted by blue vertical bars. Distributions show saccade latency locked on signal onset, allowing pooling of trials across the three SOAs to best visualize the timing of dip onset (blue dots) and maximum (red).

## 4. Empirical data - Results

### 4.1. Latency distributions

Figure 7A shows the saccade latency distributions for a typical participant (P1 in Experiment 1) in each context and each SOA. Figures S1 in Appendix shows all individual distributions. As expected, the IGNORE context is characterized by dips in the distribution following signal onsets, comparable to those in previous studies of saccade inhibition (Bompas & Sumner, 2011; Buonocore & McIntosh, 2008, 2012; Buonocore & McIntosh, 2013; Edelman & Xu, 2009; Reingold & Stampe, 2002, 2004). The distributions of failed inhibitions in the STOP context also show dips, but these are followed by little or no recovery, indicating mostly successful stops in the latter part of each distribution. Although one can start to appreciate the temporal alignment of *T_0_* across contexts, this is more clearly illustrated by pooling across SOAs (section 4.2).

### 4.2. Temporal alignment of dip onsets across contexts

Figure 8A-C shows the expected strong linear relationship between dip onset and the timing of the signal. This locking of *T_0_* on distractor onset justifies pooling across SOAs based on time-since-distractor in order to improve the estimates of *T_0_* by using all the available data, a standard practice in many studies on saccadic inhibition (see section 3.5 and Reingold & Stampe, 2002). Figure 8D-F show these Signal-to-Response latency distributions for each participant and illustrate the temporal proximity of dip onsets across instructions.

Table 4 presents the key summary statistics across all three experiments (see Table S1 and Figures S2-3 for more). Across all three experiments, dip onsets were on average 5, 4 and 1 ms earlier under the IGNORE compared to the STOP instruction, but the 95% confidence intervals all included zero (this was also the case for each individual participant). We therefore concluded that there was no significant difference in *T_0p_* across instructions. Dip onsets in the present study are around 98 ms on average under the IGNORE instruction (102 under STOP), slightly later than reported previously, but it is known that stimulus properties affect dip onset (see e.g. Figure 6 in Bompas and Sumner, 2011), and the precise timing of its detection is affected by trial numbers and smoothing (Bompas et al., 2017). Dip maxima (red symbols) also occur at similar times in each context, though the exact timing of dip maximum is affected by the properties of the recovery, and thus less directly interpretable than dip onset.

**Table 4.** Summary of empirical measures (in ms) for individual participants and average (A) from each experiment. RT_No_ and SD_No_ are the mean and standard deviation of RT in the No-Signal condition. T_0p_ is the dip onset estimated from pooled distribution across all SOAs locked on signal onset (see Figure 8), except for participant 1 in Exp 2 for whom T_0_ at SOA 50 was used as this was the only SOA showing a dip. Values within brackets indicate the bootstrap 95% confidence interval. The last column indicates the observed difference in T_0p_ between the IGNORE and STOP instructions, along with the bootstrap 95% confidence interval on this difference. For the STOP condition, the % of failed stops for each SOA and the mean SSRT across SOAs are also provided. Note that, for Exp 3, the IGNORE and STOP conditions are interleaved, so there is only one No-Signal condition. See Table S1 in Appendix for mean RT, SD and SSRT at each SOA.

| Exp | P | IGNORE |  |  | STOP |  |  |  |  | Difference |  |
| --- | --- | --- | --- | --- | --- | --- | --- | --- | --- | --- | --- |
| | | $RT_{No}$ | $SD_{No}$ | $T_{Op}$ | $RT_{No}$ | $SD_{No}$ | $T_{Op}$ | % Error | SSRT | $T_{Op}$ | |
| 1 | 1 | 207 | 29 | 90 [64 107] | 241 | 36 | 105 [90 107] | 13-26-69 | 139 | 15 [-4 39] |  |
|  | 2 | 210 | 42 | 96 [62 102] | 239 | 58 | 99 [96 102] | 15-31-62 | 124 | 3 [-4 37] |  |
|  | 3 | 196 | 40 | 111 [79 117] | 242 | 64 | 112 [95 116] | 27-46-86 | 141 | 1 [-17 34] |  |
|  | 4 | 231 | 27 | 91 [81 99] | 261 | 34 | 98 [94 100] | 17-25-49 | 155 | 7 [-2 17] |  |
|  | A | 211 | 37 | 97 [83 102] | 245 | 51 | 103 [98 105] | 18-32-62 | 140 | 5 [-1 21] |  |
| 2 | 1 | 170 | 34 | 108 [102 135] | 288 | 71 | 109 [102 120] | 6-17-44 | 140 | 1 [-30 12] |  |
|  | 2 | 195 | 40 | 96 [84 103] | 280 | 60 | 98 [85 113] | 3-4-22 | 120 | 2 [-14 19] |  |
|  | 3 | 178 | 29 | 86 [76 106] | 290 | 68 | 101 [95 105] | 12-12-40 | 141 | 15 [-7 25] |  |
|  | 4 | 132 | 20 | NA | 289 | 60 | 95 [89 108] | 6-8-17 | 114 | NA |  |
|  | A | 169 | 39 | 97 [85 108] | 287 | 65 | 101 [96 107] | 7-10-31 | 129 | 4 [-7 18] |  |
| 3 | 1 | 223 | 63 | 98 [88 109] | 223 | 63 | 95 [89 110] | 31-46-66-89 | 122 | -3 [-16 18] |  |
|  | 2 | 247 | 74 | 102 [85 105] | 247 | 74 | 105 [92 109] | 23-40-57-85 | 134 | 3 [-8 19] |  |
|  | 3 | 293 | 38 | 97 [77 108] | 293 | 38 | 94 [89 111] | 7-11-19-72 | 153 | -3 [-14 29] |  |
|  | 4 | 284 | 40 | 105 [101 108] | 284 | 40 | 112 [96 117] | 10-20-26-81 | 153 | 7 [-10 13] |  |
|  | A | 262 | 62 | 100 [93 105] | 262 | 62 | 101 [98 109] | 18-29-42-82 | 141 | 1 [-5 12] |  |

As expected, strategic adjustments across tasks existed in Experiments 1 and 2, and were large in Experiment 2 (where the two contexts were kept fully separated). Since the baseline distributions differed depending on context, but the timing of the dips (relative to the signal) is similar across contexts, the dip is therefore earlier *relative* to the main mode of the distribution in the STOP context, and thus the height of the pre-dip distribution was normally smaller in the STOP context. This is just a consequence of the baseline distributions. The critical question here was whether the leading edges of the dips are coincident. The large differences in baseline distributions in Experiment 2 meant the visual signal often arrived too late to have much effect in the ignore condition, especially for the fastest participants (P1 and P4), consistent with previous work (Bompas & Sumner, 2011). Nevertheless, when dips were observed in both contexts, these were temporally aligned, like in Experiment 1 and 3.

### 4.3. Inhibition function, SSRT and dip recovery

We now focus on the typical metrics reported within the stop-task literature. The inhibition function (Figure 9A and Table 4) showed the expected increase in the proportion of failed-stops with SOA. At the shortest SOA under the stop instruction, our participants produced on average 13% errors. This number was higher in Experiments 1 and 3 (18%), where participants were required to switch between instructions, compared to Experiment 2 (7%), where instruction were kept separate across blocks as in the standard stop task. Figures 7 and S1 show that the latency distribution of failed stops is often bimodal. Indeed, for all participants in Experiments 1 and 3, and P3 in Experiment 2, there is a partial recovery from the dip even in the STOP context. This failure to inhibit the saccade on some trials well after the time when a participant is usually able to do so has been reported before (Akerfelt et al., 2006; Hanes & Carpenter, 1999). In our modeling, we will suggest it may indicate occasional failure to trigger the inhibition command (Emilio Salinas & Stanford, 2013; Skippen et al., 2019), possibly fueled by temporary confusion about which instruction applied (see section 5.3).

**Figure 9.**
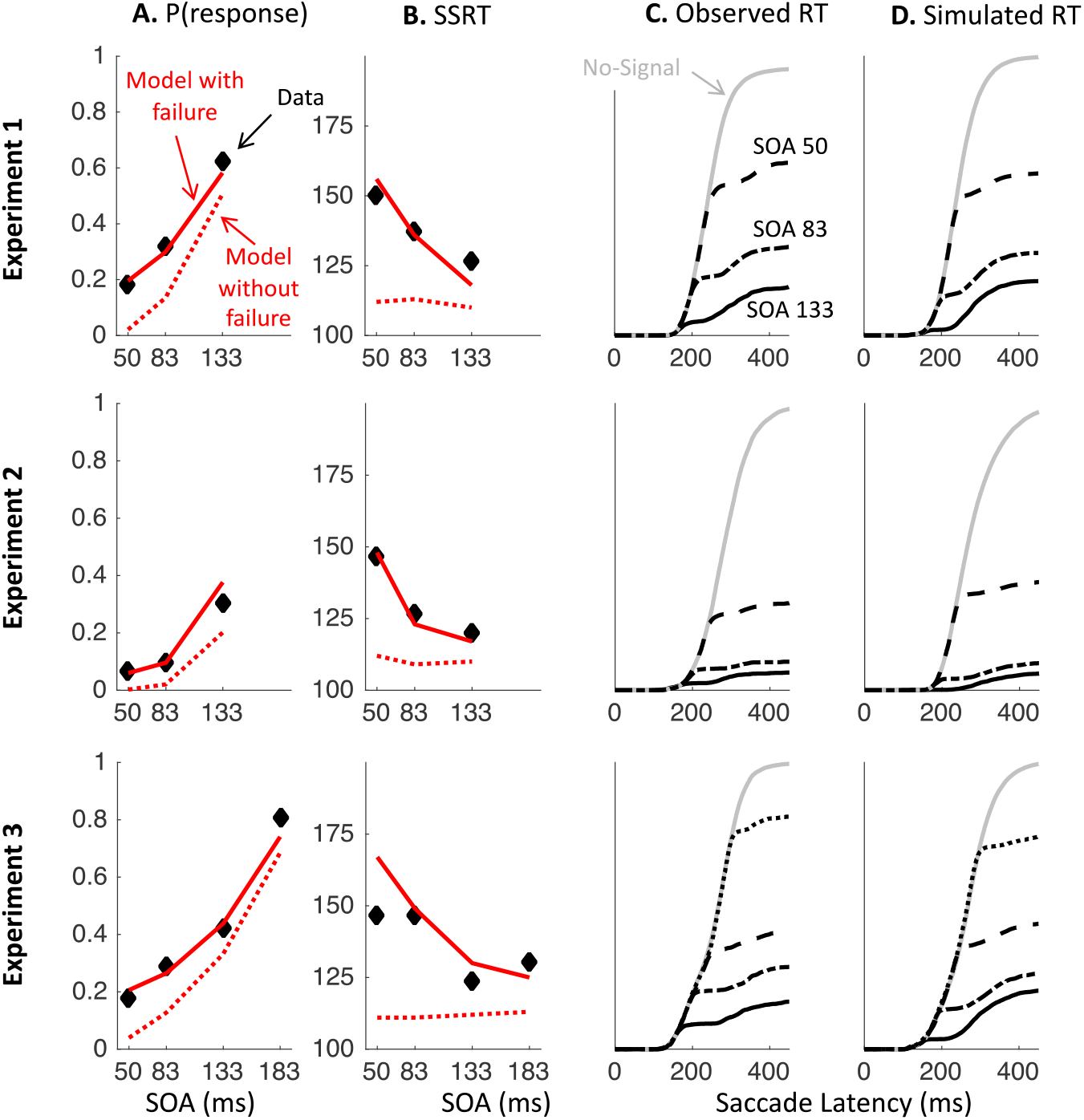
Traditional stop signal task measures from observed and simulated data. **A-B.** Proportion of failed stops (A) and stop signal reaction time (SSRT, B) across SOAs, from the pooled data across observers (black diamonds) and in DINASAUR simulations (red lines) with and without failure (continuous and dotted lines). The SSRT was calculated using the integration method (Verbruggen et al., 2013). **C.** Cumulative distributions for observed no-signal (light grey) and signal trials (black continuous, semi-dashed, dashed and dotted for SOA 50, 83, 133 and 183 respectively). **D.** Same as C for DINASAUR simulation (with failure), also pooled across observers. See Figure S3 in Appendix for individual data.

Table 4 and Figure 9B also show the SSRT estimates, obtained using the integration method (Verbruggen et al., 2013). These were comparable to previous reports for saccade countermanding in human (on average 134 ms, Hanes & Carpenter, 1999), i.e. about 32 ms after dip onset and 30 ms longer than in rhesus monkeys (Hanes et al., 1998; Hanes & Schall, 1995; Paré & Hanes, 2003). These showed a clear dependency on SOA (Figure 9B), as previously reported in the manual (Band, van der Molen, & Logan, 2003; de Jong, Coles, Logan, & Gratton, 1990; Logan & Burkell, 1986; Logan & Cowan, 1984; Matzke, Love, & Heathcote, 2017) and saccadic (Hanes & Schall, 1995) domains. As our modeling will suggest, this may also be entirely related to the partial recovery from dips in the stop task, and therefore down to the reliability with which the stop instruction is being applied.

We also plotted the cumulative distributions of RT (Figure 9C). Contrary to the custom in the stop-signal task literature, we did not normalize these on the number of saccades executed, which, in our eyes, would have masked the main feature of interest here: the exquisite overlap in the signal and no-signal distributions until the departure point (*T_0_*) and the dependency of this point on the SOA, both hallmarks of dips in the saccadic inhibition literature. Rather, cumulative distributions were normalized to the number of trials available in each condition.

## 5. Modeling Results

This section aims to test our prediction that a general model ought not to need specific parameters for countermanding, i.e. ought to be able to predict stopping behavior from parameters derived from basic behavior in baseline and ignore trials. To do this, we individually adjust three of the parameters in the DINASAUR model: the visual delay; the strength of endogenous signals during fixation; the strength of endogenous signals in response to the target. To further improve the fits to the no-signal distribution, we add two refinements to the model. The first is a holding period to account for strategic slowing down in the stop task. The second is a failure parameter, allowing a proportion of trials in the stop task to be effectively treated as ignore trials. These adjustments are illustrated in Figure 7, summarized in Table 5, and explained fully below.

**Table 5.** Parameters adjusted in 200N-DINASAUR to capture data from Experiments 1, 2 and 3 (see Table 2 for full list of parameters). Grey boxes indicate parameter values from Bompas & Sumner (2011) or those directly set by stimulus location or from another parameter. White boxes indicate free parameters used to capture new data.

| Name | Description | Bompas & Sumner (2011) | Exp 1 |  | Exp 2 |  | Exp3 |  |
| --- | --- | --- | --- | --- | --- | --- | --- | --- |
|  |  |  | IGN | STOP | IGN | STOP | IGN | STOP |
| $Ecc_{Dist}$ | Distractor eccentricity in SC (mm) | -1.76 | 0 | | | | | |
| $Ecc_{Targ}$ | Target eccentricity in SC (mm) | 1.76 | 2.25 | | | | | |
| $\delta_{vis}$ | Visual delay (ms) | 50 | Individual $T_{Op-IGNORE} - 15$<br>fixed across IGNORE and STOP (see Table 4) | | | | | |
| $\delta_{endo}$ | Endogenous delay (ms) | 75 | $\delta_{vis} + 25$ | | | | | |
| $\delta_{holding}$ | Holding period after fixation offset | 0 | Fitted to individual NO-SIGNAL RT distributions, separately for IGNORE and STOP contexts (see Table S2) | | | | Fitted to individual NO-SIGNAL RT distributions (see Table S2) | |
| $a_{endo-fix}$ | Amplitude of endogenous inputs at fixation | 10 | | | | | | |
| $a_{endo-targ}$ | Amplitude of endogenous inputs to target | 14 | | | | | | |
| $F$ | Failure rate | 0 | Individual STOP error rate at SOA = 0 (see Table 4) | | | | | |

### 5.1. Visual delay

In previous work, we have explained why and illustrated how sensory conduction times for visual signals can be directly estimated from dip onset time (Bompas et al., 2017; Bompas & Sumner, 2011). Providing *δ_vis_* and *δ_out_* are constant across trials and a large number of trials are available, *T_0_* = *SOA* + *δ_vis_* + *δ_out_*. This is because the earliest effect a visual stimulus can have on a saccade RT distribution represents the case where a distractor signal arrives (*SOA* + *δ_vis_* after target onset) at the selection system just before the decision threshold is reached by the target activity (*δ_out_* before the response would have occurred). Using 20 ms for output time (consistent with previous work) and a similar smoothing as in observed data (which anticipates dips by 5 ms), we set *δ_vis_* for each individual to *T_0p_* - 15 ms. In the data presented in section 4, we observed that *T_0_* hardly changes across contexts and experiments, despite the large differences in mean RT observed across blocks. This confirms that *δ_vis_* does not contribute to pro-active slowing, consistent with the automatic nature of exogenous signals in DINASAUR, and consistent with the behavior from visuo-movement neurons. We therefore based all our modeling on the *T_0p_* from the IGNORE condition only (e.g. Figure 7B top panel). For simplicity, we assume that *δ_vis_* is equal for targets and distractors (this is a simplification as they have different eccentricity and sizes).

### 5.2. Baseline parameters from NO-SIGNAL trials

The next step was to adjust as few parameters as possible to fit the model to the baseline conditions (Figure 7B red lines). When IGNORE and STOP instructions are delivered in different blocks, such as in Experiment 2, participants adjust their behavior overall, leading to slower RT in the STOP block irrespective of signal presence (see section 3.1). This proactive slowing is present to a smaller degree in Experiment 1 when stop trials were always present but differed in frequency between blocks. In Experiment 3, the baseline RTs were the same in the two instructions and also suggested pro-active slowing (the mean RT were close to the STOP blocks in Experiments 1 and 2). To allow a fair test of the model’s ability to generalize from distraction to countermanding, it is essential to fit the different latency distributions of the baseline conditions. Critically, we adjusted the model parameters solely based on NO-SIGNAL trials.

It is common to assume that pro-active slowing would be best captured by an increase in initiation threshold (Bogacz, Wagenmakers, Forstmann, & Nieuwenhuis, 2010; Forstmann et al., 2010; Ratcliff & McKoon, 2008). This is indeed what simple models such as the independent race model would suggest (Heitz & Schall, 2012; Verbruggen & Logan, 2009). However, this assumption is not confirmed by electrophysiological recordings from monkeys (Heitz & Schall, 2012; Pouget et al., 2011; Reppert et al., 2018). Specifically, in SC neurons, firing rates some 10-20 ms prior to saccade initiation (i.e. the threshold) were the same under a speed and accuracy conditions (Reppert et al., 2018). Similarly, no change in threshold was observed after stop-signal trials, another way in which pro-active slowing has been investigated (Pouget et al., 2011). In FEF neurons, firing rates prior to saccades were actually *lower* in the accuracy condition compared with the speed condition, in direct contradiction to the increase in threshold suggested by the fit from the independent race model on concurrent behavioral data from these monkeys (Heitz & Schall, 2012). In contrast, both SC and FEF visuo-motor neurons consistently showed modulation in baseline firing rate (before target onset), as well as delayed target selection time (Reppert et al., 2018), i.e. the time at which the activity diverges depending on whether the receptive field of the neuron contains a task-relevant or task-irrelevant stimulus. Last, in SC visuo-motor neurons, changes from fast to accurate instructions were not accompanied by a modulation in visual gain (Reppert et al., 2018, i.e. the intensity of the visual response to stimulus onset that would be identical for targets and distractors).

In the DINASAUR model, baseline firing is directly related to the strength of endogenous fixation drive during the fixation period (*a_endo_fix_*), while delayed target selection can be produced by increasing the delay (*δ_endo_*) or reducing the strength (*a_endo_targ_*) of the endogenous drive to the target. Indeed, stronger fixation drive in the stop task would, via lateral inhibition, reduce baseline firing rate in all peripheral nodes, making it more difficult to produce fast but possibly erroneous saccades to the target (in line with Wong-Lin et al., 2010). Similarly, RT to the target largely relies on endogenous drives, since exogenous drives are most of the time insufficient to reach the threshold. Furthermore, visual gain directly maps to the strength of visual signals (*a_exo_*), which was therefore kept fixed across instructions (as was the delay of exogenous signals, see section 5.1), consistent with their automatic nature. As for all the other parameters in the model, in the absence of specific hypothesis for why they may differ 1) across instructions or 2) compared with previous work, we refrained from altering these, providing the strictest test of our model.

*δ_endo_* was not originally conceived as a free parameter in DINASAUR, as it is by default tied to *δ_vis_* (see Section 2.7). We therefore first varied *a_endo_fix_* and *a_endo_targ_* systematically to search for the most suitable pair for each individual NO-SIGNAL distribution. *a_endo_fix_* was varied from 5 to 60 in steps of 1, while *a_endo_targ_* was varied from 10 to 20 in steps of 0.5. In Experiments 1 and 2, this was done separately for the IGNORE and STOP contexts, as these were acquired in separate blocks and were therefore open to strategic adjustments. In Experiment 3, the two tasks were interleaved, producing only one NO-SIGNAL distribution per participant. Individual distributions were each compared with 1000 trials simulated using each parameter combination, scaled to match the available trial number from each participant (Figure 7 illustrates the outcome of this procedure on one example participant). All fits were based on minimizing the *X^2^* distance between observed and simulated NO-SIGNAL RT distributions. To increase the sensitivity to the exact shape of the whole RT distribution, we used a fixed bin size (3.33 ms, the same as for the distributions throughout the article with the same smoothing) rather than a small number of quantiles. This choice led us to use the mean over two complementary estimates, *X^2^_data_* and *X^2^_model_*. Within each bin, *X^2^_data_ = (N_data_ – N_model_)^2^ / N_data_*, with N denoting the number of saccades for which RT fell within this bin, while, *X^2^_model_ = (N_data_ – N_model_)^2^ / N_model_*. This mean estimate therefore penalizes simulations producing saccades in bins where none are observed, as well as simulations failing to produce saccades in bins where some are observed. The overall *X^2^* was the sum of the *X^2^* over all the bins where *N_data_*(or *N_model_*) was at least 1. Although this approach was the most intuitive to us, we note that using alternative fitting approaches (*X^2^_data,_ X^2^_model_* or *X^2^_data_* on 10 quantiles) actually made little difference to the fit and no difference to our conclusion.

Although most fits were satisfying, four (out of 20) remained poor and these were specifically misrepresenting distributions with very long mean RT but comparatively small standard deviations, in conditions subject to large pro-active slowing (Exp 3, or the STOP context of Exp 1 and 2). Increasing *a_endo-fix_* mainly prevents short responses, while decreasing *a_endo-targ_* increases most RT, but to the cost of also increasing variability. Instead, the pattern of data was suggestive of participants strategically waiting before disengaging from fixation, presumably to avoid errors at short SOAs. Such holding period has been proposed before as a mechanism for pro-active slowing, from the behavior and neuronal activity of monkeys performing a saccadic stop task (Lo et al., 2009). We therefore added this new parameter to DINASAUR and re-ran the fits, allowing the fixation-holding period to vary from 0 to 50 ms in steps of 10, while *a_endo_fix_* and *a_endo_targ_* were varied in steps of 5 and 1 respectively. This improved the fit in the four cases mentioned above (X^2^ were now below 500) and produced marginal improvement in another three cases. The best set of parameters across the two fitting procedures was then chosen (see Table S2). Note that the focus of this article is not on modeling strategic pro-active slowing, but to identify the common automatic components between countermanding and SI. We therefore made no attempts to formally compare models and test whether adding a free parameter to the model was worth it. Rather, our purpose is limited to using neurophysiologically plausible adjustments in order to provide a satisfying fit to our NO-SIGNAL distributions, so these parameters can be taken forward for testing the generalization to the SIGNAL conditions.

### 5.3. Generalization to SIGNAL-IGNORE and SIGNAL-STOP trials

Crucially, once the adjustments to the NO-SIGNAL trials were made to account for proactive slowing, we could test the ability of the model to generalize to the SIGNAL conditions for each SOA (note that our parameters were never allowed to differ between SOAs). The model was able to produce the expected dips from the IGNORE condition across all SOAs, as illustrated for one example participant on Figure 7, and from pooled data across participants and SOA on Figure 10 (see Table S3 for X^2^ distances between observed and simulated data). Unsurprisingly, the generalization from *T_0p_*(used to fit the visual delay) to each SOA was excellent (Figure S2), confirming the validity of the approach. Simulated dips were often sharper than observed one (the recovery was quicker), but note that we did not attempt to fit the strength and transience of the automatic input associated with the signal onset (these were inherited from previous work using peripheral small black distractors). These may well be different in the current design (larger central white circles) but this was not the focus here.

**Figure 10.**
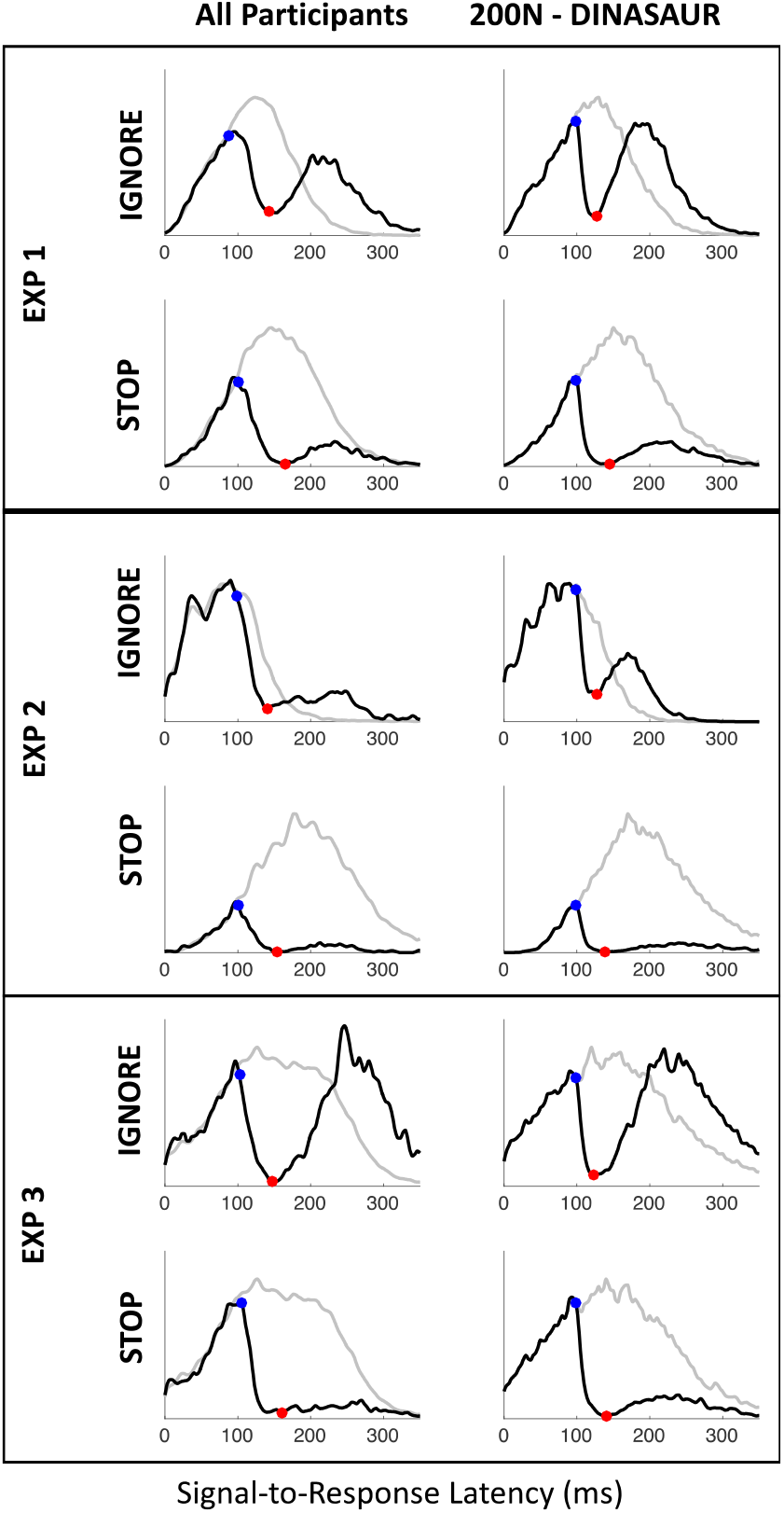
Distributions of RT locked on signal onset, pooled across all SOAs and observers, along with simulations using 200N-DINASAUR model pooled in the same way. Same conventions as Figure 8.

The critical step was then to test how well behavior on SIGNAL-STOP trials could be predicted from our model under the following assumptions: i) the automatic exogenous activation should be identical to the IGNORE context (in both amplitude and delay); ii) the endogenous response to the signal is not free, its timing is fully constrained by the automatic signal delay (*δ_endo_* = *δ_vis_* + 25 ms) and its amplitude is inherited from previous work (*a_endo-fix-post_* = 10). We assess the model against both the shape of the RT distributions (Figures 7, 10 and Table S3), as well as dip onset times across SOAs (Figure S2) and typical measures related to the stop-signal task (Figure 9 and S3).

At first, we did not introduce any new parameter between the IGNORE and STOP contexts (dotted red lines on Figure 9A-B). Like in Figure 3, this first attempt was able to produce the overall pattern of the stop condition, producing very similar effects as the state of the art model for saccadic countermanding, Blocked Input 2.0. However, similar to Blocked Input models (Figure 3C) but in contrast to observed data, there were no “late” errors: the small recovery from the dip observed in all the participants in Experiment 1 and 3, and one participant in Experiment 2 was absent in the model. As a result, the inhibition function (the proportion of failed stops as a function of SOA, dotted lines on Figure 9A) was systematically underestimated. Second, again similar to Blocked Input 2.0, DINASAUR predicted stop-signal reaction time (SSRT) to remain constant across SOAs (dotted lines on Figure 9B), in contrast to observed data showing a consistent decrease as a function of SOA in both experiments (diamonds on Figure 9B).

Within the framework of the independent race model, a decrease in measured SSRT can be explained by assuming the *true* SSRT varies across trials, and that varying the SOA leads to differently sampling this underlying distribution (Logan & Cowan, 1984). Since at short SOAs, most responses are successfully inhibited, the estimated SSRT is close to the true mean of SSRT. However, at long SOAs, only the shortest SSRT lead to successful inhibition, therefore leading to a systematic underestimation of the mean SSRT. This interpretation works mathematically, but from our perspective, a simpler mechanistic interpretation seems to be in terms of failure to trigger the stop instruction, which would occur on some proportion of trials (Band et al., 2003; Emilio Salinas & Stanford, 2013).

In the framework of the DINASAUR model, the same idea (variability of stop drive across trials) can be implemented in a simple way by adding a “failure” (or inattention) parameter, i.e. a random proportion of trials where the STOP instruction is forgotten and in which the system behaves exactly as in IGNORE trials. This refinement is conceptually similar to that proposed in Hanes & Carpenter (1999), but is now explicitly linked to the ignore condition, which the system defaults to when the instruction to stop occasionally fails to be implemented. It is also well in line with similar suggestions made in the more cognitive domain and using manual responses (Band et al., 2003; Matzke et al., 2017; Skippen et al., 2019). In DINASAUR, top-down drives are either on or off while, realistically, their strength and delay probably vary across trials. One could envisage that, on some trials, the blocking occurs but is incomplete or occurs too late, leading to the saccade being triggered anyway. These cases would be difficult to distinguish from a complete failure to apply the instruction to stop, and are therefore also captured by our failure parameter.

It is essential to note that this failure rate parameter does not account for any unexplained differences between ignore and stop behavior. Rather it accounts for more-than-expected similarity by simply putting the model back into ignore mode for a proportion of trials. This adjustment allowed late recovery from stop-signals, which improved the match to the inhibition function (continuous lines on Figure 9A), allowing more errors to be made by the model, bringing it more in line with human participants. On these occasions when the stop instruction is not applied, everything happens as if the instruction was to ignore the signal. The saccade recovers after a pause, creating a long tail just like in the standard saccadic inhibition paradigm, only much reduced in size since this failure affects only a minority of trials. This failure parameter was set to be equal to the percentage of errors on SIGNAL-STOP trials at the shortest SOA (50 ms), which ranged from 3 to 31% across individuals (see Table 4). The rationale is that, at such short SOA, all trials should be inhibited successfully if the instruction were applied correctly. Although these values may seem high, we note that our participants were all novices on the stop task, in contrast to monkeys or humans from labs where this task is intensely investigated. Furthermore, this number is in line with estimates from recent work, also involving novices, suggesting an average value of 17%, though using a different set-up (Skippen et al., 2019). Crucially, although this failure parameter is constant across SOA (like all other parameters), the proportion of saccades eligible for recovery decreases as SOA increases, and this now makes our model successfully capture the dependency of SSRT on SOAs (continuous lines on Figure 9B).

All model simulations on Figure 10, S2 and S3 use this failure parameter. Figure 10 illustrates the ability for the model to capture all aspects of the STOP data (see Table S3 for individual X^2^ measures). Figure S2 and S3 illustrate the excellent prediction of *T_0-STOP_*, error rate and SSRT at each SOA. *T_0-STOP_* generalized equally well as *T_0-IGNORE_*, as assessed by the sum of the X^2^ distance between observed simulated values (23 for IGNORE and 17 for STOP, non-significantly different).

## 6. Discussion

### How do brains halt action plans? Intertwined influences of automatic and top-down processes

The thesis in the present article is that the functional outcome of top-down control occurs initially via automatic indiscriminate mechanisms, which are followed by goal directed processes in the traditional view. When halting an action plan following new information in the world, the first process is a rapid automatic interference from the new sensory signal itself – which occurs regardless of the goal to halt. This indiscriminate interference has dynamics arising from the transient nature of rapid visual signals (such as the magnocellular pathway) and lateral inhibition in motor decision areas. It results in slowing down the process that leads to action, temporarily interrupting it. The endogenous command to alter the on-going action plan can then piggy-back on the already-unfolding automatic interruption. This account offers a simple interpretation for a wealth of data showing how “low-level” factors affect our ability to stop (Armstrong & Munoz, 2003; Asrress & Carpenter, 2001; Boucher, Stuphorn, et al., 2007; Cabel et al., 2000; Hanes & Carpenter, 1999; Hanes et al., 1998; Hanes & Schall, 1995; Ito et al., 2003; Morein-Zamir & Kingstone, 2006; Paré & Hanes, 2003; Stuphorn et al., 2000). It also allows quantitative predictions for many other factors, which have been shown to automatically interfere with speeded responses but may not have been studied in the context of countermanding (see “Empirical predictions and future directions” below).

This is not to say that rapid interference is entirely goalless in the broader sense: our brains may allow this interference to happen because it is helpful on average. In other words, natural selection seems to have preserved some apparently very basic – and probably phylogenetically old – processes that allow new and often irrelevant sensory information to rapidly travel to motor decision areas and influence action choices within 100 ms. We envisage this as one of the initial building blocks for how flexible behavior becomes possible as brains develop additional pathways that are more selective but slower. Further, while in simple visual scenes (such as in these experiments) all new stimuli may provide indiscriminate interference, in complex everyday scenes the degree of rapid interruption is likely to be modulated by relevance to on-going tasks (‘attention’ or ‘task-set’). It is known that spatial attention modulates sensory signals from the earliest stages of processing (as early as the lateral geniculate nucleus for visual signals, O’Connor, Fukui, Pinsk, & Kastner, 2002). Similarly, sub-conscious motor priming is highly conditional on task-set (current task goals; i.e. whether the priming stimuli have a current motor mapping or not), suggesting automatic flows of activity through the brain show *conditional* automaticity (see Kunde et al., 2003 for an in-depth discussion on this topic) – and hence are not entirely goal-free. This dependency of automatic drives on task-set is also illustrated in pro-active control (Verbruggen, Best, et al., 2014; Verbruggen, Stevens, & Chambers, 2014). Therefore, although the present article develops the idea that top-down processes piggy back on automatic ones, we see it as complementary to the literature showing that automatic processes often piggy-back on top-down processes, pointing towards a close intertwining of automatic and volitional drives (Boy, Husain, & Sumner, 2010; Sumner & Husain, 2008).

Our conclusions are convergent with previous literature showing how task goals, such as stopping, can be influenced by invisible or task-irrelevant primes (see Verbruggen, Best, et al., 2014 for a review). Our viewpoint is also compatible with other recent theories of countermanding. Here we investigated the effect of visual stimuli on oculomotor control in humans, but our conceptualization is in line with other literatures describing animal behavior, such as freezing, as proposed in the Pause and Cancel model in rodents (Schmidt & Berke, 2017). Our conclusions are reminiscent of those from Bisset & Logan (2014) on selective stopping paradigms, where participants are asked to stop to some signals but ignore others within the same session. In this context, it has been suggested that participants use a Stop then Discriminate strategy, in which they stop indiscriminately whenever a signal occurs and restart only if the signal is an ignore signal. However, we portray the initial stage as slowing down rather than stopping, and as an automatic process rather than a strategy.

### Movement vs visuomovement neurons

Once we clearly conceptualize the first process in halting as transient automatic interference, we can clarify the alignment between recent models of countermanding and low-level mechanisms. The early process in previous countermanding models such as Blocked-Input 2.0 or in (Lo et al., 2009) was already conceptualized as stimulus driven with a short delay, although it was implemented as a sustained signal. An important implementation difference with DINASAUR relates to the distinction between visuomovement and movement neurons. DINASAUR units are simplified visuomovement SC neurons. As a result, they will show an automatic transient visual response, followed by a buildup of activity when the task requires it. In contrast, units in models such as Blocked Input 2.0 are thought to reflect FEF movement neurons. This means that they will not show the automatic visual response, but only the task-related accumulation. It has been argued that only movement (and not visuomovement) neurons reflect the accumulation of evidence that leads to saccadic decision (Ray et al., 2009). The fact that movement neurons (but not visuomovement neurons) showed activity profiles that matched those expected of GO units in a race model contributed to this assumption. Reciprocally, the presence of neurons with activity resembling the hypothetical GO units also contributed to legitimize the race model.

Counter to this prevailing view, it is precisely the visuomovement nature of DINASAUR units (their automatic transient response to visual stimuli as well as their strategic drives) that makes DINASAUR capture tasks it was not originally designed for – the saccadic inhibition and countermanding tasks – as well as several hallmarks of visuo-oculomotor behavior such as the gap effect (Bompas & Sumner, 2011, 2015; Trappenberg et al., 2001), and visuo-manual interference (Bompas et al., 2017). Similarly, our upgrade of Blocked Input 2.0 to Blocked Input 3.1 consisted precisely in turning units from movement neurons into visuomovement neurons. The fact that neurons exist that behave in a similar way to units in our model is a necessary condition for this model to be “biologically plausible” but surely does not prove the model is right, nor that these neurons are precisely the ones “taking the decision”. Although it is essential to simplify complex behaviors and concepts into workable models, we keep in mind that this simplification makes all computational models intrinsically wrong. Ultimately, the proposed framework offers the opportunity to generate precise quantitative predictions, which can then be tested empirically (see “Empirical predictions and future directions” below). The endeavor here is not to “validate” one particular model or show it outperforms other models in specific tasks, but rather to employ a precise framework to bridge gaps across paradigms and literatures.

### Converging modeling approaches

As developed above, the crucial difference between the models lies in the transience and indiscriminate nature of the stimulus-driven signal. Apart from this, the delay times and other aspects of the model logic were similar. We inherited the logic of blocking input for the endogenous signal from the most comprehensive model of countermanding (Logan et al., 2015), but we inherited nearly all actual parameters from saccadic inhibition (either previous work or the baseline and ignore conditions here). Countermanding behavior then drops out of the model. The model’s activity dynamics are also consistent with monkey neurophysiological data (Boucher, Palmeri, et al., 2007; Hanes et al., 1998) – an important test-bed for previous models of countermanding (e.g. Logan et al., 2015).

To allow a match to every aspect of the data, we made two additions: a strategic fixation-holding period and a failure parameter to capture the occasional late errors. The first parameter was only introduced to improve the fit to the no-signal condition, in line with previous behavioral and neurological work in monkeys performing a saccadic countermanding task (Lo et al., 2009). The second parameter is needed not because the model did not sufficiently change its behavior between ignoring and countermanding, but because human behavior actually remains more similar across the conditions than the model predicts, as if they sometimes forget to countermand. Both parameters are new to DINASAUR, but their plausibility has been already well supported in the context of the stop-task (Band et al., 2003; Lo et al., 2009; Matzke et al., 2017; Skippen et al., 2019).

However, even without these post-hoc additions, the model was able to generate good predictions in a behavior it had never been constrained for. It is worth emphasizing how rare it is for psychological models to capture new behavior for which they were not designed without being fit directly with plenty of free parameters. This might have been even more challenging when crossing a conceptual boundary – such as from bottom-up interference to top-down control. However, our thesis is that this should not be considered a conceptual boundary. Situations requiring top-down control do not differ qualitatively from those that stimulate automatic interference and most of the same brain mechanisms are engaged in both situations. Moreover, although elegant parsimonious mathematical models designed to capture specific tasks may often struggle to generalize to other tasks (unless completely re-fit or parameters are added that change the model characteristics), generalization is more natural in more complex models conceived to mimic a biological system. Of course, more parameters mean more flexibility, should one allow all these parameters to vary freely. That is why our approach is the opposite: we keep most parameters fixed and only allow very few parameters to vary in a highly constrained, hypothesis-driven manner. The ability of such models to generalize to new behaviors, combined with a clear logic for what should be allowed to differ and what should be fixed, are great strengths, which, in our eyes, outweigh the loss in parsimony and mathematical elegance.

Although our account bears conceptual resemblance to other recently proposed models of stopping, there remain important implementation differences. Specifically, the Pause then Cancel model (Schmidt & Berke, 2017) relies on an unspecific increase in the action initiation threshold following the stop signal event. Similarly, in Aron & Wessel (2017), a temporary slowing can be triggered in response to any unexpected events. Both accounts suggest this indiscriminate response could be mediated by the Basal Ganglia (BG), which has inhibitory connections with the SC. In contrast, DINASAUR mimics topologic relations between the visual field and the direction of saccades, as is commonly seen in SC buildup neurons during visually-driven saccades. This difference in implementation could arise from a focus on different animal species and therefore on different types of action (ballistic head movements in rodents and saccades in monkeys). However, both BG and SC are involved in both actions in both species and it is therefore likely that both should contribute to stopping behaviors, the former as a general freezing mechanism and the later as a more spatially specific mechanism able to resolve competition across multiple stimuli in the visual field. Although simplified and limited, the spatial extent of the DINASAUR model allows us to test future predictions related to the spatial specificity of stopping behavior (see “Empirical predictions and future directions” below). Future research investigating this spatial specificity could cast light on the relative contribution of the Basal Ganglia (possibly less spatially specific) and the Superior Colliculus into saccade countermanding.

### Model simplifications

Our approach to minimize the number of free parameters in the model led to three main simplifying assumptions (beyond the fact that all models are simpler than neuronal processes). First, most parameters not of direct interest here were inherited from previous work, including the spatial profile of excitation and inhibition, the spatial extent of excitation from visual onsets and the temporal profile of exogenous signals. These parameters were based on monkey neurophysiology (Trappenberg et al., 2001), and appear sufficient for simulating currently existing human datasets (present and past, see Bompas & Sumner 2011).

Second, we assumed visual onsets triggered the same automatic response (delay and amplitude), irrespective of their eccentricity. Visual eccentricity is known to decrease sensitivity and acuity, which could, in the model, mean weaker and functionally slower signals. On the other hand, oculomotor behavior is, by definition, designed to orient towards peripheral stimuli, which may therefore be prioritized in oculomotor planning. To fully compare conduction delays (*T_0_*) across eccentricity is beyond the current data, but a proxy can be obtained from the very quickest saccades that are not guesses (i.e. the shortest-latency in which there are more correct than error saccades). In our data, this latency was 106 ms, and occurred in the condition expected to have lowest engagement with fixation: the IGNORE condition of Experiment 2. This suggests that *T_0_* for these peripheral stimuli would have been approximately 100ms, allowing for a minimum amount of decision time and a slight pooling delay needed to detect above chance performance. This proxy estimate is similar to our estimate for *T_0_* at fixation (98ms), and suggests our simplifying assumption of equal latency was sufficiently sound.

Third, we assumed that, apart from the strategic holding period adopted by some participants, all endogenous delays were equal, including fixation release, saccade planning and blocking. This assumption followed from our endeavor to predict the pattern of countermanding behavior from lower-level oculomotor behaviors without separately fitting a special inhibitory or blocking mechanism. It is off course possible that these delays may differ slightly, in a way that relates interestingly to task-set or individual differences.

### How fast are top-down commands?

The traditional purpose of countermanding research is to understand and measure how rapidly a top-down signal can overturn an action plan, quantified by the SSRT. One of the implications of the close relationship between bottom-up and top-down processes is that the effective speed of top down signals depends on bottom-up factors. This conclusion is actually consistent with a wealth of research showing that SSRT depends on the exact experimental condition, and we provide here a general framework for explaining this. In this framework, all top-down drives, including stopping, are about translating sensory information into task-related action outcomes. Therefore, the speed of top-down drives will heavily depend on non-decision time, i.e. sensory conduction time and motor output time, which will depend on the nature of sensory information and action modalities under investigation.

This being said, within the context of one task, one can usefully discuss the speed of top-down drives associated with a given sensory signal, action domain and instruction set. One key implication of conceptualizing the first phase of halting as automatic is that the truly endogenous signal does not have to be so rapid. This point echoes that of the Pause-then-Cancel theory of basal ganglia mechanisms (Schmidt & Berke, 2017), where it is argued that a fast pause mechanism is followed by a cancel process that extends well beyond the traditional SSRT, and therefore we may have been looking in the wrong temporal window for neural evidence of such mechanisms.

However, in our present results, the latency remains relatively short for the top-down signals. SSRT is normally estimated as between 100 and 150 ms in humans for saccades (Campbell, Chambers, Allen, Hedge, & Sumner, 2017; Hanes & Carpenter, 1999). In our model there are two relevant input delays: visual and endogenous delay. These are respectively 83 ms for the transient automatic signal to start interfering with saccade build-up activity, and 108 ms for endogenous support to switch back to fixation. For comparison with SSRT, we need to add motor output time, in this case 20 ms, because SSRT is a measure of the time needed between a stop signal and when a response would otherwise have *occurred*, not just the time before the inhibition signal reaches motor maps. This gives us 103 and 128 ms. One could therefore conclude that the new conceptualization overall supports previous estimates for the window of inhibitory signals.

Importantly though, neither of these two delays in DINASAUR can be interpreted as reflecting the timing of inhibition *per se*. Indeed, the first is the delay of automatic excitatory signals. When these automatic signals project to fixation neurons, they have an inhibitory effect on the plan to move the eyes to the target, but only indirectly, via lateral inhibition. The second only indexes the *start* of the endogenous switch, while the inhibition disrupting the link between the visual stimulus and the intention to saccade needs to be sustained throughout a long period to prevent saccades from recovering from the dip. Besides, the timing of this later drive is not specific to stopping, but is shared with all top-down drives in the model.

How stopping is conceptualized also impacts the conceptual ordering of go and stop command speed. As previously envisaged within the influential independent race model of countermanding, the go signal always comes first and stop commands always have to *catch up* to take effect. This would have misled many into thinking that stop commands are on average faster than go commands. In contrast, in Blocked Input 2.0, the stopping delay (*D_control_*) is larger (62 and 90 ms for Monkey A and C) than the delay for producing go saccades (*D_move_*, 44 and 47). In our model the two delays facilitating stops (83 and 108 ms) are identical to those producing go saccades to the target. How then is it possible for stimuli occurring *after* the target to trigger a majority of stops if the relevant delay parameters are equal to or longer than those driving go saccades?

The answer is that in an interactive model a go saccade only occurs after an accumulation process, which takes some amount of time after the signals start getting integrated into this process. However as soon as a new signal, or a change in signal (e.g. one being turned off), reaches that process it can immediately change the accumulation, potentially stopping activity that was about to reach threshold doing so. In other words, go response latency depends on both the input delays and the accumulation time (plus output time), while inhibition speed depends mainly on the input delays (plus output time for behavioral evidence of inhibition). This distinction was of course known to previous researchers using interactive models. However, it does not appear to be widely discussed that stop processes can be successful and appear to ‘overtake’ go processes without there having to be neural mechanisms that are themselves more speedy for inhibition than for initiation of responses.

Although Boucher et al. (2007) stress that the stop signal is ‘late and potent’, while we have referred to rapid transient inhibition, this difference of language merely occurs because of different reference positions. This signal is rapid when compared with human saccade latency distributions, or to the later influences of top-down signals. But it is late in the sense that it accounts for most of measured SSRT. It is potent in both models, in the sense that as soon as the signals reach the neural maps, lateral inhibition creates a strong impediment to saccade planning and has an almost immediately measurable effect in the reduction of saccade likelihood.

### The importance of sensory pathway dynamics in motor decision

Our findings confirm the suspicions of Cabel et al. (2000) and Morein-Zamir & Kingstone (2006) that stimulus properties (such as salience) often influence task performance by engaging both automatic and top-down processes. This warns us not to assume that well-known behavioral effects in tasks associated with higher-level processes always measure mechanisms at that level. The model framework we use provides a natural explanation for the influence of stimulus properties, which dictates both the timing and amplitude of the automatic dips (Bompas & Sumner, 2011; Reingold & Stampe, 2002). Likewise there are known differences between SSRT arising from visual and auditory stop signals (Armstrong & Munoz, 2003; Boucher, Stuphorn, et al., 2007; Cabel et al., 2000; Morein-Zamir & Kingstone, 2006), which might traditionally be ascribed to the time needed to detect the stop signal before issuing the countermand, but in the model would also be captured by different dip size and delay. Auditory signals also produce dips, which happen sooner than following visual stimuli, although these have only been studied on microsaccades (Rolfs, Kliegl, & Engbert, 2008).

Even changes to response modality – saccadic vs manual – which might not intuitively be associated with different stimulus-driven effects, in fact do affect the balance of drive from different sensory pathways (Bompas & Sumner, 2008), and thus the delay and amplitude of stimulus-driven activity (see Bompas et al., 2017 for discussion and demonstration of the presence of dips in the manual modality). This could be part of the reason why SSRT differs between modalities (Boucher, Stuphorn, et al., 2007) and possibly also why saccadic and manual SSRT are differentially susceptible to influences such as alcohol (Campbell et al., 2017).

Some task designs (e.g. manual responses with low-salience stop signals) may entail a sufficiently small automatic effect that explicitly including it in models would not alter conclusions in any important way. Indeed, the standard horse-race model of countermanding has been applied successfully to very many studies. However, we should not assume this will be the case for all manual designs, and we advocate paying close attention to the nature of stimuli and the non-linear activity they produce. For instance, it is possible for masked no-go or stop stimuli to slow down responses and slightly increase the rate of missed responses (van Gaal, Lamme, Fahrenfort, & Ridderinkhof, 2011), suggesting those invisible stimuli can partially prime activity, even if this would not manifest obviously in latency distributions under ignore instruction (for example if there was no strong lateral inhibition at the stage this priming reaches). Therefore top-down inhibition may partially piggyback on automatic processes even when it is difficult for us to detect this behaviorally.

### Non-independence of go and stop processes

The fact that mean RT for failed stops tends to be shorter than mean RT for correct saccades has long been interpreted as evidence that the go and stop processes are independent. This concept, known as contextual independence, states that the finishing time of the go process is unaffected by the presence of the stop signal (see Bissett & Logan, 2014 for a recent explanation). The (flawed) logic underlying this conclusion is that, if the stop signal interferes with the action plan triggered initially by the go signal, and therefore slows it down, we would expect the RT in stop trials to be longer, not shorter, than in go trials. However, longer RT would actually only be expected for the fraction of trials for which the stop signal 1) reaches the competition before the saccade plan has reached threshold and 2) fails to prevent the saccade plan from reaching threshold. These saccades will be those with RT longer than T_0_. In contrast, the bulk of the failed stop RT distribution is populated by trials where the saccade plan was quick enough to escape all influence from the stop signal (RT < T_0_), and therefore have short unaltered RTs. For these trials, the stop and go signals did remain independent because the stop signal was still in sensory transmission. On the other hand, when the stop signal does reach the integration stage and interferes with the go process, a large proportion of these slowed action plans never reach fruition; they are successfully stopped. Therefore, they do not appear in the calculation of mean latency.

In other words, the fact that mean RT for failed stops tends to be shorter than mean RT for correct saccades should be interpreted to mean that the majority of escaping saccades (failed stops) were those for which non-decision time (the period of signal independence) dominated their overall latency. It does not mean the entire processes are independent. Indeed, Lo et al.’s model, Blocked Input and DINASAUR all have in common that they do *not* adhere to this independence concept (fixation and move nodes are mutually inhibitory), and yet the mean RT of simulated failed stops also tend to be shorter than the mean no-signal RT. This demonstrates that this behavioral pattern is not a strong test for contextual independence. It clearly isn’t a sufficient condition.

More generally, we stress that our understanding of these phenomena emerged from direct comparisons between the shapes of full distributions of no-signal and failed stop RT (or spatial properties), rather than relying on summary statistics such as mean RT of failed stops and accuracy, which can hide underlying patterns. Conceptually similar difficulties for mean RT can occur in any paradigm in which some portion of the RT distribution in one condition does not show up in another condition (e.g. when errors occur, these trials remove themselves from correct RT distributions, and the missing correct RTs will often be biased to one end of the distribution).

Note that the same reasoning holds when comparing the landing position or peak velocity of saccades between go and failed stop trials, as attempted in Hanes & Schall (1995): only a fraction of failed-stops would be expected to be hypometric while all the others will be identical, making any difference difficult to observe unless one can be directed by a model to examine the latency bins where hypometria is expected. The saccades most affected by the interaction process are successfully stopped and removed from the calculations.

Previous work using saccades with visual (Gulberti, Arndt, & Colonius, 2014; Ozyurt, Colonius, & Arndt, 2003) and tactile (Akerfelt et al., 2006) stop signals show violations of the independent race predictions, suggesting interaction between go and stop processes (Colonius & Diederich, 2018). In contrast, it has been claimed that the idea of independence of the go and stop activity had been validated in neuronal recordings in FEF (Hanes et al., 1998) and SC (Paré & Hanes, 2003), because there was no difference in saccade-related activity in failed stops and correct trials when RT < SSRT + SOA, and no peak velocity or eccentricity difference in the saccades made (these would be behavioral consequences of any difference in SC activity). However, we now show that this way of selecting trials is very similar to RT < *δ_vis_* + SOA, when no influence from the signal is yet measurable (see Lo et al., 2009 for a similar logic). Figure S8 in Appendix offers a clear demonstration of this. In all models the stop and go signals remain independent while the stop signal is in sensory transmission before it reaches the integration process. The proportion of failed stops that occur during this time are expected to show contextual independence.

### What does SSRT reflect?

Simulations using published parameters for Blocked Input 2.0 produced *T_0_* around 60 ms and this value maps well onto the sum of excitatory input delay (47) and output time (10), just like in DINASAUR. Using the standard integration calculation for SSRT (but see Skippen et al., 2019), the same simulations produce SSRT estimates of 73 ms for Monkey A and 93 ms for Monkey C (similar to observed SSRT, 71 and 94 ms), irrespective of SOA. These values map approximately onto the sum of *D_control_* + *δ_out_* for Monkey A (62 + 10), less clearly so for Monkey C (90 + 10). However, the proximity may be coincidental, since SSRT is also clearly influenced by other parameters in the model (*D_move_* and *D_fix_*), though not in straightforward ways.

Our SSRT estimates systematically decrease with increasing SOA, as previously noted in the countermanding literature. This trend suggests that SSRT does not directly reflect the timing of some unique underlying parameters of the sensorimotor system (as already noted by Emilio Salinas & Stanford, 2013’, using a much simpler model), as these would not be expected to vary with SOA. Linking saccade countermanding to saccadic inhibition and modeling both tasks with DINASAUR offers a quantitative explanation for this. The SSRT measure ignores the RT of failed inhibition, and therefore treats late errors equivalently to early errors. Given that dips are never so sharp that the distribution falls to zero straight after dip onset, there are always failed stops beyond dip onset. Their number contributes to SSRT and is influenced by nearly all parameters in the two models we considered. Therefore, SSRT is always higher than *T_0_*, and is a compound measure of all parameters that contribute to the success, or not, of stopping, rather than a reflection of inhibitory delay alone.

SSRT is typically interpreted as the time required for an action plan to be cancelled. However, within our current framework, the saccade plan is never truly cancelled. Rather, increased activity within the fixation system and interrupted support to this saccade plan may reduce movement activity sufficiently to make recovery very unlikely. Yet, the probabilistic nature of this mechanism means that it is possible that a saccade recovers. Relatedly, there is no room in this framework for the concept of cancel time, which has been proposed to index the efficiency of the cancelling process (Boucher, Palmeri, et al., 2007; Lo et al., 2009; Logan et al., 2015). The cancel time is defined as the difference between SSRT and the time at which neuronal activity starts to diverge between failed-stop trials and latency matched no-signal trials (diverging time on Figure 6). In contrast, the diverging time itself is entirely related to dip onset (in DINASAUR as well as Blocked Input and most other models relying on an interactive accumulation to threshold), since it is equal to SOA + sensory delay, i.e. T_0_ – motor output time. Therefore, although the diverging time directly maps onto one parameter in a range of models, the SSRT does not, and therefore the difference between SSRT and diverging time doesn’t either.

Many researchers use SSRT to measure individual differences in stopping ability. In light of the above, individual differences in SSRT could reflect variability within multiple aspects of visuomotor decisions (including properties of exogenous signals), rather than a unique construct or even a compound construct mainly indexing top-down control. For instance, if in some clinical condition, the sensory conduction delay associated to the go signal (eg. a peripheral stimulus) was increased *more than* the delay associated with the stop signal (eg. a central stimulus), this would be equivalent to effectively reducing the SOA, resulting in an increase in SSRT, even though none of the endogenous aspects are affected. Whether this multi-dependence of a key measure is practically beneficial or detrimental for researchers depends ultimately on how correlated low level and high-level aspects are within the population, which we do not know for now (see section below for future directions). We can only speculate that the answer will presumably depend on the specific design chosen to investigate these individual differences (stimuli, action modality and instructions), begging caution when drawing conclusions from experiments using different set-ups. The model supplies a conceptually useful distinction that is merged in SSRT: whether better “ability to stop” translates into quicker/stronger application of top-down control (a longer-lasting dip as top down control takes over from the automatic inhibition) or more consistent blocking behavior across trials (fewer late errors/failures). This is well in line with very recent work, suggesting correcting SSRT estimates for trigger failure improves correlation with impulsivity trait (Skippen et al., 2019).

### Empirical predictions and future directions

To further test the model framework, one can use quantitative predictions arising from changing the bottom-up parameters. Many “low-level” factors, such as signal contrast, chromaticity or position in the visual field, have been shown to modulate the automatic delaying of saccades. Using previous quantitative estimates for how these factors precisely influence the delay and strength of exogenous signals, quantitative predictions for stopping behavior can be easily derived from DINASAUR. For instance, we have previously described how increasing the signal’s contrast equates, in DINASAUR, with increasing the strength and decreasing the delay of exogenous signals (Bompas & Sumner, 2009, 2011). Similarly, our modeling suggests that some chromatic signals (“S-cone stimuli”) are delayed by 25 ms compared to achromatic signals (Bompas & Sumner, 2008, 2011). Previous research has also shown that stimuli presented in the temporal hemifield (such as left visual hemifield when viewed with the left eye), interfere more with saccade latency compared with nasal stimuli (right visual hemifield when viewed with the left eye) (Walker, Mannan, Maurer, Pambakian, & Kennard, 2000). From this we can make quantitative predictions for how much harder it should be to stop in response to low contrast, nasal or chromatic stimuli, compared to high contrast, temporal or chromatic stimuli. Conversely, the present data show that dip onset, which we use to constrain the delay of exogenous inputs, can also be estimated from the stop signal task. This means that existing stop task datasets with sufficient trials could be reanalyzed using the present framework in order to investigate automatic inhibition.

The current DINASAUR model is only 1D and its spatial aspects are still largely under-constrained (we have not allowed them to vary; they were inspired by recordings in monkeys but were never systematically tested against human behavior). Nevertheless, the fact that it possesses such spatial layout contrasts with most decision models (which possess typically 2 nodes), and offers the possibility to investigate the effect of spatial attributes of signals and targets, such as size and location. For instance, DINASAUR correctly accounts for the fact that interference can be triggered by visual stimuli appearing at any location in the visual field but it also predicts that the interference should be modulated by where the stop signal specifically appears, in relation to the fixation and the saccade target. Previous research has shown that, in the stop task, signals appearing at the same location as the target were less potent than contralateral signals (Ozyurt et al., 2003). This is consistent with our previous work showing such stimuli fail to induce any saccadic inhibition (Bompas & Sumner, 2011), possibly due to the existence of a refractory period preventing two bursts of visual activity to occur close in time at the same location. It is therefore possible that these signals do not produce any automatic interference and act purely via top-down signals, providing an interesting design for isolating top-down factors.

Another prediction from our framework is that factors mainly influencing top-down drives or the ability to apply these consistently (such as task switching, dual tasking, workload etc) should affect primarily the ability to stop saccades from recovering after the dips, but not dip onsets. More generally, the influences of clinical conditions, medications or other individual differences (age, personality traits etc) may well manifest as a combination of automatic and top-down drives differences. Therefore, disentangling the early (automatic dip) and late (blocking) stages in saccade countermanding, as the DINASAUR framework offers, should help in revealing more specifically those higher-level factors researchers are often primarily interested in.

So far, we have assumed that the delay of endogenous drives, including blocking, is fully determined by the delay of exogenous drives, being simply 25 ms longer. This choice was driven by parsimony and justified by the fact that all our signals are visual and had similar properties. Endogenous signals are simply viewed as further-processed versions of exogenous signals. However, it would be interesting to validate this assumption empirically, by measuring to what extent the exogenous delay (indexed by dip onset time) correlates with the endogenous delay (further constrained by the shape of the go distribution), across participants or across conditions. Within the context of individual differences, this would also allow us to test whether the blocking has indeed the same delay as the endogenous signals driving the saccade to the target. Similarly, it could be tested whether endogenous timing is indeed the largest source of variability across people, as is commonly assumed in the countermanding literature.

## 7. Conclusions

To conclude, the theoretical, simulation and experimental work presented here suggests that automatic stimulus-driven interference accounts for much of the characteristic behavior in countermanding tasks, in contrast to the traditional and widespread idea that these tasks primarily index higher level cognitive control. This highlights the importance of stimulus-driven effects in paradigms generally associated with higher cognition. More generally, we hope to help shift the traditional separation of automatic and voluntary processes towards a more integrated understanding of how automatic and voluntary control work together, alongside parallel endeavors to untangle the mysteriously intelligent control homunculus into the emergent activity of an army of idiots.

## 8. Author note

The authors acknowledge funding from the School of Psychology at Cardiff University, Alcohol Research UK (RS 12/01; AEC), the Economic and Social Research Council (ES/K002325/1; AB and PS) and the Wellcome Trust (104943/Z/14/Z). Funding sources had no involvement at any stage of the reported research. We are grateful to Geoffrey Megardon for help with the analysis and to Craig Hedge and Christopher D. Chambers for useful comments on the manuscript. The original submission of this article was published on BIORXIV/2019/535104. Some of the data were presented at the European Conference on Eye Movements in 2017.

## Appendix

**Table S1-A.** Individual mean reaction times (standard deviations) and SSRT in ms in Exp 1.

| Exp1 | RT – IGNORE |  |  |  | RT – STOP |  |  |  | SSRT |  |  |
| --- | --- | --- | --- | --- | --- | --- | --- | --- | --- | --- | --- |
| SOA | No-Signal | 50 | 83 | 133 | No-Signal | 50 | 83 | 133 | 50 | 83 | 133 |
| P1 | 207<br>(29) | 279<br>(67) | 245<br>(67) | 209<br>(40) | 241<br>(36) | 289<br>(67) | 247<br>(65) | 236<br>(49) | 153 | 137 | 127 |
| P2 | 210<br>(42) | 302<br>(83) | 262<br>(92) | 226<br>(79) | 239<br>(58) | 268<br>(117) | 215<br>(82) | 207<br>(46) | 133 | 120 | 120 |
| P3 | 196<br>(40) | 261<br>(95) | 220<br>(80) | 206<br>(70) | 242<br>(64) | 275<br>(89) | 224<br>(76) | 221<br>(47) | 153 | 143 | 127 |
| P4 | 231<br>(27) | 272<br>(39) | 275<br>(51) | 254<br>(59) | 261<br>(34) | 299<br>(72) | 295<br>(67) | 287<br>(74) | 180 | 157 | 127 |

**Table S1-B.**
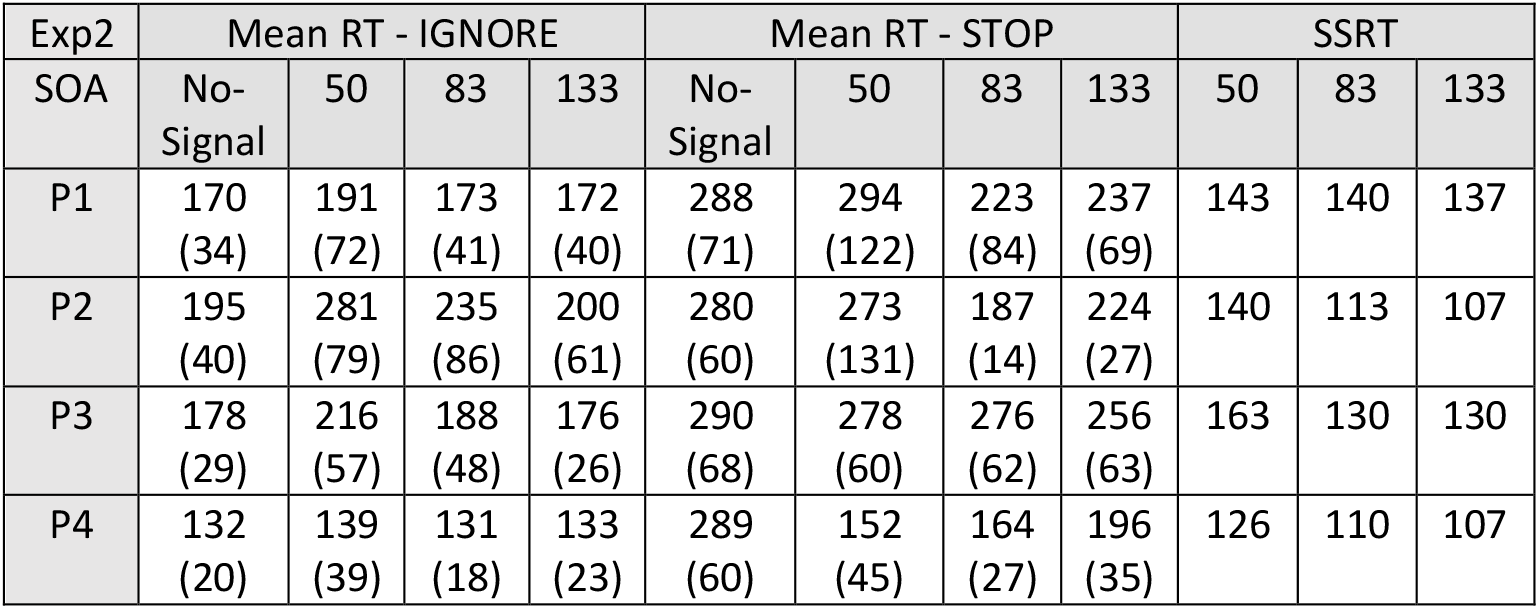
Same as S1-A for Experiment 2. Mean RT appearing in grey should be treated with caution as they were calculated on less than 50 trials.

**Table S1-C.** Same as S1-B.

| Exp3 | Mean RT No-signal | Mean RT - IGNORE |  |  |  | Mean RT - STOP |  |  |  | SSRT |  |  |  |
| --- | --- | --- | --- | --- | --- | --- | --- | --- | --- | --- | --- | --- | --- |
| SOA |  | 50 | 83 | 133 | 183 | 50 | 83 | 133 | 183 | 50 | 83 | 133 | 183 |
| P1 | 223<br>(63) | 273<br>(89) | 268<br>(105) | 247<br>(98) | 238<br>(85) | 180<br>(68) | 177<br>(54) | 194<br>(52) | 213<br>(59) | 130 | 120 | 117 | 123 |
| P2 | 247<br>(74) | 351<br>(110) | 317<br>(125) | 277<br>(112) | 253<br>(96) | 296<br>(129) | 209<br>(82) | 213<br>(77) | 226<br>(53) | 140 | 130 | 113 | 153 |
| P3 | 293<br>(38) | 345<br>(50) | 379<br>(47) | 395<br>(71) | 351<br>(108) | 356<br>(58) | 371<br>(56) | 320<br>(88) | 287<br>(47) | 190 | 167 | 127 | 130 |
| P4 | 284<br>(40) | 346<br>(55) | 358<br>(54) | 356<br>(78) | 300<br>(67) | 349<br>(65) | 330<br>(60) | 283<br>(68) | 276<br>(38) | 186 | 167 | 127 | 133 |

**Figure S1a.**
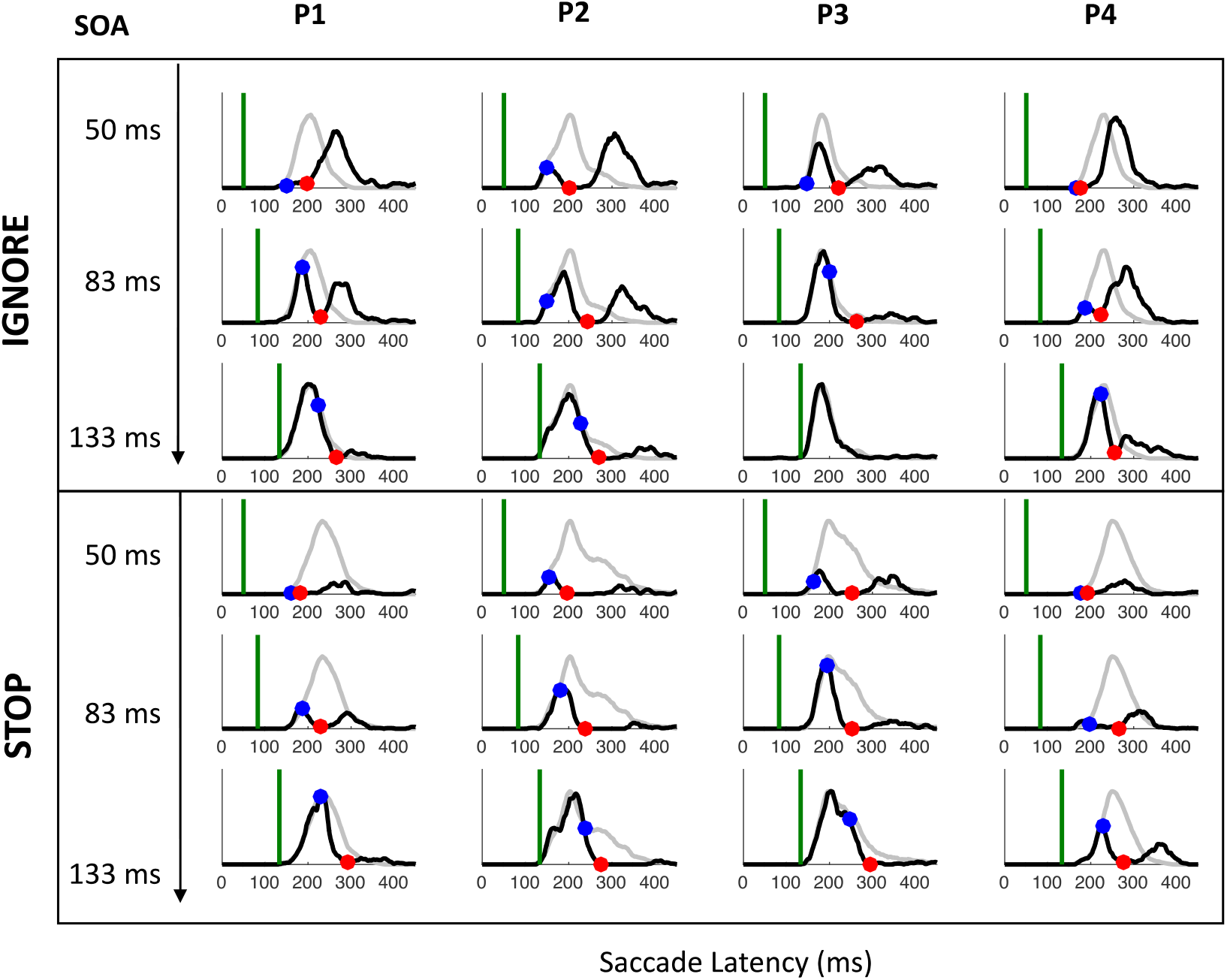
Latency distributions for each participant (columns) and SOA (rows) in the IGNORE and STOP contexts of Experiment 1. Green lines indicate the signal onset. Grey lines indicate distributions in which no signal was presented. Black lines indicate distributions of trials in which a signal occurred. Blue dots indicate the dip onset (i.e. where the distributions diverge, not necessarily where one takes a down-turn); red dots show dip maximum.

**Figure S1b.**
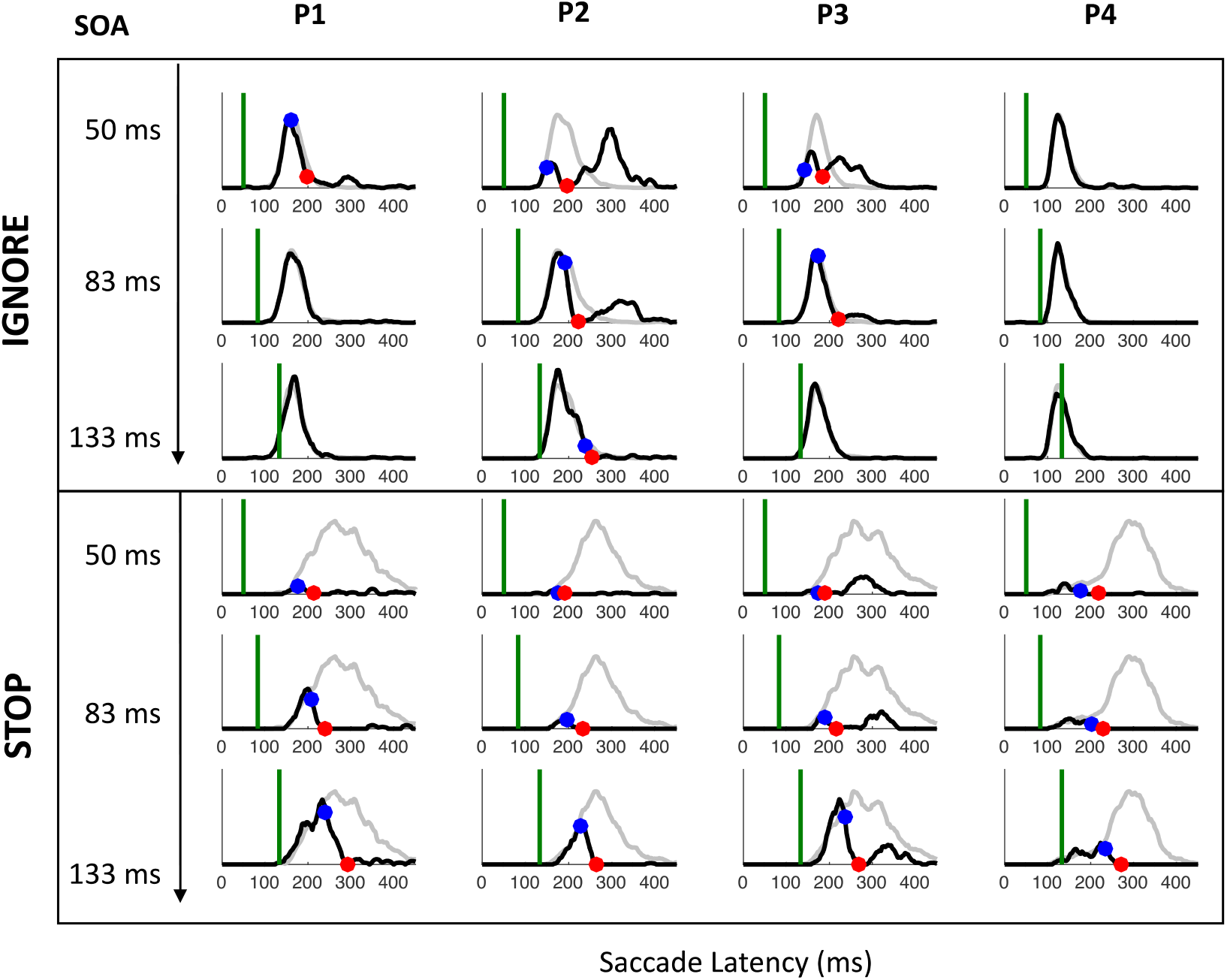
Same as S1a for Experiment 2. As expected, strategic adjustments across conditions were particularly large in Experiment 2 (where the two contexts were kept fully separated) and meant the visual signal often arrived too late to have much effect, especially for the fastest participants (P1 and P4). Nevertheless, when dips were observed in both contexts, Experiment 2 confirmed the results from Experiment 1.

**Figure S1c.**
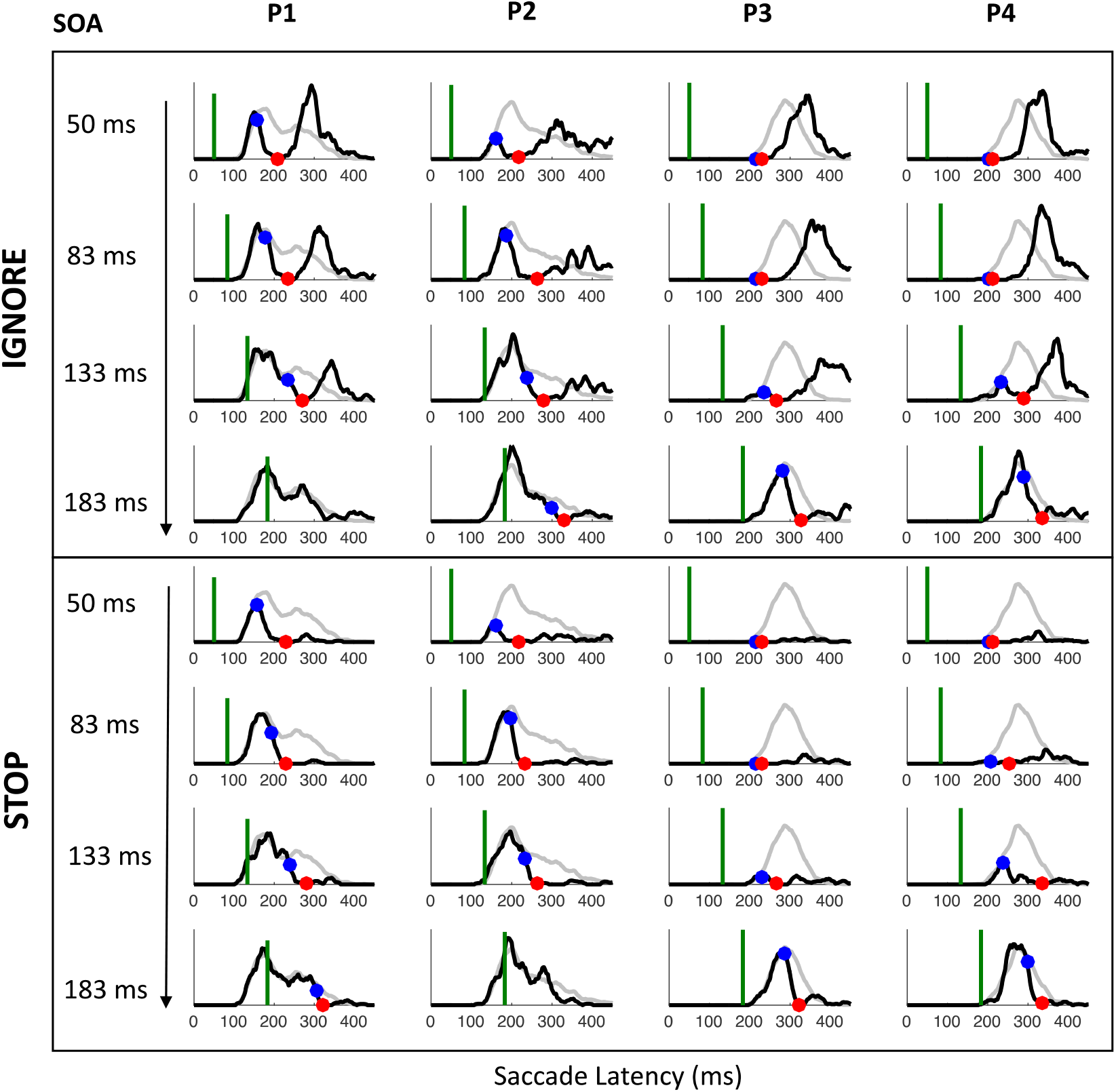
Same as S1a for Experiment 3 after pooling data from white and dark signals.

**Figure S2.**
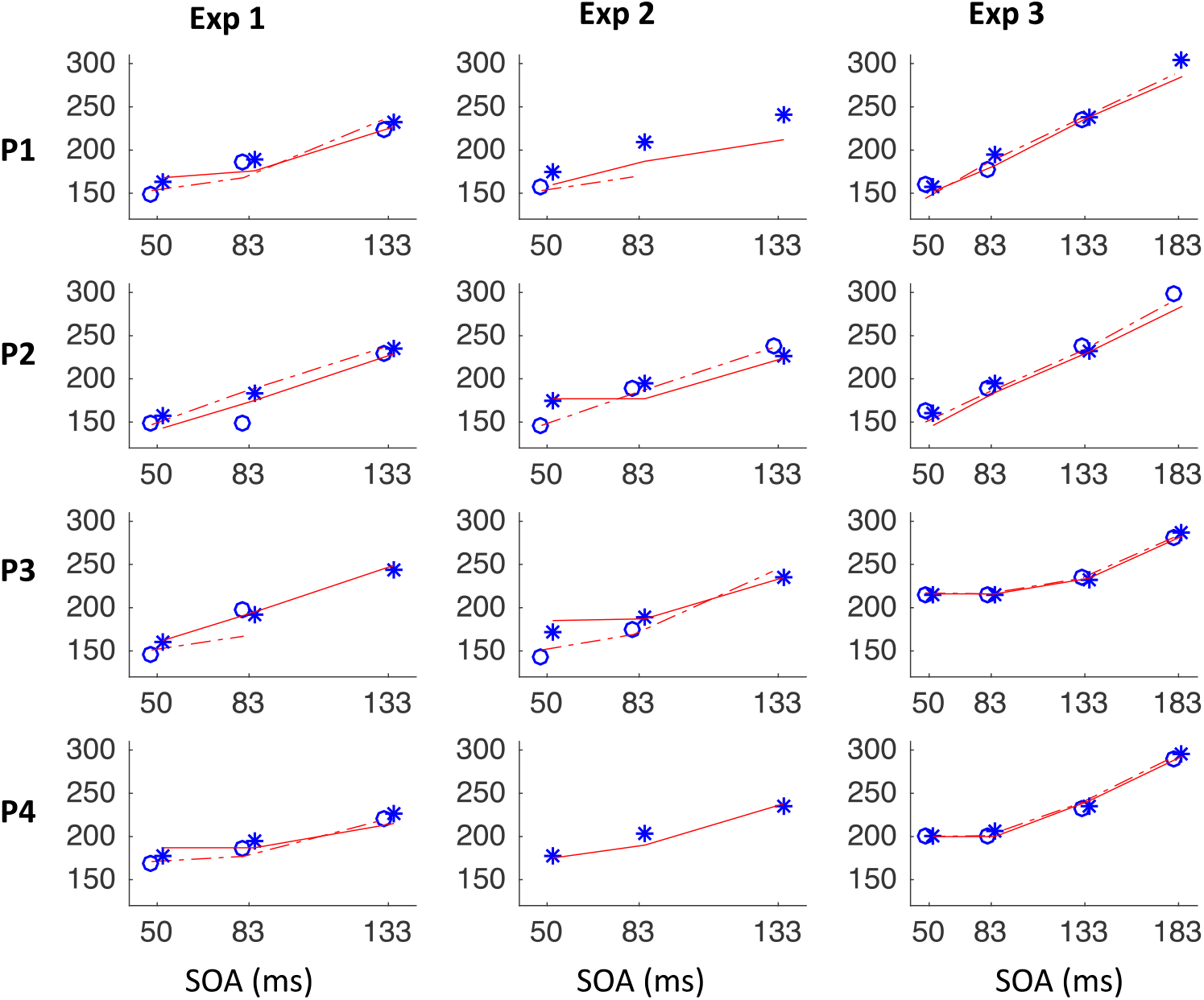
Individual T_0_ at each SOA in the IGNORE (circles) and STOP instructions (stars) along with simulated T_0_ using the IGNORE (dashed lines) and STOP (continuous line) parameters from Table S2. Missing points and lines indicate cases when the observed or simulated data did not show dips. Even though we only use T_0p_ from the IGNORE condition to constrain the model, note how well the model generalizes to each SOA and across instructions.

**Figure S3a.**
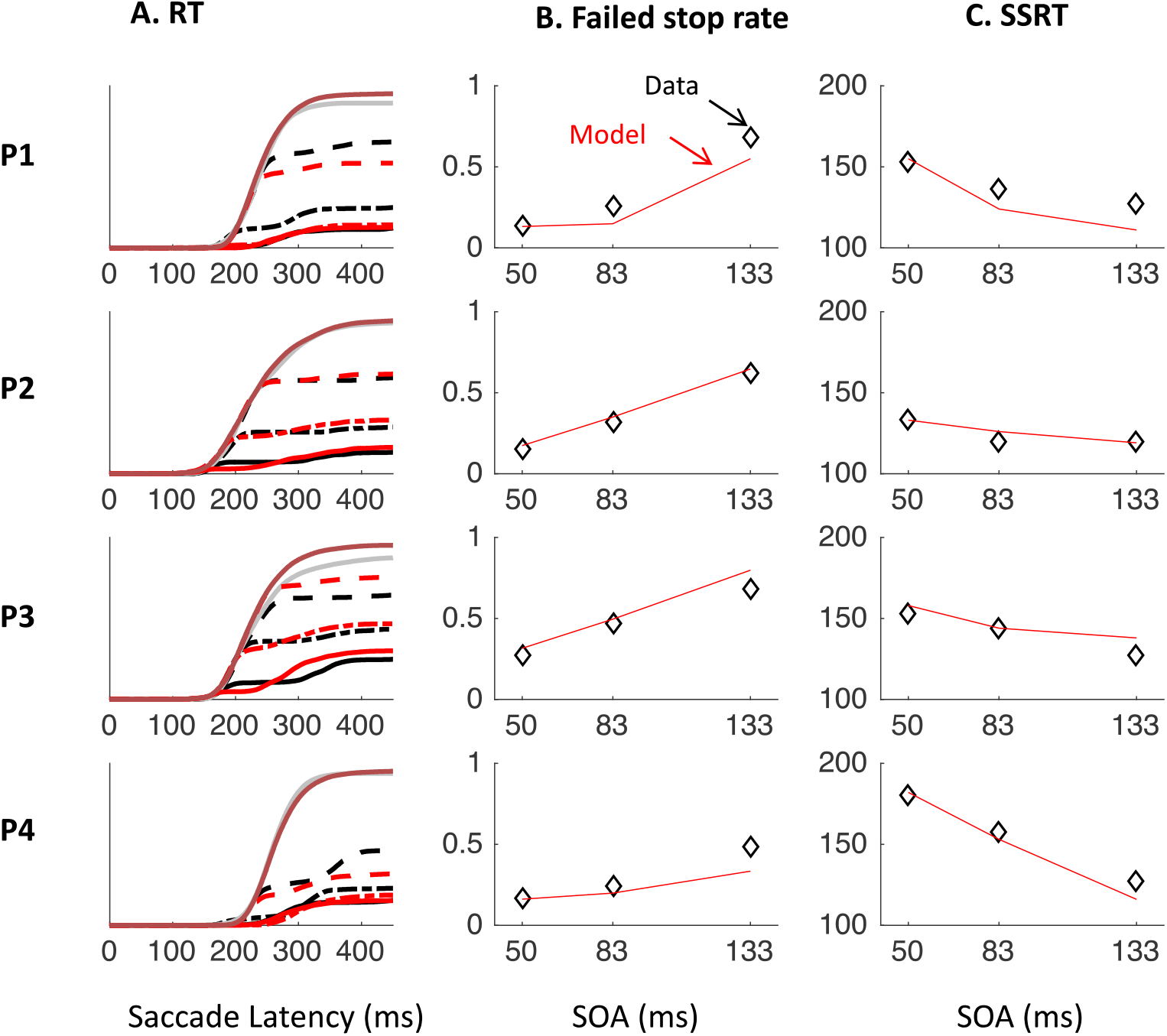
Observed data in the STOP task (grey and black) along with model simulations (red). **A.** Cumulative distribution of NO-SIGNAL RT (grey and dark red) and SIGNAL RT for SOAs 50, 83 and 133 (continuous, semi-dashed and dashed black lines for observed data and bright red lines for model). **B-C.** Inhibition function and SSRT for observed (diamonds) and simulated (lines) STOP-SIGNAL data.

**Figure S3b.**
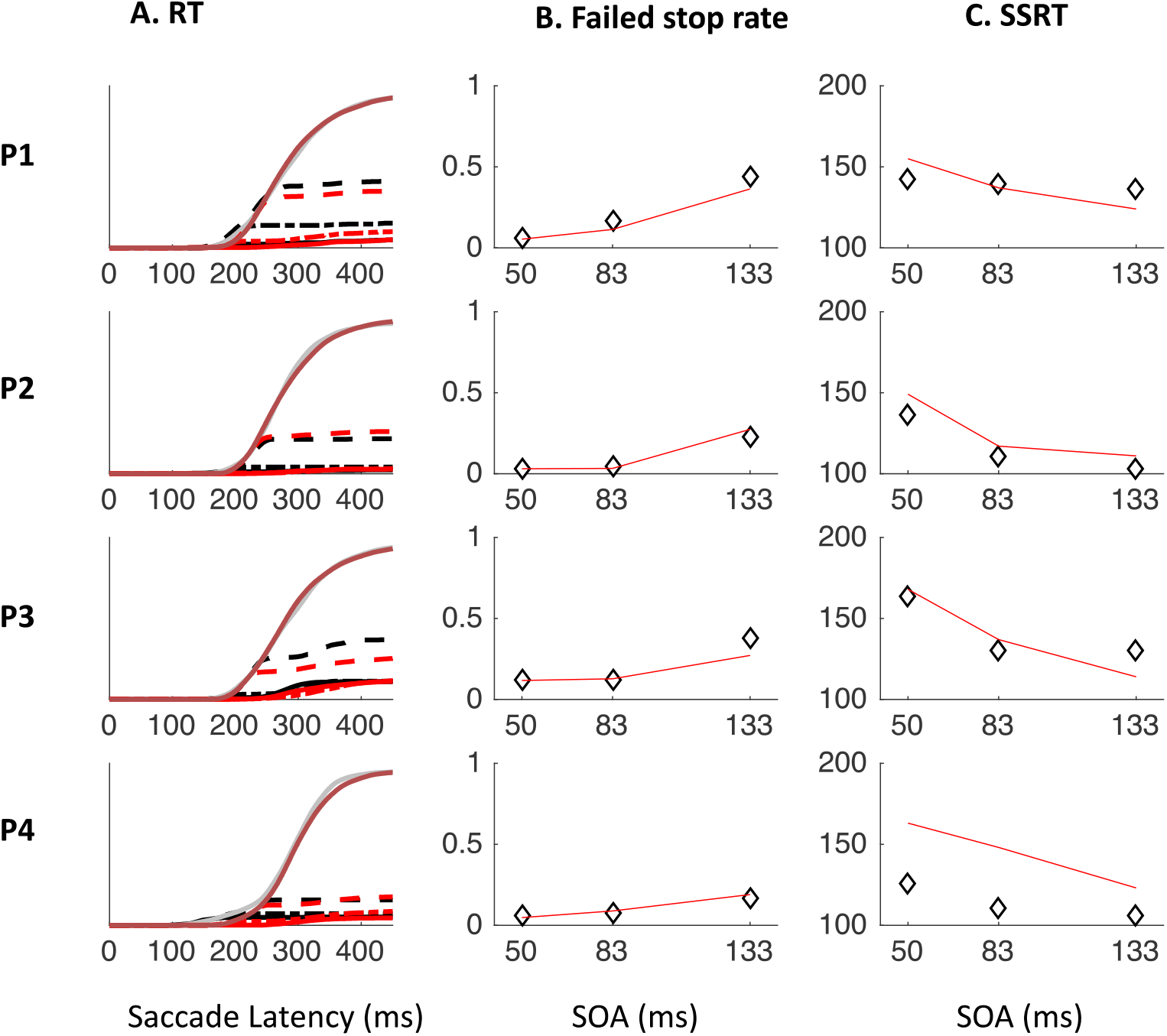
Same as S3a for Experiment 2.

**Figure S3c.**
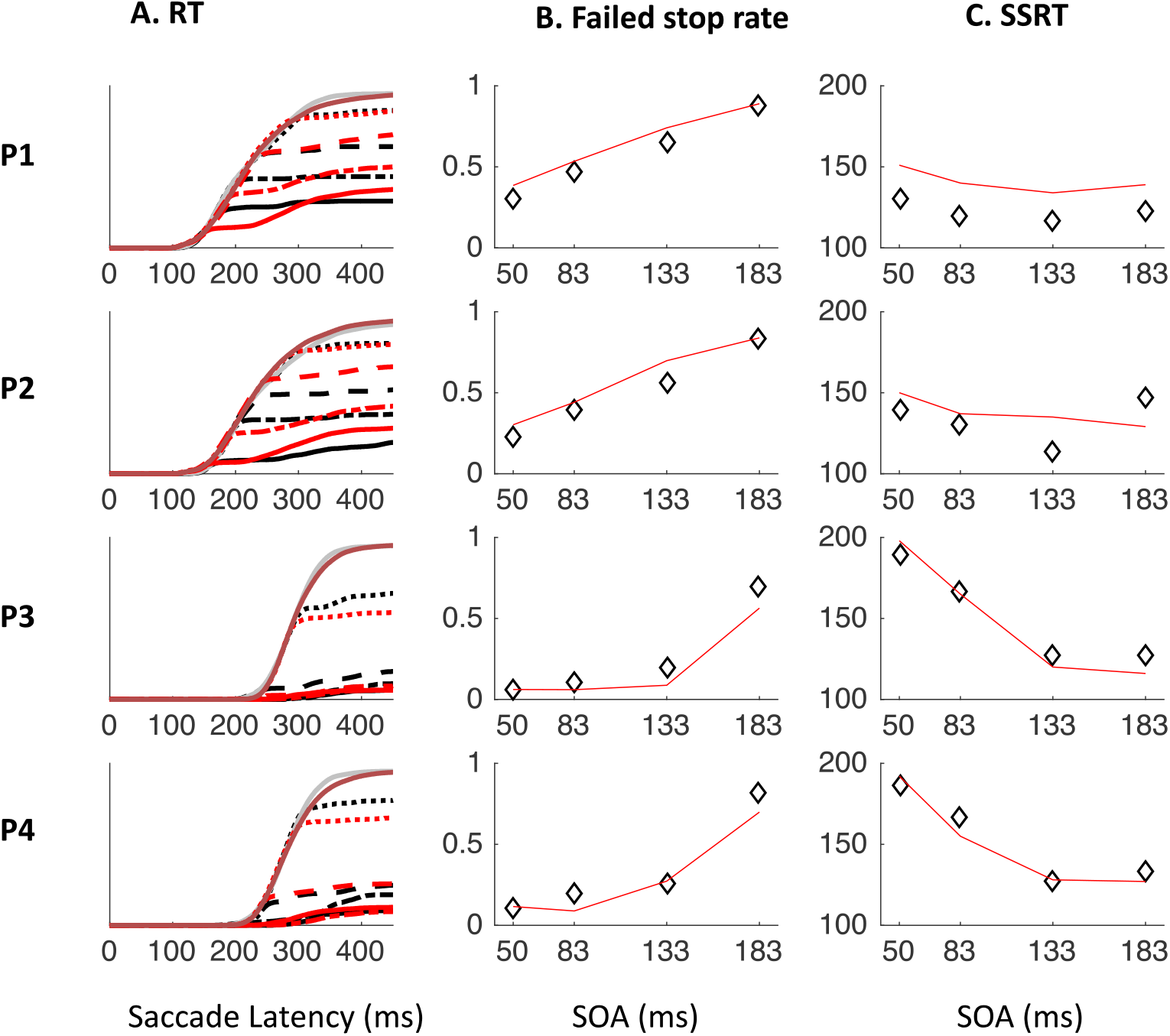
Same as S3a for Experiment 3.

**Table S2.** Parameter estimates for each individual in each NO-SIGNAL condition and goodness of fit (X^2^distance between observed and fitted data). a_endo-fix_ and a_endo-targ_ are the strength of endogenous inputs to the fixation and target nodes respectively. δ_holding_ is the duration of the strategic “holding period”. I1 and S1 correspond to NO-SIGNAL trials during the IGNORE and STOP blocks of Experiment 1. I2 and S2 correspond to NO-SIGNAL trials during the IGNORE and STOP blocks of Experiment 2. IS3 correspond to the NO-SIGNAL trials of Experiment 3.

| | $a_{\text{endo-fix}}$ | | | | $\delta_{\text{holding}}$ | | | | $a_{\text{endo-targ}}$ | | | | $\chi^2$ | | | |
| --- | --- | --- | --- | --- | --- | --- | --- | --- | --- | --- | --- | --- | --- | --- | --- | --- |
|  | P1 | P2 | P3 | P4 | P1 | P2 | P3 | P4 | P1 | P2 | P3 | P4 | P1 | P2 | P3 | P4 |
| I1 | 26 | 15 | 15 | 25 | 0 | 0 | 0 | 20 | 16.5 | 14.5 | 19.5 | 17 | 98 | 103 | 198 | 108 |
| S1 | 30 | 15 | 15 | 25 | 10 | 0 | 0 | 30 | 15 | 13 | 14 | 15 | 152 | 170 | 144 | 133 |
| I2 | 10 | 15 | 20 | 5 | 0 | 0 | 0 | 0 | 19.5 | 16 | 19.5 | 20 | 205 | 94 | 41 | 638 |
| S2 | 24 | 25 | 60 | 25 | 0 | 10 | 0 | 50 | 12.5 | 13 | 12.5 | 14 | 226 | 170 | 204 | 494 |
| IS3 | 12 | 13 | 30 | 35 | 0 | 0 | 50 | 20 | 12.5 | 12.5 | 15 | 14 | 519 | 249 | 275 | 191 |

**Table S3.** Accuracy of the predicted RT distribution for each SIGNAL condition, as measured by the X^2^ between predicted and observed individual RT distributions.

|  | P1 |  |  |  | P2 |  |  |  | P3 |  |  |  | P4 |  |  |  |
| --- | --- | --- | --- | --- | --- | --- | --- | --- | --- | --- | --- | --- | --- | --- | --- | --- |
|  | 50 | 83 | 133 | 183 | 50 | 83 | 133 | 183 | 50 | 83 | 133 | 183 | 50 | 83 | 133 | 183 |
| I1 | 1483 | 4223 | 505 |  | 7196 | 2620 | 770 |  | 15504 | 1145 | 358 |  | 369 | 819 | 993 |  |
| S1 | 195 | 697 | 1000 |  | 328 | 855 | 239 |  | 1246 | 359 | 430 |  | 111 | 267 | 846 |  |
| I2 | 1375 | 563 | 281 |  | 7180 | 5930 | 441 |  | 8081 | 1831 | 94 |  | 990 | 812 | 517 |  |
| S2 | 66 | 486 | 477 |  | 44 | 81 | 124 |  | 124 | 206 | 792 |  | 120 | 160 | 254 |  |
| I3 | 2290 | 1407 | 1009 | 950 | 2829 | 4192 | 1075 | 529 | 3907 | 7388 | 7138 | 2514 | 4081 | 2260 | 1152 | 1102 |
| S3 | 1498 | 753 | 471 | 857 | 459 | 871 | 295 | 518 | 86 | 180 | 431 | 795 | 211 | 484 | 283 | 580 |

**Figure S8.**
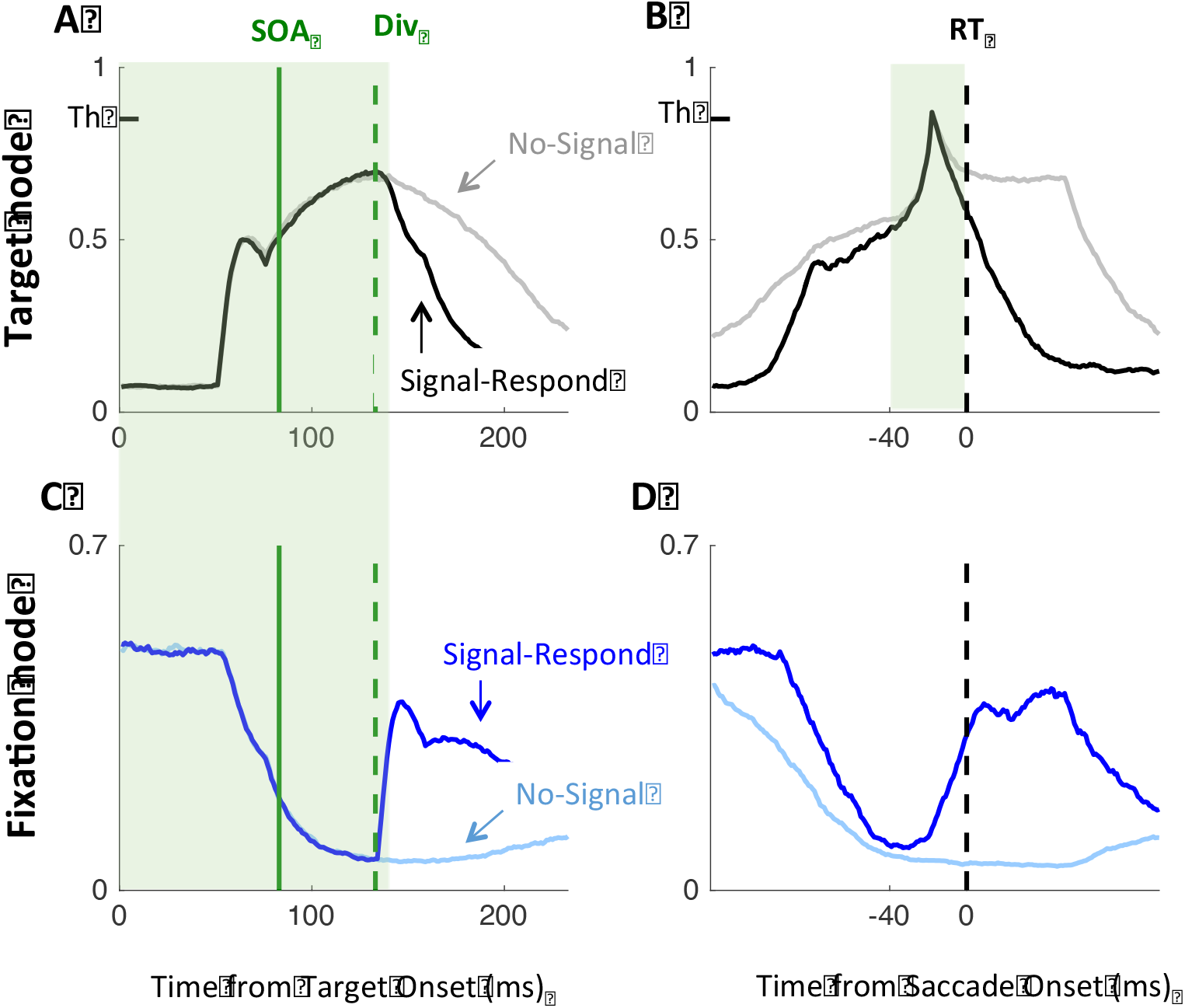
DINASAUR accounts for patterns in neural activity previously taken to imply independence of Go and Stop processes. **A&C.** Mean simulated activity during unsuccessful stop trials (signal-Respond) and latency matched No-Signal trials at SOA 83 ms, using the same convention as Figure 6 and matching Fig. 4 A&C in Boucher et al. (2007). **B.** Same data as in A but locked on saccade onset, following Fig. 3F in Paré & Hanes (2003). **D.** Same data as in C but locked on saccade onset (not shown in Paré & Hanes (2003), shown here for completion). Green shades indicate those time windows chosen in these two previous articles to illustrate the equality of neural activity between Signal-Respond and fast No-Signal trials. Clear differences are apparent outside these time windows.

## References

Akerfelt, A., Colonius, H., & Diederich, A. (2006). Visual-tactile saccadic inhibition. Exp Brain Res, 169(4), 554–563. doi: 10.1007/s00221-005-0168-x

Armstrong, I. T., & Munoz, D. P. (2003). Inhibitory control of eye movements during oculomotor countermanding in adults with attention-deficit hyperactivity disorder. Exp Brain Res, 152(4), 444–452. doi: 10.1007/s00221-003-1569-3

Asrress, K. N., & Carpenter, R. H. S. (2001). Saccadic countermanding: A comparison of central and peripheral stop signals. Vision Research, 41(20), 2645–2651. doi: 10.1016/S0042-6989(01)00107-9

Band, G. P. H., van der Molen, M. W., & Logan, G. D. (2003). Horse-race model simulations of the stop-signal procedure. Acta Psychologica, 112, 105–142. doi: 10.1016/S0001-6918(02)00079-3

Bissett, P. G., & Logan, G. D. (2014). Selective stopping? Maybe not. J Exp Psychol Gen, 143(1), 455–472. doi: 10.1037/a0032122

Bogacz, R., Wagenmakers, E. J., Forstmann, B. U., & Nieuwenhuis, S. (2010). The neural basis of the speed-accuracy tradeoff. Trends Neurosci, 33(1), 10–16. doi: 10.1016/j.tins.2009.09.002

Bompas, A., Hedge, C., & Sumner, P. (2017). Speeded saccadic and manual visuo-motor decisions: Distinct processes but same principles. Cogn Psychol, 94, 26–52. doi: 10.1016/j.cogpsych.2017.02.002

Bompas, A., & Sumner, P. (2008). Sensory sluggishness dissociates saccadic, manual, and perceptual responses: An S-cone study. Journal of Vision, 8(8), 1–13.

Bompas, A., & Sumner, P. (2009). Temporal dynamics of saccadic distraction. Journal of Vision, 9(9), 1–14.

Bompas, A., & Sumner, P. (2011). Saccadic inhibition reveals the timing of automatic and voluntary signals in the human brain. Journal of Neuroscience, 31(35), 12501–12512. doi: 10.1523/jneurosci.2234-11.2011

Bompas, A., & Sumner, P. (2015). Saccadic inhibition and the remote distractor effect: One mechanism or two? J Vis, 15(6), 1–8. doi: 10.1167/15.6.15

Boucher, L., Palmeri, T. J., Logan, G. D., & Schall, J. D. (2007). Inhibitory control in mind and brain: An interactive race model of countermanding Saccades. Psychological Review, 114(2), 376–397. doi: 10.1037/0033-295x.114.2.376

Boucher, L., Stuphorn, V., Logan, G. D., Schall, J. D., & Palmeri, T. J. (2007). Stopping eye and hand movements: Are the processes independent? Perception & Psychophysics, 69(5), 785–801.

Boy, F., Husain, M., & Sumner, P. (2010). Unconscious inhibition separates two forms of cognitive control. Proceedings of the National Academy of Sciences of the United States of America, 107(24), 11134–11139. doi: 10.1073/Pnas.1001925107

Buonocore, A., & McIntosh, R. D. (2008). Saccadic inhibition underlies the remote distractor effect. Experimental Brain Research, 191(1), 117–122.

Buonocore, A., & McIntosh, R. D. (2012). Modulation of saccadic inhibition by distractor size and location. Vision Research, 69, 32–41. doi: 10.1016/J.Visres.2012.07.010

Buonocore, A., & McIntosh, R. D. (2013). Attention modulates saccadic inhibition magnitude. Quarterly Journal of Experimental Psychology, 66(6), 1051–1059. doi: 10.1080/17470218.2013.797001

Buonocore, A., Purokayastha, S., & McIntosh, R. D. (2017). Saccade Reorienting Is Facilitated by Pausing the Oculomotor Program. Journal of Cognitive Neuroscience. doi: 10.1162/jocn_a_01179

Cabel, D. W. J., Armstrong, I. T., Reingold, E., & Munoz, D. P. (2000). Control of saccade initiation in a countermanding task using visual and auditory stop signals. Experimental Brain Research, 133(4), 431–441. doi: 10.1007/s002210000440

Camalier, C. R., Gotler, A., Murthy, A., Thompson, K. G., Logan, G. D., Palmeri, T. J., & Schall, J. D. (2007). Dynamics of saccade target selection: Race model analysis of double step and search step saccade production in human and macaque. Vision Research, 47(16), 2187–2211. doi: Doi 10.1016/J.Visres.2007.04.021

Campbell, A. E., Chambers, C. D., Allen, C. P. G., Hedge, C., & Sumner, P. (2017). Impairment of manual but not saccadic response inhibition following acute alcohol intoxication. Drug Alcohol Depend, 181, 242–254. doi: 10.1016/j.drugalcdep.2017.08.022

Colonius, H., & Diederich, A. (2018). Paradox resolved: stop signal race model with negative dependence. Psychological Review, In Press.

Cutsuridis, V., Smyrnis, N., Evdokmds, L., & Perantonis, S. (2007). A neural model of decision-making by the superior colicullus in an antisaccade task. Neural Networks, 20(6), 690–704.

de Jong, R., Coles, M. G. H., Logan, G. D., & Gratton, G. (1990). In search of the point of no return: the control of response processes. J Exp Psychol Hum Percept Perform, 16(1), 164–182. doi: 10.1037/0096-1523.16.1.164

Dorris, M. C., Olivier, E., & Munoz, D. P. (2007). Competitive integration of visual and preparatory signals in the superior colliculus during saccadic programming. Journal of Neuroscience, 27(19), 5053–5062.

Edelman, J. A., & Xu, K. Z. (2009). Inhibition of Voluntary Saccadic Eye Movement Commands by Abrupt Visual Onsets. Journal of Neurophysiology, 101(3), 1222–1234. doi: 10.1152/Jn.90708.2008

Elchlepp, H., Lavric, A., Chambers, C. D., & Verbruggen, F. (2016). Proactive inhibitory control: A general biasing account. Cogn Psychol, 86, 27–61. doi: 10.1016/j.cogpsych.2016.01.004

Forstmann, B. U., Anwander, A., Schafer, A., Neumann, J., Brown, S., Wagenmakers, E. J., … Turner, R. (2010). Cortico-striatal connections predict control over speed and accuracy in perceptual decision making. Proc Natl Acad Sci U S A, 107(36), 15916–15920. doi: 10.1073/pnas.1004932107

Gulberti, A., Arndt, P. a., & Colonius, H. (2014). Stopping eyes and hands: evidence for non-independence of stop and go processes and for a separation of central and peripheral inhibition. Frontiers in Human Neuroscience, 8(February), article 61. doi: 10.3389/fnhum.2014.00061

Hanes, D. P., & Carpenter, R. H. S. (1999). Countermanding saccades in humans. Vision Research, 39(16), 2777–2791. doi: 10.1016/S0042-6989(99)00011-5

Hanes, D. P., Patterson, W. F., & Schall, J. D. (1998). Role of frontal eye fields in countermanding saccades: Visual, movement, and fixation activity. Journal of Neurophysiology, 79(2), 817–834.

Hanes, D. P., & Schall, J. D. (1995). Countermanding Saccades in Macaque. Visual Neuroscience, 12(5), 929–937.

Harrison, J. J., Freeman, T. C. A., & Sumner, P. (2014). Saccade-like behavior in the fast-phases of optokinetic nystagmus: An illustration of the emergence of volitional actions from automatic reflexes. Journal of Experimental Psychology: General. doi: 10.1037/a0037021

Heitz, R. P., & Schall, J. D. (2012). Neural mechanisms of speed-accuracy tradeoff. Neuron, 76(3), 616–628. doi: 10.1016/j.neuron.2012.08.030

Ito, S., Stuphorn, V., Brown, J. W., & Schall, J. D. (2003). Performance monitoring by the anterior cingulate cortex during saccade countermanding. Science, 302(5642), 120–122.

Kopecz, K. (1995). Saccadic reaction times in gap/overlap paradigms: a model based on integration of intentional and visual information on neural, dynamic fields. Vision Res, 35(20), 2911–2925.

Kopecz, K., & Schoner, G. (1995). Saccadic motor planning by integrating visual information and pre-information on neural dynamic fields. Biol Cybern, 73(1), 49–60.

Kunde, W., Kiesel, A., & Hoffmann, J. (2003). Conscious control over the content of unconscious cognition. Cognition, 88(2), 223–242.

Lo, C. C., Boucher, L., Pare, M., Schall, J. D., & Wang, X. J. (2009). Proactive Inhibitory Control and Attractor Dynamics in Countermanding Action: A Spiking Neural Circuit Model. Journal of Neuroscience, 29(28), 9059–9071. doi: 10.1523/Jneurosci.6164-08.2009

Logan, G. D., & Burkell, J. (1986). Dependence and Independence in Responding to Double Stimulation: A Comparison of Stop, Change, and Dual-Task Paradigms. Journal of Experimental Psychology: Human Percection and Performance, 12(4), 549–563.

Logan, G. D., & Cowan, W. B. (1984). On the Ability to Inhibit Thought and Action - a Theory of an Act of Control. Psychological Review, 91(3), 295–327.

Logan, G. D., Yamaguchi, M., Schall, J. D., & Palmeri, T. J. (2015). Inhibitory control in mind and brain 2.0: blocked-input models of saccadic countermanding. Psychol Rev, 122(2), 115–147. doi: 10.1037/a0038893

Matzke, D., Love, J., & Heathcote, A. (2017). A Bayesian approach for estimating the probability of trigger failures in the stop-signal paradigm. Behav Res Methods, 49(1), 267–281. doi: 10.3758/s13428-015-0695-8

McBride, J., Boy, F., Husain, M., & Sumner, P. (2012). Automatic motor activation in the executive control of action. Frontiers in Human Neuroscience, 6(April), Article 82. doi: 10.3389/fnhum.2012.00082

McIntosh, R. D., & Buonocore, A. (2014). Saccadic inhibition can cause the remote distractor effect, but the remote distractor effect may not be a useful concept. J Vis, 14(5), 15. doi: 10.1167/14.5.15

Meeter, M., Van der Stigchel, S., & Theeuwes, J. (2010). A competitive integration model of exogenous and endogenous eye movements. Biological Cybernetics, 102(4), 271–291.

Monsell, S., & Driver, J. (2000). Banishing the control homunculus. In S. Monsell & J. Driver (Eds.), Control of cognitive processes: Attention and performance XVIII (pp. 3–32). Cambridge, MA: MIT Press.

Morein-Zamir, S., & Kingstone, A. (2006). Fixation offset and stop signal intensity effects on saccadic countermanding: a crossmodal investigation. Experimental Brain Research, 175(3), 453–462. doi: 10.1007/s00221-006-0564-x

Munoz, D. P., & Wurtz, R. H. (1993a). Fixation cells in monkey superior colliculus. I. Characteristics of cell discharge. J Neurophysiol, 70(2), 559–575.

Munoz, D. P., & Wurtz, R. H. (1993b). Fixation cells in monkey superior colliculus. II. Reversible activation and deactivation. J Neurophysiol, 70(2), 576–589.

Noorani, I., & Carpenter, R. H. (2017). Not moving: the fundamental but neglected motor function. Philos Trans R Soc Lond B Biol Sci, 372(1718). doi: 10.1098/rstb.2016.0190

O’Connor, D. H., Fukui, M. M., Pinsk, M. A., & Kastner, S. (2002). Attention modulates responses in the human lateral geniculate nucleus. Nat Neurosci, 5(11), 1203–1209. doi: 10.1038/nn957

Ozyurt, J., Colonius, H., & Arndt, P. A. (2003). Countermanding saccades: evidence against independent processing of go and stop signals. Percept Psychophys, 65(3), 420–428.

Paré, M., & Hanes, D. P. (2003). Controlled Movement Processing: Superior Colliculus Activity Associated with Countermanded Saccades. The Journal of neuroscience: the official journal of the Society for Neuroscience, 23(16), 6480–6489.

Pouget, P., Logan, G. D., Palmeri, T. J., Boucher, L., Pare, M., & Schall, J. D. (2011). Neural Basis of Adaptive Response Time Adjustment during Saccade Countermanding. Journal of Neuroscience. doi: 10.1523/JNEUROSCI.1868-11.2011

Purcell, B. A., Heitz, R. P., Cohen, J. Y., Schall, J. D., Logan, G. D., & Palmeri, T. J. (2010). Neurally Constrained Modeling of Perceptual Decision Making. Psychological Review, 117(4), 1113–1143. doi: Doi 10.1037/A0020311

Ramakrishnan, A., Sureshbabu, R., & Murthy, A. (2012). Understanding How the Brain Changes Its Mind: Microstimulation in the Macaque Frontal Eye Field Reveals How Saccade Plans Are Changed. Journal of Neuroscience, 32(13), 4457–4472. doi: 10.1523/Jneurosci.3668-11.2012

Ratcliff, R., & McKoon, G. (2008). The diffusion decision model: theory and data for two-choice decision tasks. Neural Comput, 20(4), 873–922. doi: 10.1162/neco.2008.12-06-420

Ray, S., Pouget, P., & Schall, J. D. (2009). Functional Distinction Between Visuomovement and Movement Neurons in Macaque Frontal Eye Field During Saccade Countermanding. Journal of Neurophysiology, 102(6), 3091–3100. doi: 10.1152/Jn.00270.2009

Reingold, E. M., & Stampe, D. M. (2002). Saccadic inhibition in voluntary and reflexive saccades. J Cogn Neurosci, 14(3), 371–388.

Reingold, E. M., & Stampe, D. M. (2004). Saccadic inhibition in reading. Journal of Experimental Psychology-Human Perception and Performance, 30(1), 194–211. doi: 10.1037/0096-1523.30.1.194

Reppert, T. R., Servant, M., Heitz, R. P., & Schall, J. D. (2018). Neural mechanisms of speed-accuracy tradeoff of visual search: saccade vigor, the origin of targeting errors, and comparison of the superior colliculus and frontal eye field. J Neurophysiol, 120(1), 372–384. doi: 10.1152/jn.00887.2017

Rolfs, M., Kliegl, R., & Engbert, R. (2008). Toward a model of microsaccade generation: the case of microsaccadic inhibition. J Vis, 8(11), 5 1-23. doi: 10.1167/8.11.5

Salinas, E., & Stanford, T. R. (2013). The countermanding task revisited: fast stimulus detection is a key determinant of psychophysical performance. The Journal of neuroscience: the official journal of the Society for Neuroscience, 33(13), 5668–5685. doi: 10.1523/JNEUROSCI.3977-12.2013

Salinas, E., & Stanford, T. R. (2018). Saccadic inhibition interrupts ongoing oculomotor activity to enable the rapid deployment of alternate movement plans. Sci Rep, 8(1), 14163. doi: 10.1038/s41598-018-32224-5

Schall, J. D., Palmeri, T. J., & Logan, G. D. (2017). Models of inhibitory control. Philos Trans R Soc Lond B Biol Sci, 372(1718). doi: 10.1098/rstb.2016.0193

Schall, J. D., & Thompson, K. G. (1999). Neural selection and control of visually guided eye movements. Annual Review of Neuroscience, 22, 241–259.

Schmidt, R., & Berke, J. D. (2017). A Pause-then-Cancel model of stopping: evidence from basal ganglia neurophysiology. Philos Trans R Soc Lond B Biol Sci, 372(1718). doi: 10.1098/rstb.2016.0202

Schmolesky, M. T., Wang, Y. C., Hanes, D. P., Thompson, K. G., Leutgeb, S., Schall, J. D., & Leventhal, A. G. (1998). Signal timing across the macaque visual system. Journal of Neurophysiology, 79(6), 3272–3278.

Shadlen, M. N., Britten, K. H., Newsome, W. T., & Movshon, J. A. (1996). A computational analysis of the relationship between neuronal and behavioral responses to visual motion. Journal of Neuroscience, 16(4), 1486–1510.

Skippen, P., Matzke, D., Heathcote, A., Fulham, W. R., Michie, P., & Karayanidis, F. (2019). Reliability of triggering inhibitory process is a better predictor of impulsivity than SSRT. Acta Psychologica, 192, 104–117. doi: 10.1016/j.actpsy.2018.10.016

Stevenson, S. A., Elsley, J. K., & Corneil, B. D. (2009). A “Gap Effect” on Stop Signal Reaction Times in a Human Saccadic Countermanding Task. Journal of Neurophysiology, 101(2), 580–590. doi: 10.1152/jn.90891.2008

Stuphorn, V., Taylor, T. L., & Schall, J. D. (2000). Performance monitoring by the supplementary eye field. Nature, 408(6814), 857–860.

Sumner, P., & Husain, M. (2008). At the edge of consciousness: automatic motor activation and voluntary control. The Neuroscientist, 14(5), 474–486.

Trappenberg, T. P., Dorris, M. C., Munoz, D. P., & Klein, R. M. (2001). A model of saccade initiation based on the competitive integration of exogenous and endogenous signals in the superior colliculus. Journal of Cognitive Neuroscience, 13(2), 256–271.

Usher, M., & McClelland, J. L. (2001). The time course of perceptual choice: The leaky, competing accumulator model. Psychological Review, 108(3), 550–592.

van Gaal, S., Lamme, V. A., Fahrenfort, J. J., & Ridderinkhof, K. R. (2011). Dissociable brain mechanisms underlying the conscious and unconscious control of behavior. J Cogn Neurosci, 23(1), 91–105. doi: 10.1162/jocn.2010.21431

Verbruggen, F., Best, M., Bowditch, W. A., Stevens, T., & McLaren, I. P. (2014). The inhibitory control reflex. Neuropsychologia, 65, 263–278. doi: 10.1016/j.neuropsychologia.2014.08.014

Verbruggen, F., Chambers, C. D., & Logan, G. D. (2013). Fictitious inhibitory differences: how skewness and slowing distort the estimation of stopping latencies. Psychol Sci, 24(3), 352–362. doi: 10.1177/0956797612457390

Verbruggen, F., & Logan, G. D. (2008). Automatic and Controlled Response Inhibition: Associative Learning in the Go/No-Go and Stop-Signal Paradigms. Journal of Experimental Psychology-General, 137(4), 649–672.

Verbruggen, F., & Logan, G. D. (2009). Proactive Adjustments of Response Strategies in the Stop-Signal Paradigm. Journal of Experimental Psychology-Human Perception and Performance, 35(3), 835–854.

Verbruggen, F., McLaren, I. P., & Chambers, C. D. (2014). Banishing the Control Homunculi in Studies of Action Control and Behavior Change. Perspect Psychol Sci, 9(5), 497–524. doi: 10.1177/1745691614526414

Verbruggen, F., Stevens, T., & Chambers, C. D. (2014). Proactive and reactive stopping when distracted: an attentional account. J Exp Psychol Hum Percept Perform, 40(4), 1295–1300. doi: 10.1037/a0036542

Walker, R., Mannan, S., Maurer, D., Pambakian, A. L. M., & Kennard, C. (2000). The oculomotor distractor effect in normal and hemianopic vision. Proc. R. SOc. Lond. B, 267, 431–438.

Wessel, J. R., & Aron, A. R. (2017). On the Globality of Motor Suppression: Unexpected Events and Their Influence on Behavior and Cognition. Neuron, 93(2), 259–280. doi: 10.1016/j.neuron.2016.12.013

Wiecki, T. V., & Frank, M. J. (2013). A computational model of inhibitory control in frontal cortex and basal ganglia. Psychol Rev, 120(2), 329–355. doi: 10.1037/a0031542

Wong-Lin, K., Eckhoff, P., Holmes, P., & Cohen, J. D. (2010). Optimal performance in a countermanding saccade task. Brain Research, 1318, 178–187. doi: Doi 10.1016/J.Brainres.2009.12.018

Xu, K. Z., Anderson, B. A., Emeric, E. E., Sali, A. W., Stuphorn, V., Yantis, S., & Courtney, S. M. (2017). Neural Basis of Cognitive Control over Movement Inhibition: Human fMRI and Primate Electrophysiology Evidence. Neuron, 96(6), 1447. doi: 10.1016/j.neuron.2017.11.010

